# Optimised DNA extraction and library preparation for minute arthropods: application to target enrichment in chalcid wasps used for biocontrol

**DOI:** 10.1101/437590

**Authors:** Astrid Cruaud, Sabine Nidelet, Pierre Arnal, Audrey Weber, Lucian Fusu, Alex Gumovsky, John Huber, Andrew Polaszek, Jean-Yves Rasplus

## Abstract

Enriching subsets of the genome prior to sequencing allows focusing effort on regions that are relevant to answer specific questions. As experimental design can be adapted to sequence many samples simultaneously, using such approach also contributes to reduce cost. In the field of ecology and evolution, target enrichment is increasingly used for genotyping of plant and animal species or to better understand the evolutionary history of important lineages through the inference of statistically robust phylogenies. Limitations to routine target enrichment by research laboratories are both the complexity of current protocols and low input DNA quantity. Thus, working with tiny organisms such as micro-arthropods can be challenging. Here, we propose easy to set up optimisations for DNA extraction and library preparation prior to target enrichment. Prepared libraries were used to capture 1432 Ultra-Conserved Elements (UCEs) from microhymenoptera (Chalcidoidea), which are among the tiniest insects on Earth and the most commercialized worldwide for biological control purposes. Results show no correlation between input DNA quantities (1.8-250ng, 0.4 ng with an extra whole genome amplification step) and the number of sequenced UCEs. Phylogenetic inferences highlight the potential of UCEs to solve relationships within the families of chalcid wasps, which has not been achieved so far. The protocol (library preparation + target enrichment), allows processing 96 specimens in five working days, by a single person, without requiring the use of expensive robotic molecular biology platforms, which could help to generalize the use of target enrichment for minute specimens.

## INTRODUCTION

Enriching subsets of the genome prior to sequencing (target enrichment, Mamanova *et al.* 2010) allows effort to be concentrated on genomic regions that are relevant to answer specific research questions. Using this approach also contributes to reducing cost, as experimental design can be adapted to sequence many samples simultaneously. In the fields of ecology and evolution, target enrichment has been used for genotyping or phylogenomics of plant and animal species (Gasc *et al.* 2016; Lemmon & Lemmon 2013), to characterize phenotypic traits (e.g., Muraya *et al.* 2015) or to explore microbial ecosystems (Gasc & Peyret 2018).

However, routine target enrichment by research laboratories is limited both by the complexity of current protocols, and by input DNA quantity that may be very low for some minute species (e.g. micro-arthropods < 1mm) and /or old /rare (museum) specimens. Indeed, current protocols (DNA extraction, library preparation, target-enrichment) are time consuming and require handling expertise. They have been initially developed to work with large amounts of input DNA (e.g., vertebrates or large/medium size insects; Faircloth *et al.* 2015; McCormack *et al.* 2013) and include many purification steps that increase DNA loss. Working on hyper-diverse groups of microarthropods is challenging, as it requires one to perform the extraction on i) a large number of specimens/species to be representative of the overall diversity of the group, without the possibility of using pipetting robots that increase DNA loss, ii) single individuals because species complexes are frequent (Al Khatib *et al.* 2014; Kenyon *et al.* 2015; Mottern & Heraty 2014), iii) the whole insect without destruction for vouchering and often prior to species identification, iv) rare species that have been collected once and may be represented in collection by a few specimens or only one specimen and, sometimes, v) old and dry museum specimens used for species description (types).

In this study, we propose optimised protocols for DNA extraction and library preparation for target enrichment purposes, as well as a custom pipeline to analyse the sequence data obtained. We used these protocols and customised pipeline to capture and analyse Ultra Conserved Elements (UCEs) in minute wasps, the chalcids (Insecta: Hymenoptera: Chalcidoidea, Heraty *et al.* 2013; Noyes 2018), that are key components of terrestrial ecosystems. Chalcids are key models for basic and applied research. With an estimated diversity of more than 500,000 species these microhymenoptera have colonised almost all extant terrestrial habitats. Many of them develop as parasitoids of arthropod eggs, larvae or pupae. As such, they are both key regulators of the populations of many other arthropod species in natural ecosystems and are increasingly used worldwide as biocontrol agents (e.g., Consoli *et al.* 2010; Heraty 2009). A few of them, especially *Nasonia* (Pteromalidae) or *Trichogramma* (Trichogrammatidae) species are also used as model systems to answer challenging questions about sex determination, genetics of speciation, host-symbiont interactions or behavioural ecology (e.g., Pinto *et al.* 1991; Stouthamer *et al.* 1990; Werren & Loehlin 2009). Chalcidoidea has undergone a spectacular radiation resulting in a huge diversity of morphologies and sizes (Gibson *et al.* 1999; Heraty *et al.* 2013), but are generally small insects (< 2 mm long). Among them, *Kikiki huna* Huber (Mymaridae) at 158 μm long is the smallest winged insect currently known and the wingless male of *Dicopomorpha echmepterygis* Mockford at 139 μm is the smallest insect currently known (Huber & Noyes 2013). Notably, most species used for biological control, belonging mainly to five families (Aphelinidae, Encyrtidae, Eulophidae, Mymaridae and Trichogrammatidae) are among the tiniest wasps on earth (< 1mm).

Their small size, huge diversity and widespread morphological convergence make chalcid wasps difficult to identify to species by non-expert taxonomists, which limits their use in biological control. Attempts have been made to resolve the phylogeny of the whole superfamily (Heraty *et al.* 2013; Munro *et al.* 2011) or a few families (Burks *et al.* 2011; Chen *et al.* 2004; Cruaud *et al.* 2012; Desjardins *et al.* 2007; Janšta *et al.* 2017; Owen *et al.* 2007) but none has succeeded. Indeed, the few markers that could be targeted with Sanger sequencing were not informative enough to solve deeper relationships. A study based on transcriptome data (3,239 single-copy genes) obtained from 37 species of chalcids and 11 outgroups also failed to solve relationships within the superfamily (Peters *et al.* 2018). As only a representative sampling in both markers and taxa will allow one to draw accurate conclusions on the history of this hyperdiverse group, target enrichment approaches appear relevant. More specifically, targeting UCEs and their flanking regions that have been proven useful to solve ancient and recent divergences (Faircloth *et al.* 2012; Smith *et al.* 2014) seems pertinent. Indeed, a set of probes has been developed to target UCEs in Hymenoptera (Faircloth *et al.* 2015). This set and an enriched one (Branstetter *et al.* 2017c) were successfully used to solve the phylogeny of a few groups of ants, wasps and bees for which the amount of DNA was not limiting (Blaimer *et al.* 2015; Blaimer *et al.* 2016a; Blaimer *et al.* 2016b; Bossert *et al.* 2017; Branstetter *et al.* 2017a; Branstetter *et al.* 2017b; Jesovnik *et al.* 2017; Prebus 2017; Ward & Branstetter 2017). Thus, contributing to the global effort to solve the Hymenoptera tree of life while addressing the challenge of the phylogeny of chalcid wasps seemed sound.

Here, we provide a detailed description of the optimised protocol for DNA extraction and library preparation, followed by a description of the phylogenetic trees obtained through target enrichment of UCEs from 96 species belonging to seven families and one subfamily of chalcids used for biological control (Aphelinidae, Azotidae, Encyrtidae, Eulophidae, Mymaridae, Pteromalidae: Eunotinae, Signiphoridae, Trichogrammatidae) as well as three outgroups in Mymmaromatidae, the putative sister group to Chalcidoidea (Gibson *et al.* 2007; Heraty *et al.* 2013).

## MATERIALS AND METHODS

### Sampling

Samples were taken from the personal collections of the co-authors of this paper, or borrowed from the Queensland Museum (Australia) or the Australian National Insect Collection, Canberra. Details of the samples included in the analysis are presented in Table S1. Most specimens sampled in the field were placed directly into ethanol for storage. On average, specimens spent 3.5 years in ethanol before being processed (maximum storage time in alcohol = 34 years). Two specimens were critical point dried 25 or 34 years ago. UCE data for three specimens were retrieved from a previous study (Branstetter *et al.* 2017a): *Euplectrus* sp. (empirical data); *Copidosoma floridanum* and *Trichogramma pretiosum* (*in silico* extraction of UCEs from genomes).

### DNA extraction

DNA was extracted using the Qiagen DNeasy Blood and Tissue kit. All extractions were conducted without destruction of the specimens’ external (and certain internal) structures, with digestion and lysis of just the soft tissues. In this way, actual or potential type specimens are preserved. An often-essential feedback to the morphology is also preserved which is critical in this difficult group. The following modifications were made to manufacturer’s protocol. Samples were incubated overnight in an Eppendorf thermomixer (temperature = 56°C, mixing frequency = 300 rpm). To increase DNA yield, two successive elutions (50 μL each) were performed with heated buffer AE (56°C) and an incubation step of 15 minutes followed by centrifugation (8000 rpm for 1 minutes at room temperature). Eppendorf microtubes LoBind 1,5ml were used for elution and to store DNA at −20°C until library preparation. DNA was quantified with a Qubit® 2.0 Fluorometer (Invitrogen). The final version of the DNA extraction protocol is available as an additional file 1. Vouchers were deposited at CBGP, Montferrier-sur-Lez, France or returned to their owner.

### Whole genome amplification

DNA extracted from two specimens was subjected to ethanol precipitation and whole genome amplification (WGA) using the GenomiPhi™ V2 DNA Amplification kit (GE Healthcare) as described in Cruaud *et al.* (in press). 1μl of concentrated DNA was used as input (i.e. 4 ng or 0.4 ng, Table S1).

### Library preparation

Our starting point was the protocol described in http://ultraconserved.org and Faircloth *et al*. (2015). The final goal was to obtain a standardized protocol that could be implemented by one person and which, in 5 working days, would made possible the manual preparation of libraries and the capture of UCEs from 96 samples in parallel. Steps by steps optimisations were made specially to remove time-consuming purification steps and different reagents were tested. The final version of the protocol is available as additional file 2. Briefly, input DNA was sheared to a size of ca 400 bp using the Bioruptor® Pico (Diagenode). End repair, 3’-end adenylation, adapters ligation and PCR enrichment were then performed with the NEBNext Ultra II DNA Library prep kit for Illumina (NEB). We used barcoded adapters that contained amplification and Illumina sequencing primer sites, as well as a nucleotide barcode of 5 or 6 bp long for sample identification (additional file 3). Pools of 16 samples were made at equimolar ratio. We enriched each pool using the 2749 probes designed by Faircloth *et al.* (2015) using MYbaits kits (MYcroarray, Inc.). We followed manufacturer’s protocol (MYbaits, user manual version 3, http://www.mycroarray.com/pdf/MYbaits-manual-v3.pdf). The hybridization reaction was run for 24h at 65°C. Post enrichment amplification was performed on beads with the KAPA Hifi HotStart ReadyMix. The enriched libraries were quantified with Qubit, an Agilent Bioanalizer and qPCR with the Library Quantification Kit-Illumina/Universal from KAPA (KK4824). They were then pooled at equimolar ratio. Paired-end sequencing (2*300bp) was performed on an Illumina Miseq platform at UMR AGAP (Montpellier, France) to get longer flanking regions and, as a consequence, more information to differentiate closely related species.

### Raw data cleaning

The analytical workflow is summarized in figure S1. In the next paragraph, chosen parameter values (different from default value) are provided between parentheses. Quality control checks were performed on raw sequence data with FastQC v.0.11.2 (Andrews 2010). Quality filtering and adapter trimming were performed with Trimmomatic-0.36 (Bolger *et al.* 2014) (LEADING:20 TRAILING:20 SLIDINGWINDOW:4:20 MINLEN:180, with PrefixPE/1= AATGATACGGCGACCACCGAGATCTACACTCTTTCCCTACACGACGCTCTTCCGA TCT and PrefixPE/2 = CAAGCAGAAGACGGCATACGAGATCGGTCTCGGCATTCCTGCTGAACCGCTCTTC CGATCT). Overlapping reads were merged using FLASH-1.2.11 (-M 300) (Magoc & Salzberg 2011). Demultiplexing was performed using a bash custom script (no mismatch in barcode sequences was allowed, additional file 4). Assembly of cleaned reads was performed using CAP3 (-i 25 -o 25 -s 400) (Huang & Madan 1999). The 2749 probes designed by Faircloth *et al.* (2015) were assembled into non-overlapping UCEs (hereafter called reference UCEs, n=1432, additional files 5 and 6) using Geneious 8.1.8 (Kearse *et al.* 2012) and contigs were aligned to this set of reference UCEs using LASTZ Release 1.02.00 (Harris 2007). Contigs that aligned with more than one reference UCE and different contigs that aligned with the same reference UCE were filtered out using Geneious 8.1.8.

### Data analysis

UCEs for which sequences were available for more than 25% of the taxa were kept in the next steps of the analysis. Alignments were performed with MAFFT v7.245 (Katoh & Standley 2013) (-linsi option). Ambiguously aligned blocks were removed using Gblock_0.91b with relaxed constrains (-t=d -b2=b1 -b3=10 -b4=2 -b5=h) (Talavera & Castresana 2007). The final data set was analysed using supermatrix approaches and coalescent-based summary methods. Two gene tree reconciliation approaches were used: ASTRAL-III v5.6.1 (Zhang *et al.* 2018), which computes the phylogeny that agrees with the largest number of quartet trees induced by the set of input gene trees and ASTRID (Vachaspati & Warnow 2015) which takes a set of gene trees, computes a distance matrix (*ca* sum of number of edges in the path between two samples divided by the number of gene trees in which the two samples are represented) and infers a phylogeny from this distance matrix. Following recommendations for incomplete distance matrices, BioNJ was used to compute the phylogeny. Individual trees were inferred from each UCE using raxmlHPC-PTHREADS-AVX (Stamatakis 2014) (version 8.2.4; -f a -x 12345 -p 12345 -# 100 -m GTRGAMMA). ASTRAL and ASTRID analyses were performed with 100 multi-locus bootstrapping (MLBS, site-only resampling (Seo 2008)). Phylogenetic trees were estimated from the concatenate, unpartitioned data set using Maximum Likelihood (ML) approaches as implemented in RAxML and IQTREE v1.6.4 (Nguyen *et al.* 2015). For the RAxML analysis, a rapid bootstrap search (100 replicates) followed by a thorough ML search (-m GTRGAMMA) was performed. For the IQTREE analysis, a ML search with the best-fit substitution model automatically selected was performed with branch supports assessed with ultrafast bootstrap (Minh *et al.* 2013) and SH-aLRT test (Guindon *et al.* 2010) (1000 replicates).

Summary statistics for all data sets (alignment length, number of samples, number of variable sites, number of parsimony informative sites etc.) were calculated using AMAS (Borowiec 2016). Tree annotation was performed with TreeGraph 2.13 (Stöver & Müller 2010). Linear correlation between the number of UCEs and the quantity of DNA used to build the library was tested with the Pearson’s correlation coefficient in R (R Core Team 2015). Analyses were performed on a Dell PowerEdge T630 with 10 Intel Xeon E5-2687 dual-core CPUs (3.1 GHz, 9.60 GT/s), 125 Go RAM and 13 To hard drive and on the Genotoul Cluster (INRA, Toulouse).

## RESULTS

Optimisations made for DNA extraction are detailed in the additional file 1. The final version of the library preparation protocol is available as additional files 2 and 3. Hereafter, the range of values provided between parentheses refers to the range of data that fall between the 2.5th percentile (LB=lower bound) and 97.5th percentile (UP = upper bound). Table S1 contains sequencing information for all samples. The median amount of input DNA was 25ng (LB=1.8ng; UP=250ng). An average of 76,330 reads (cleaned and merged) per sample was obtained (LB=3,359; UP=348,326). The average number of contigs was 3,454 (LB=546; UP=14,012) and the average sequencing depth was 18X (LB=3X; UP=44.0X). The average number of UCEs obtained per sample after filtering of problematic contigs was 687 (LB=193-UP=1,082) with a length comprised between LB=315 and UP=816bp (mean=603bp). Figure 1 and S2 show the variation of the number of UCEs obtained with regard to the amount of input DNA. No significant correlation was observed (Pearson’s correlation coefficient = 0.096, p-value = 0.36).

**Figure 1.**
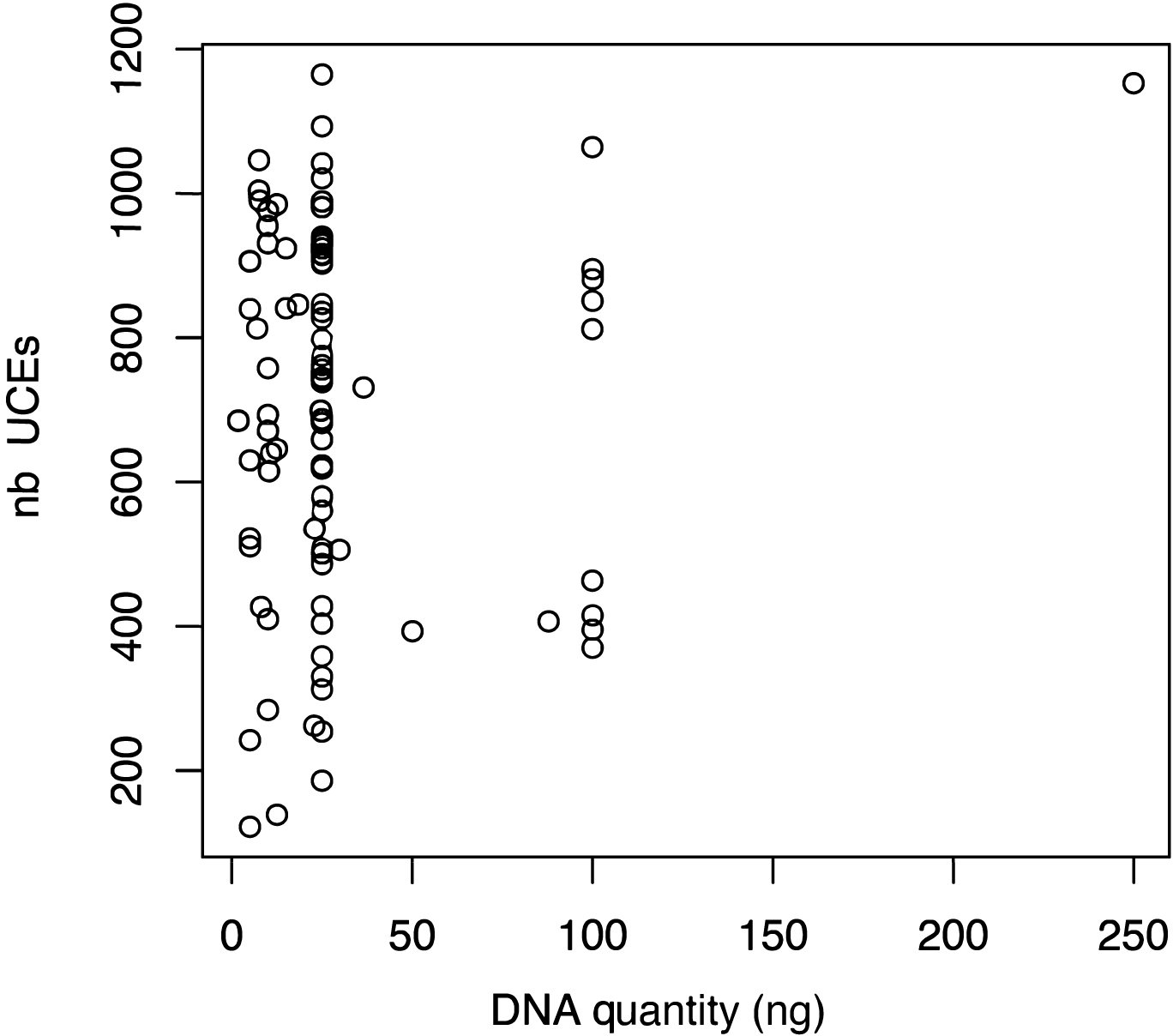
Variation of the number of UCEs with regard to the amount of input DNA (ng)

The final dataset (25%-complete matrix; i.e. at least 25 taxa on the 99 should have a sequence to keep the locus in the analysis) comprised 1,139 UCEs and 340,286bp (missing data = 47.0 %; parsimony informative sites = 72.3 %; GC-content = 42.6%).

Specimens retrieved from a previous study (Branstetter *et al.* 2017a) that were either represented by empirical data (*Euplectrus* sp.) or UCEs extracted from published genomes (*Copidosoma floridanum* and *Trichogramma pretiosum*) displayed a number of UCEs comparable to what was obtained for other specimens. Their placement in the trees (Figures 2; S3-S5) was in accordance with their morphology. Whatever the method used (supermatrix approaches versus coalescent-based summary methods), all families, except for Aphelinidae, were recovered as monophyletic with high support. Aphelinids were split into three groups: 1) a monophyletic Aphelininae + Eretmocerinae; 2) Coccophaginae; 3) *Cales* sp. (Calesinae). The position of *Cales* was ambiguous. *Cales* was either recovered as sister to Trichogrammatidae (RAxML, low support) or as a lineage distinct from all other chalcidoids (all other analyses). Except for Mymaridae that was strongly placed as sister to all other Chalcidoidea in all analyses, the tree backbone remained poorly resolved. Statistical support was much higher within families. In all analyses Azotidae clustered with Signiphoridae, with strong support.

**Figure 2.**
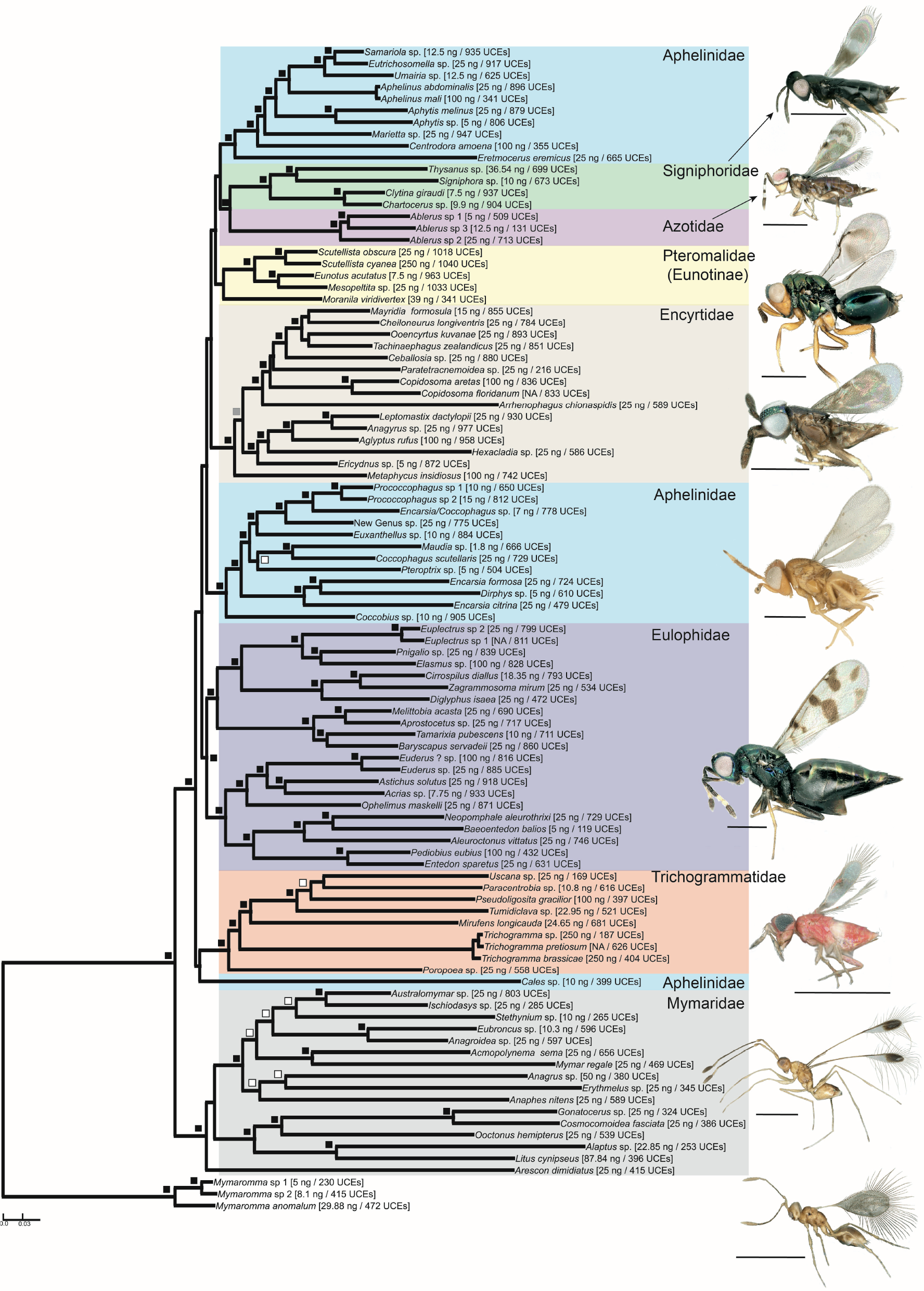
RAxML tree obtained from the analysis of the 25%-complete data set. Black squares indicate node supported with RAxML BP = 100, IQTREE aLRT = 100 / BP = 100 and ASTRAL/ASTRID BP > 75. Grey square indicates node with RAxML BP = 100, IQTREE aLRT = 100 / BP = 100 and ASTRAL BP > 75. White squares indicate nodes with RAxML BP > 95 and IQTREE aLRT > 80 / BP > 95. The DNA quantity used to build the library as well as the number of UCEs analysed for each sample is given in brackets. Photos ©J.-Y. Rasplus. Scale bars = 500 μm. IQTREE, ASTRAL and ASTRID trees are available in figures S3-S5.

## DISCUSSION

To our knowledge this study is the second after Sproul and Maddison (2017) to demonstrate success in library preparation from such low input using commercial kits, and the first to report successful sequencing of >1000 low copy genes in 96 specimens in parallel, from such low input and processing time. Our optimisations differ from what was proposed by Sproul and Maddison (2017). First, we tried to optimize DNA extraction itself by using overnight lysis with gentle mixing to preserve fragile specimens, heated elution buffer and increased incubation time before elution. Second, instead of increasing the number of time-consuming purification steps we decreased them. It is noteworthy that in this library only two historical specimens were included. This may have masked challenges posed by adapter dimers (Burrell *et al.* 2015; Sproul & Maddison 2017; Tin *et al.* 2014) that led Sproul and Maddison (2017) to add a second bead clean-up prior to library amplification. However, we have already used this protocol on hundreds of chalcid and moth species, including historical specimens that were processed the same way as fresh ones and we never had such an issue. Finally, instead of increasing the number of amplification cycles prior to target enrichment, we used a new generation mastermix including a hot start, processive and high-fidelity polymerase (NEBNext Ultra II Q5 Master Mix). Our protocol also allows one to back-up DNA at several steps that allows for multiple attempts without delay in case the first attempt fails. Finally, sequencing was performed on a MiSeq to get longer flanking regions and, as a consequence, more information to differentiate closely related species.

The protocol was successfully used on minute chalcid wasps widely used for biological control purposes. Up to 1165 valid UCEs were captured from 25ng DNA (median amount of DNA used for this study). No correlation was observed between the quantity of DNA used for library preparation and the number of captured UCEs. The average number of captured UCEs validated by our quality control workflow was 687, and 685 valid UCEs were captured from a tiny aphelinid (1.8ng of input DNA). The number of UCEs obtained per individual appears to drop within the basal clades (i.e., Mymaridae and Trichogrammatidae, Figure S2), a result probably linked to the relatively long branches observed in these groups, and that could reduce the efficiency of the probes that were designed from the genome of *Nasonia* (Faircloth *et al.* 2015). Trees were well resolved at the family level, with high statistical support, showing the potential of UCEs to solve long-standing taxonomic issues. However, the tree backbone remained unresolved, a pattern that confirms the rapid diversification of the group (Heraty *et al.* 2013; Peters *et al.* 2018). Understanding the evolutionary history of the group was not the purpose of this paper. Indeed, only a representative sampling in both markers and taxa as well as cutting-edge data analysis will allow drawing accurate conclusions. By providing suitable tools for a fast, easy and affordable acquisition of data, this paper is a first step.

Interestingly, as it has been shown previously on RADseq data (Cruaud *et al.* in press), WGA does not seem to bias the results even when input DNA used for the WGA is below the recommendation on the manufacturer kit (here 0.4 and 4 ng when the Genomiphi kit V2 requires 10 ng). This further increases the possibilities opened by our protocol as input DNA as low as 0.2-0.3 ng may be used when an extra WGA step is included after DNA extraction (Cruaud *et al.* in press). It is noteworthy that reducing the amount of DNA required for library preparation allows one to use extracted DNA for different approaches in parallel (RADseq, amplicon, Shotgun etc.). This also allows one to send DNA back to museums from which specimens were borrowed, for archival purposes. We definitely agree with Sproul and Maddison (2017) who emphasize how important it is not to waste DNA obtained from irreplaceable specimens (whether fresh or historical). It is even more important to capitalize on existing collections as collecting samples for large-scale studies may be more and more difficult, given that many countries have imposed restrictive access regulations, even to academic researchers, to reduce the risk of supposed biopiracy (Divakaran Prathapan *et al.* 2018).

All the elements discussed above indicate that this protocol may be of great help to reconstruct phylogenetic hypotheses in multiple groups of tiny arthropods, e.g., springtails, sandflies, lice, whiteflies, mites, etc. Coupled or not with WGA, some steps of our protocol may also contribute to simplify and hasten construction of other reduced-representation libraries (RAD-Seq, ddRAD-Seq, etc). These methods may then be used to analyse species relationships, hybridization or population genomics (Cruaud *et al.* 2014; Eaton & Ree 2013; Emerson *et al.* 2010; Gagnaire *et al.* 2013; Hipp *et al.* 2014) on minute arthropods or on endangered species using non-invasive DNA sampling (Vila *et al.* 2009).

## ACKNOWLEDGEMENTS

We are grateful to Nicole Fisher (CSIRO, National Museum of Australia, Canberra), Chris Burwell, Christine Lambkin and Susan Wright (Queensland Museum, Brisbane, Australia) for facilitating access to samples and to the Genotoul bioinformatics platform Toulouse Midi-Pyrenees, France for providing computing resources. AC thanks Lisa Schlée, Philippe Cruaud and Maya for their patience in the field and Jean-Pierre Rossi for his help with R. AG thanks V. Fursov (Schmalhausen Institute of Zoology, Ukraine) for his help with specimen identification. This work is part of a large NSF project (Award#1555808) led by John Heraty (UC Riverside USA) that attempts to solve the phylogeny of Chalcidoidea with NGS approaches, and was partly funded by the ANR project TriPTIC (ANR-14-CE18-0002) led by JYR and the SPE department of the INRA (French National Agronomic Institute). LF travels to France were supported by the grants IntegPar (89BM/2017) and NEVPIT (PN-III-P4-ID-PCE-2016-0233) awarded by the Romanian National Authority for Scientific Research, CNCS – UEFISCDI. This paper is dedicated to the memory of our friend Dr John La Salle, whose kindness and sense of humor will be remembered for ever.

## DATA ACCESSIBILITY

Demultiplexed reads are available as a NCBI Sequence Read Archive (ID#XXX)

## AUTHOR CONTRIBUTIONS

Designed the study: AC, JYR; contributed samples and identifications: AC, LF, AG, JH, AP, JYR; optimized protocol: SN with the help of AC, JYR, PA; performed lab work: SN with the help of AC, JYR, LF; sequenced libraries: AW; analysed data: AC, JYR; drafted the manuscript: AC, JYR. All authors commented on the manuscript.

## Supplementary material to

**Table S1:**
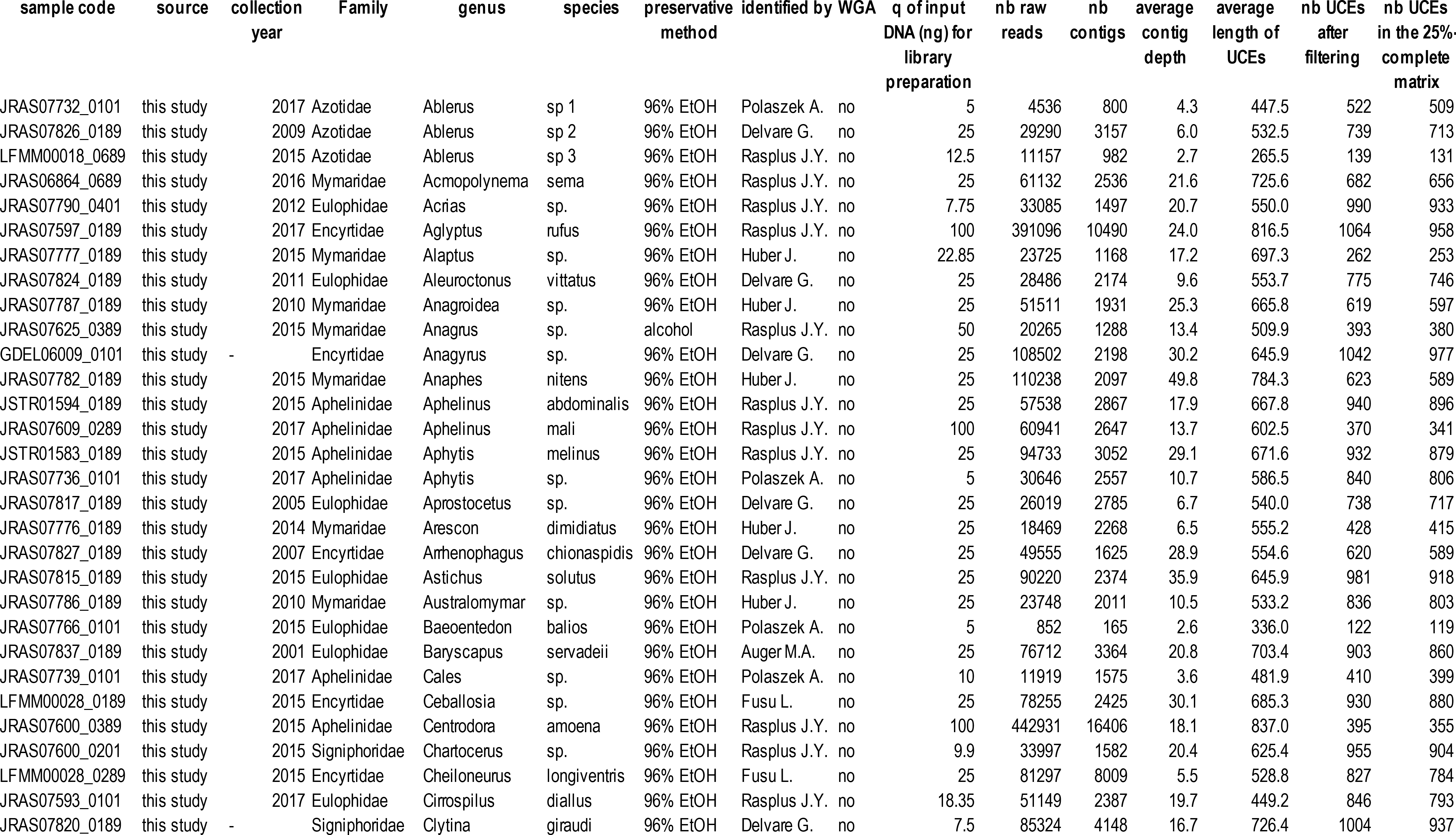
Samples used in this study and results of the UCE-enrichment experiment.

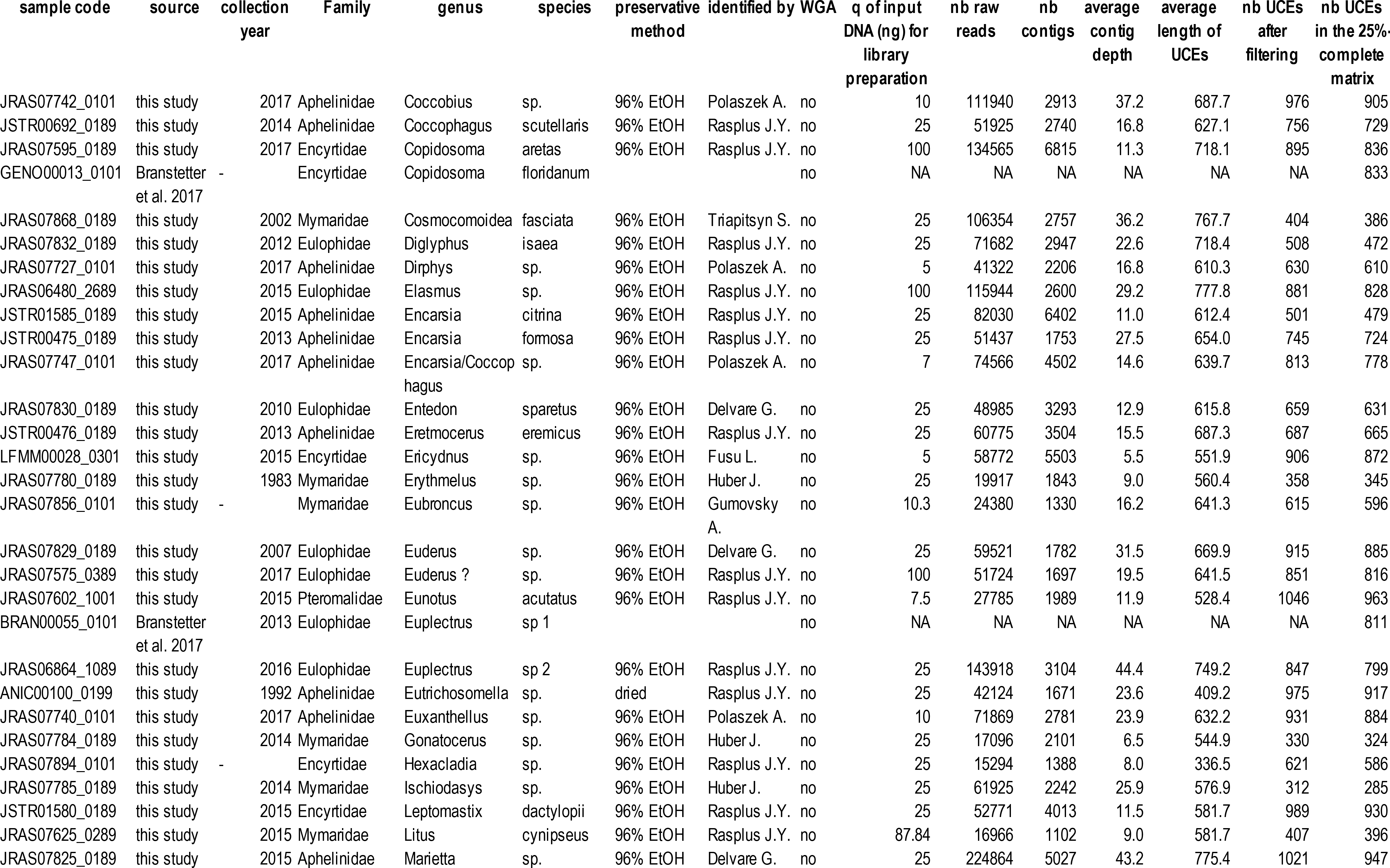

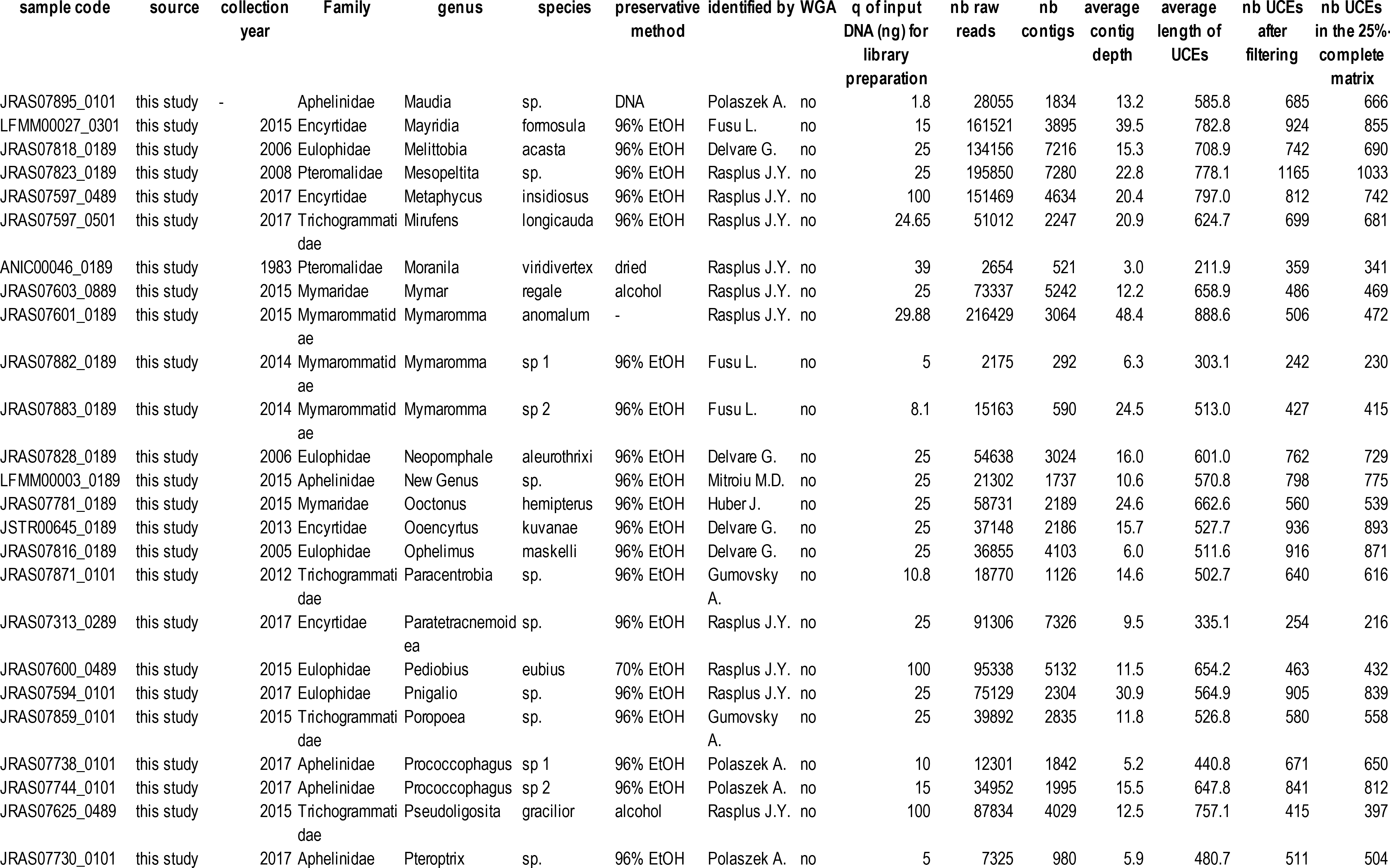

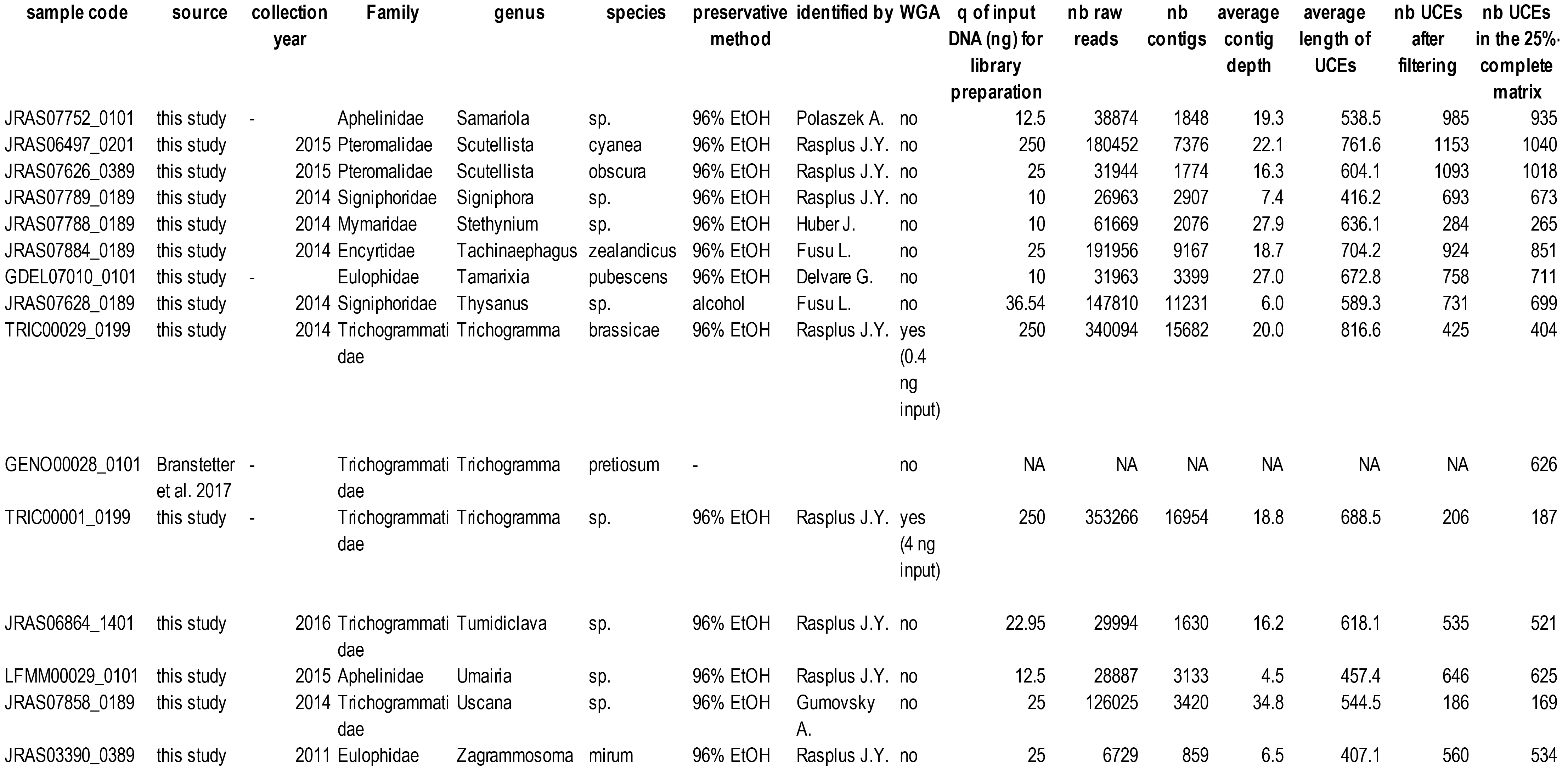

**Figure S1:**
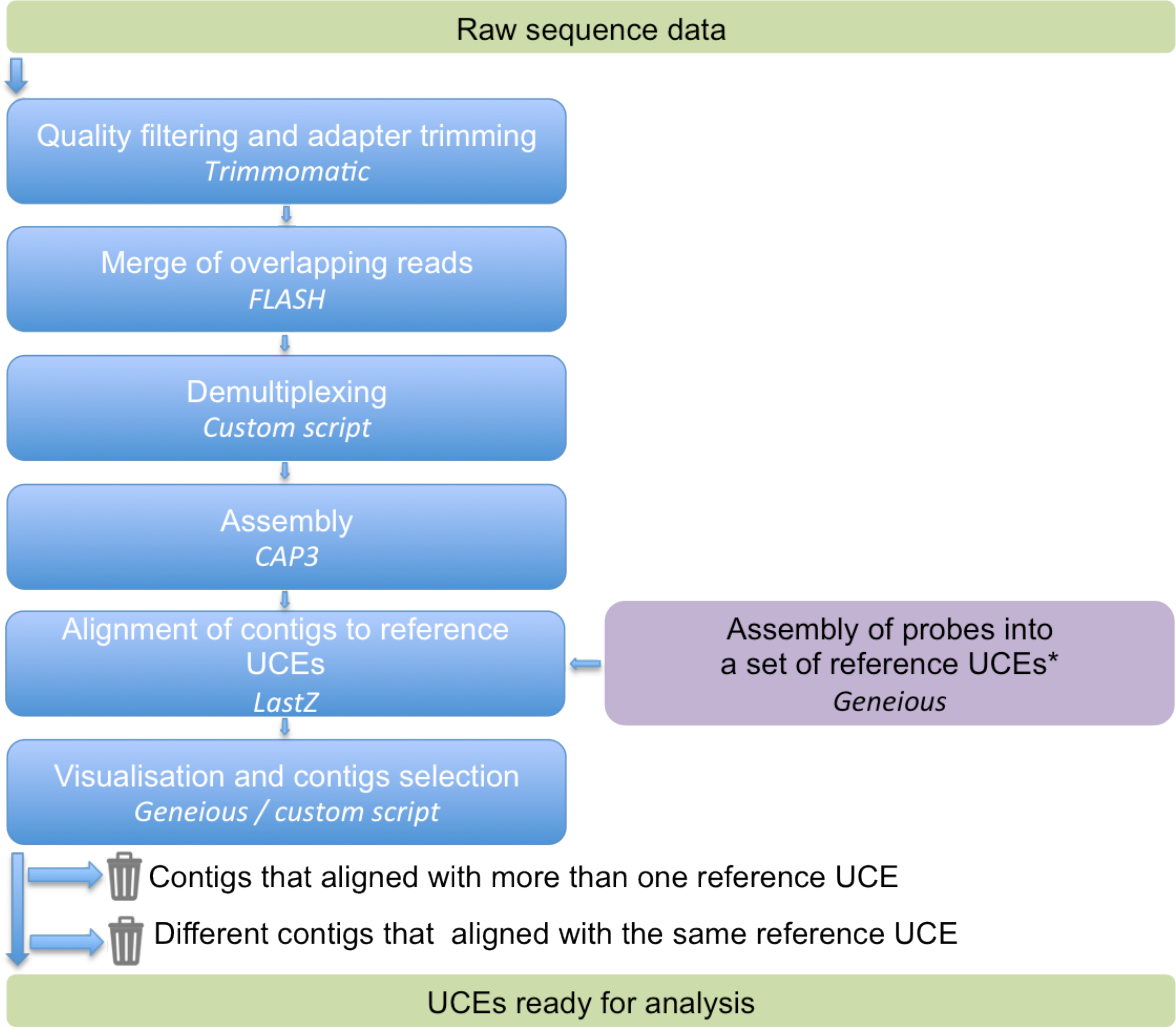
Analytical workflow - Step 1. Pre-processing of the data: from raw reads to raw UCEs. *The 2749 probes designed by Faircloth et al. (2015) were first assembled into a set of non-overlapping loci using Geneious. This set of reference UCE loci (N=1432) was then used to align sample-specific contigs obtained with CAP3. The reference UCEs are available in additional file 4.

**Figure S2:**
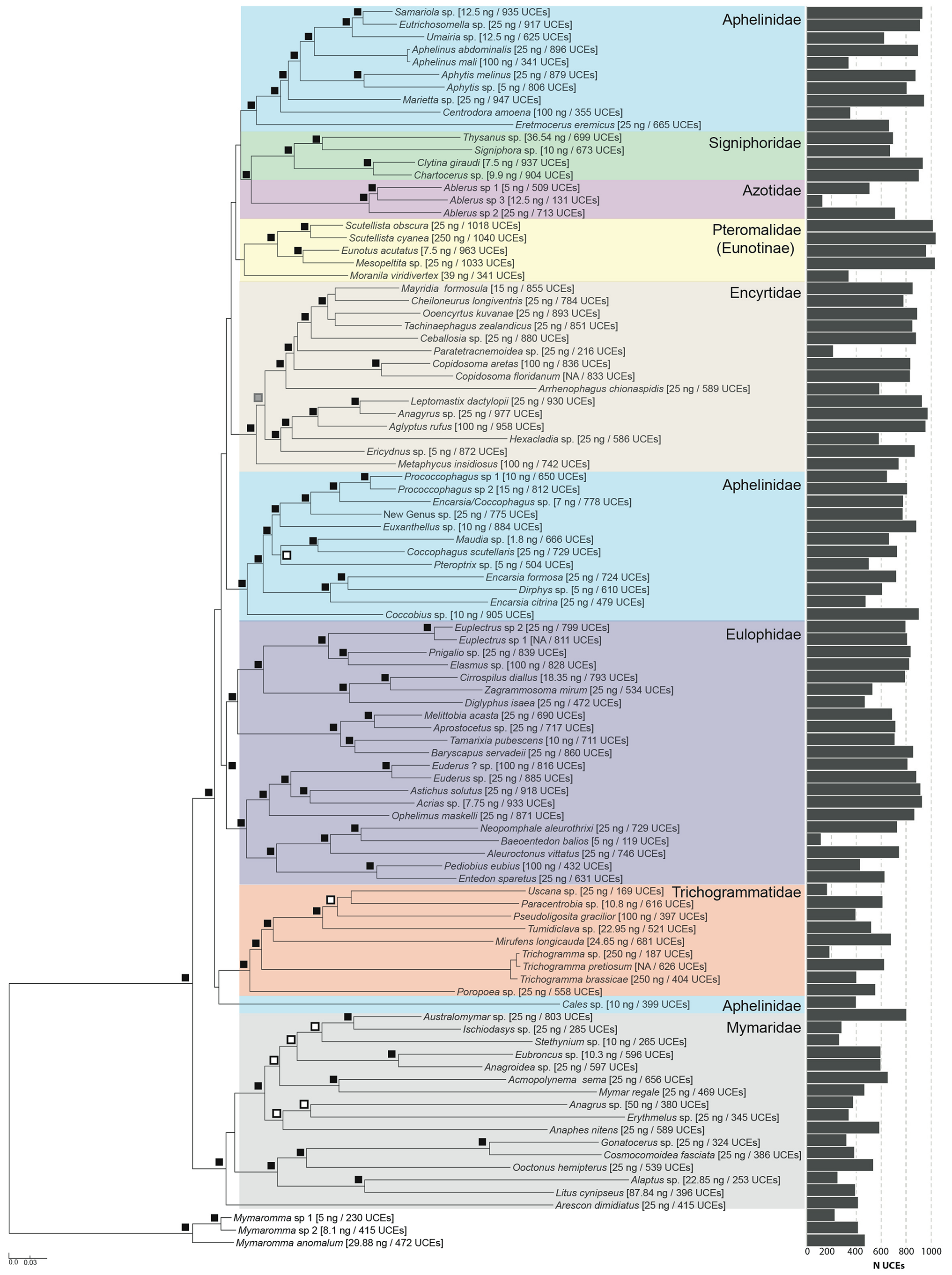
RAxML tree obtained from the analysis of the 25%-complete data set. The histogram on the side of the tree shows the distribution of the number of UCEs analysed for each sample. Black squares indicate node supported with RAxML BP = 100, IQTREE aLRT = 100 / BP = 100 and ASTRAL/ASTRID BP > 75. Grey square indicates node with RAxML BP = 100, IQTREE aLRT = 100 / BP = 100 and ASTRAL BP > 75. White squares indicate nodes with RAxML BP > 95 and IQTREE aLRT > 80 / BP > 95.

**Figure S3:**
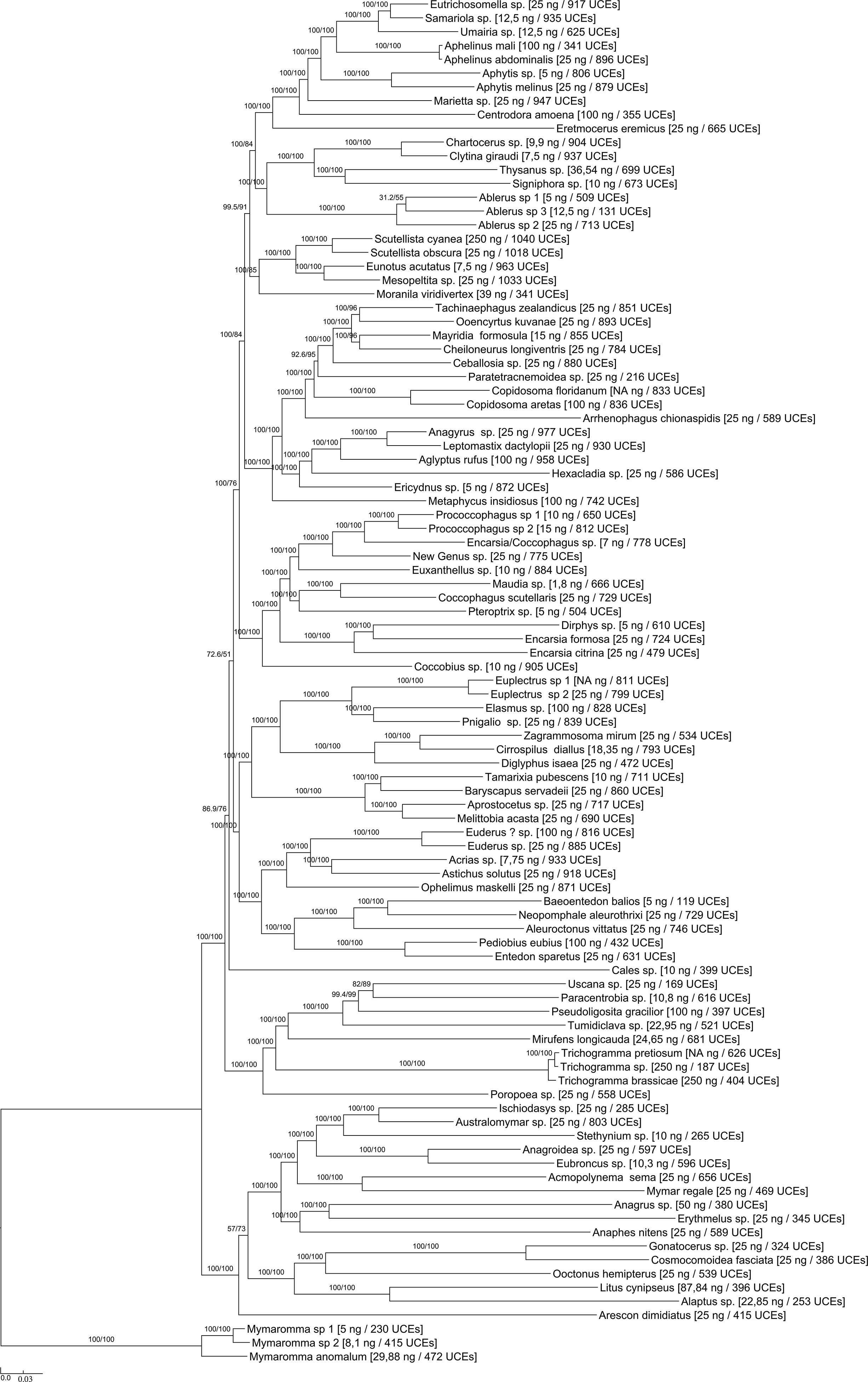
IQTREE tree obtained from the analysis of the 25%-complete data set. SH-aLRT / UFboot values are indicated at nodes. The DNA quantity used to build the library as well as the number of UCEs analysed for each sample is given in brackets.

**Figure S4.**
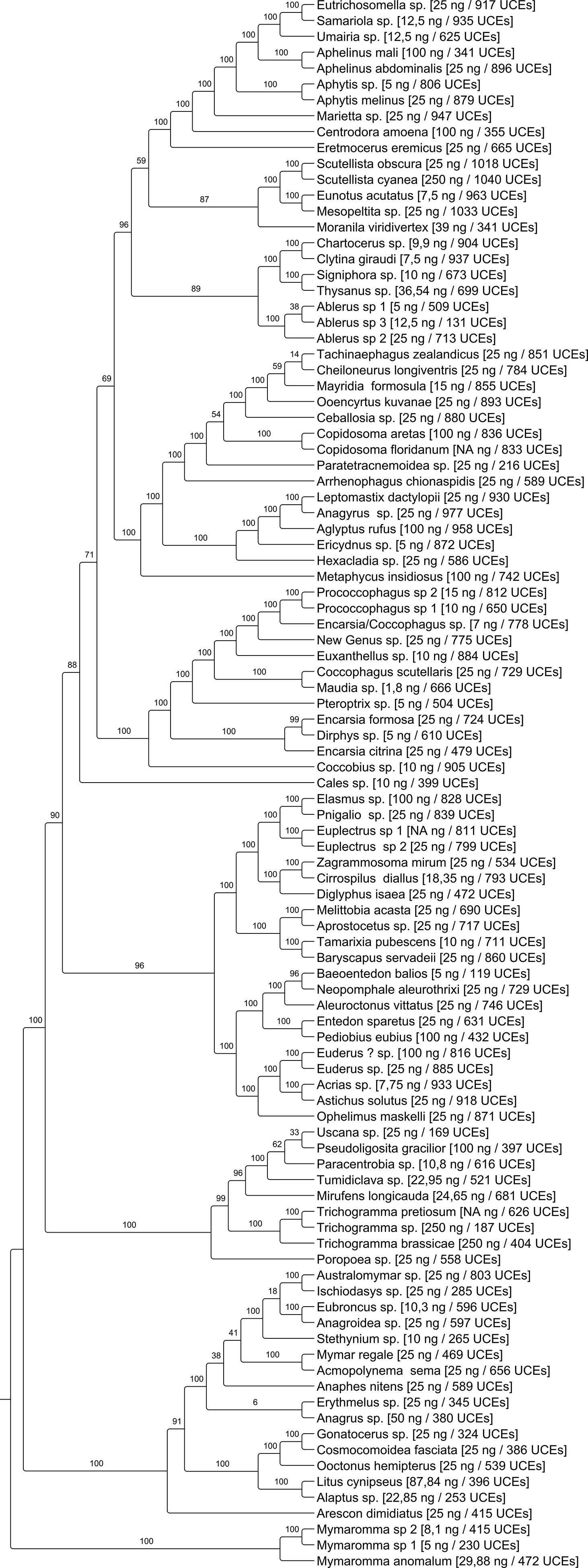
ASTRAL tree obtained from the analysis of the 25%-complete data set. Bootstrap supports (site-only resampling) are indicated at nodes. The DNA quantity used to build the library as well as the number of UCEs analysed for each sample is given in brackets.

**Figure S5.**
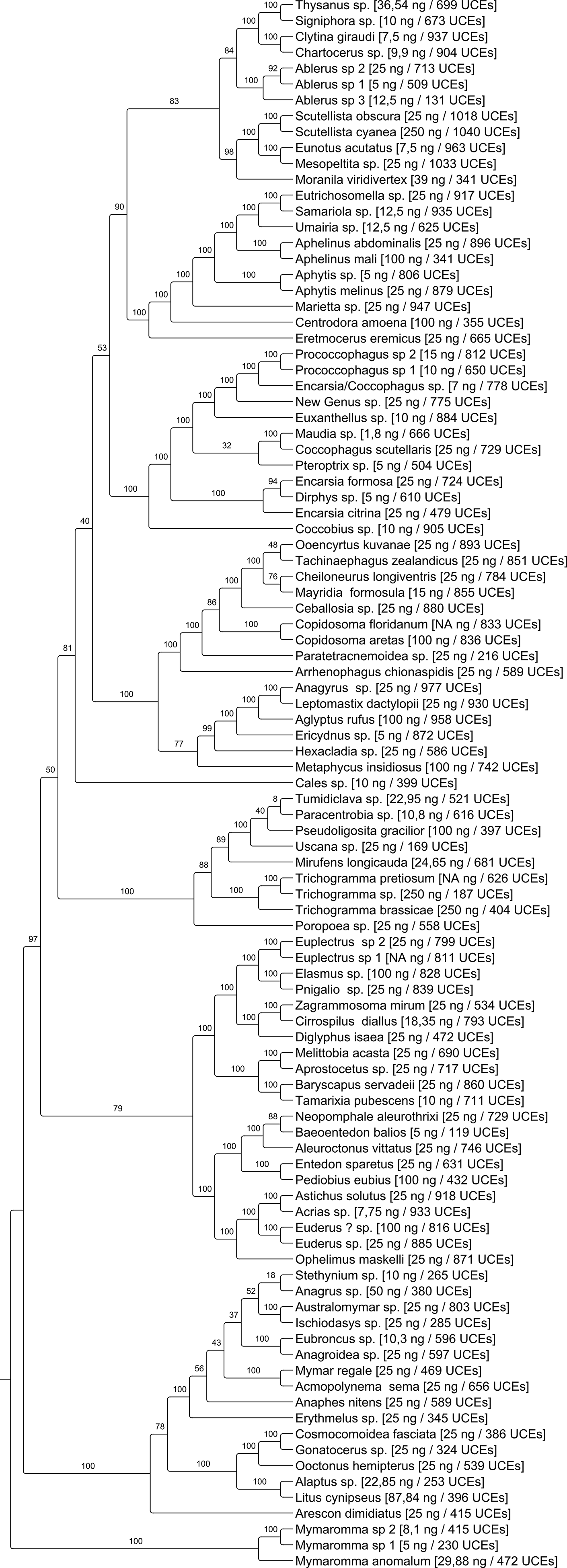
ASTRID tree obtained from the analysis of the 25%-complete data set. Bootstrap supports (site-only resampling) are indicated at nodes. The DNA quantity used to build the library as well as the number of UCEs analysed for each sample is given in brackets.

### Additional file 1: Optimized protocol for DNA extraction

Optimisations were done to the DNeasy Blood & Tissue Kit (250) protocol (Qiagen)

#### Consumables

Tweezers and fine brush/or forceps to manipulate specimens; 1 pot with bleach at a 1:10 dilution with osmosis cleaned water; 1 pot with osmosis cleaned water to wash tweezers/forceps, cup, paper towels, sterile gloves, 50ml Falcon™ tube, 1.5ml Safe-lock tubes (Eppendorf), 1,5ml Microtubes DNA LoBind (Eppendorf).

---------day 1 / afternoon---------

- Label 1.5ml safe-lock tubes with sample codes. Close caps. Put under UV light for 10 min.
- Equilibrate frozen specimens to room temperature before processing.
- Dry specimens on clean paper towel to remove EtOH. Place specimen in tube. Clean forceps to avoid contamination between 2 different specimens: bleach, then water, then dry with paper towel.
- Add 200μl buffer ATL (Qiagen) to each tube. (NB DNeasy® Blood & Tissue Handbook uses 180μl; we use 200μl to improve diffusion of the DNA from the specimen)
- Add 20μl proteinase K to each tube (or prepare a master mix of buffer ATL + proteinase K and add master mix to tubes.)
- Vortex + Pulse spin. (NB if the specimens are fragile, i.e. old and dry mounted collection specimens, dispense the master mix of buffer ATL + proteinase K to each tube and then add specimens; or dispense specimens to tubes and then add the master mix. Do not vortex at all to avoid damage to specimens and place the tubes directly in the thermomixer after ensuring that the specimens are in the liquid.)
- Place tubes in thermomixer (Eppendorf) at 56°C and 300 rpm overnight.

---------day 2 / morning---------

- Remove tubes from thermomixer.
- Vortex + pulse spin. (NB if working with fragile specimens do not vortex. Pulse spin, transfer all the liquid to a new labelled tube leaving the specimen behind and then proceed to the next step.)

(DNeasy® Blood & Tissue Handbook, p. 15, recommends using carrier DNA or RNA for samples containing less than 5 μg of DNA. Though this was not used for this publication, poly (A) RNA (2 μl of 2 μg/μl) is added to the lysate before the next step by one of us (LF) for routine DNA purification from small or old specimens. This should not interfere with library preparation as it was used for example by Sproul & Maddison (2017) as recommended in the QIAamp DNA Micro Kit by Qiagen.)

- Add 200μl buffer AL to each tube (kit Qiagen). (DNeasy® Blood & Tissue Handbook, p. 19, 29 recommends adding RNase A if RNA-free genomic DNA is required. We do not use it because the copurified RNA is not interfering with target enrichment anyway and because RNase A also degrades DNA; see Donà & Houseley (2015).)

A white precipitate may appear that will dissolve during incubation at 70°C.

- Vortex + pulse spin.
- Incubate tubes in thermomixer at 70°C and 300 rpm for 10 minutes.
- During that time, prepare DNeasy mini spin Qiagen columns and 1.5 ml DNA LoBind Eppendorf tubes (label tubes with sample codes). (NB LoBind tubes minimize DNA loss during storage by reducing sample-to-surface binding.)
- Aliquot buffer AE in Eppendorf tubes and heat at 56°C in thermomixer (heated buffer will improve the release of DNA from the resin.)
- Add 200μl absolute ethanol (not provided with the Qiagen kit) to each tube. (NB only high purity ethanol should be used; cheap ethanol may contain traces of other chemicals that will interfere with the solubilisation of the DNA from the membrane in the last step.)
- Vortex + Pulse spin.
- Pipette the liquid (including precipitate) from each tube and transfer into a DNeasy mini spin Qiagen column placed in a 2 ml collection tube (Qiagen kit).
- Centrifuge columns + collection tubes at 6000 × g (8000 rpm) for 1 minute; discard the flow-through and collection tubes. Keep the columns.
- Place spin columns in new collection tubes.
- Add 500μl buffer AW1 (kit Qiagen).
- Centrifuge at 6000 × g (8000rpm) for 1 minute; discard the flow-through and collection tubes. Keep the columns.
- Place spin columns in new collection tubes.
- Add 500μl buffer AW2 (kit Qiagen).
- Centrifuge at 20,000 × g (14,000 rpm) for 3 minutes to dry the columns.
- Rotate columns to 180 degrees in their collection tubes and centrifuge again at 20000 xg (14000rpm) for 3 minutes (this will make sure the column is well dried).
- Place dried spin columns in 1.5ml LoBind tubes that you previously labelled.
- Add 50μl of heated AE buffer. **Deposit buffer right in the middle of the resin.**
- Incubate 15 min at room temperature **(meanwhile, reheat buffer AE.)**
- Centrifuge at 6000 × g (8000 rpm) for 1 minute.
- Rotate columns to 180 degrees and centrifuge again at 6000 × g (8000 rpm) for 1 minute.
- Remove columns + tubes from the centrifuge.
- Add again 50μl of heated buffer **right in the middle of the resin.**
- Incubate 15 min at room temperature.
- Centrifuge at 6000 × g (8000 rpm) for 1 minute at room temperature.
- Rotate columns to 180 degrees and centrifuge again at 6000 × g (8000 rpm) for 1 minute.
- DNA (ca 2 × 50μl) is ready for use.
- Add distilled water to the extracted specimens and incubate at room temperature for 30 min. (NB water is used to eliminate the residual salts left by the buffers. Otherwise they usually crystallise on the specimen upon storage in ethanol.)
- Remove water and replace with ethanol. Keep specimens in the dark in a freezer until mounting. (NB if the specimens are stored at room temperature under ambient light they will become sun-bleached very quickly, much quicker than unextracted specimens.)
- Use a critical point dryer, hexamethyldisilazane, acetone or amyl acetate to dry the specimens before mounting. Do not air dry as they will collapse.

### Additional file 2: Optimized protocol for library preparation

#### 1. DNA quantification

Qubit® 2.0 Fluorometer (Invitrogen); dsDNA HS Assay Kit.

#### 2. DNA normalization and shearing

Bioruptor ® Pico, sonication bath-based rotor 12 samples in parallel.

Transfer 100μl of input DNA in tubes adapted to the Bioruptor ® Pico that will be used for DNA shearing (Diagenode tubes 0.5ml). [now we shear 30ng DNA in routine but you can use as little as 1.8 ng, see main text]

Set up the following program for DNA shearing to obtain a mean fragment size of ca 400pb: 15sec ON / 90sec OFF for 8 cycles.

You can STOP here for up to 1 week (store tubes in a freezer at −20°C).

**The NEBNext Ultra II DNA Library prep kit for Illumina by NEB will be used for End repair, A tailing, Adapter ligation and PCR enrichment of adaptor-ligated DNA.**

#### 3. End repair + A tailing

To keep a backup, only 50 μl of the sheared DNA is used [you may want to re-concentrate DNA on beads in 50 μl when you have a small amount i.e. < 5 ng]

Transfer 50 μl of sheared DNA into a strip / plate.

Thaw NEBNext Ultra II End Prep Reaction Buffer. Vortex + pulse spin.

ATTENTION: Pull out NEBNext Ultra II End Prep Enzyme Mix out of freezer just before use. LIGHT vortex + pulse spin.

|  | Master Mix: |  |
| --- | --- | --- |
|  | x1 reaction | x 96 reactions |
| NEBNext Ultra II End Prep Enzyme Mix | 3 µl | 288 µl |
| NEBNext Ultra II End Prep Reaction Buffer | 7 µl | 672 µl |
| Master mix to add to each tube | 10 µl |  |

Vortex master mix gently + pulse spin.

- Add 10μl of master mix to tubes containing 50μl of sheared DNA.
- Vortex (**mix well.)**
- Pulse spin.
- Place in a thermocycler. <u>**Enable heated lid**</u> and run the following program:

**30 min at 20°C**
**30 min at 65°C**
**Hold at 4°C.**

##### PROCEED WITHOUT DELAY TO ADAPTER LIGATION

###### 4. Adapter ligation and cleanup

See **additional file 3**for adapter sequences and hybridization.

Thaw adapters (1.5μM). Centrifuge to avoid cross contamination. Open carefully.

For 96 samples use the following tagging scheme:

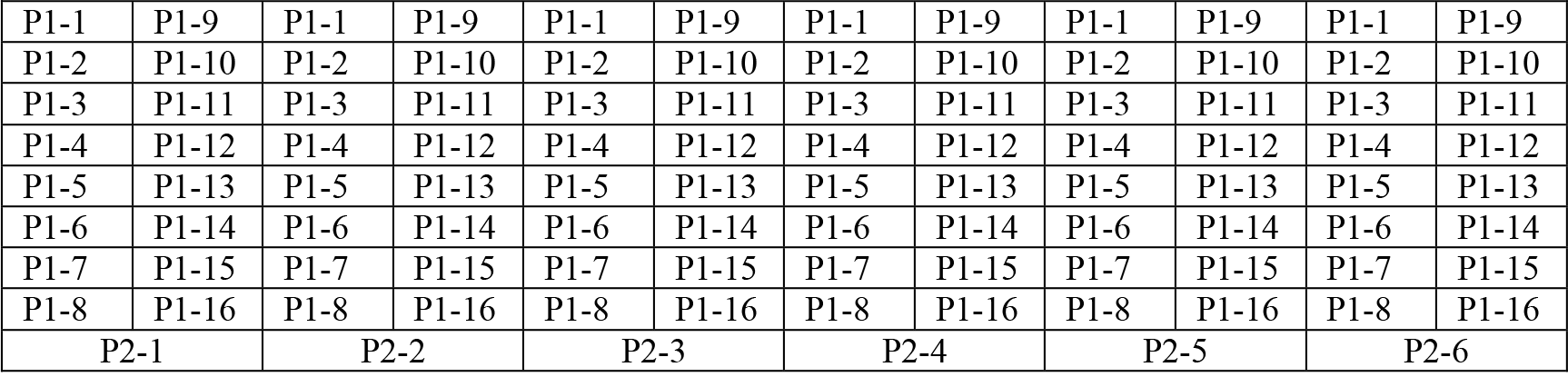

Thaw NEBNext Ultra II Ligation master mix. Light vortex + Pulse spin.

Thaw NEBNext Ligation enhancer. Light Vortex + pulse spin.

Thaw NEBNext Ultra II Ligation master mix. Light vortex + Pulse spin. Thaw NEBNext Ligation enhancer. Light Vortex + pulse spin.
|  | Master Mix: | x1 reaction | x 96 reactions |
| --- | --- | --- | --- |
| NEBNext Ultra II Ligation master mix |  | 30 µl | 2880 µl |
| NEBNext Ligation enhancer |  | 1 µl | 96 µl |
| Master mix to add to each tube |  | 31 µl |  |

###### Vortex (mix well) + pulse spin

- Add 31 μl of master mix to tubes containing the 60 μl end prep reaction mixture obtained in the previous step.
- Add 1.25 μl of P1 adapters (1.5μM) and 1.25 μl of P2 adapters (1.5 μM) to each tube.
- Vortex **(mix well.)**
- Pulse spin.
- Place in a thermocycler. **<u>Disable heated lid</u>** and run the following program:

**15 min at 20°C.** (NB samples can be stored overnight at −20°C.)

###### Clean-up adaptor ligated DNA (remove dimers of adapters)

Purification was conducted with the Agencourt AMPure XP Purification system (Beckman Coulter) and a ALPAQUA magnet plate.

Pull Ampure beads out of fridge. Make sure they are at room temperature (RT) before use.

- Vortex Ampure beads.
- Add 0.8 vol. of Ampure beads for 1 vol. of DNA (i.e. 74.8 μl of beads for 93.5 μl of DNA => vol. tot. = 168.3 μl).
- Vortex + pulse spin.
- Incubate for **5min** at RT.
- Move strip to magnetic rack and wait for **2 to 5min.**
- Remove and discard supernatant without disturbing beads.
- Do not remove tubes from magnetic rack and add **200 μl of 70 % ethanol** without disturbing beads (use freshly made 70% ethanol to ensure accurate concentration.)
- Wait for **30s**(leave cap open.)
- Remove and discard supernatant without disturbing beads.
- Add again **200 μl of 70 % ethanol**.
- Wait for **30s** (leave cap open.)
- Remove and discard supernatant without disturbing beads.
- **Make sure to remove all traces of ethanol** (use a 10μl micropipette if necessary.)
- Dry for **5 min** (leave cap open.)
- Do not remove tubes from magnetic rack and add **15μl of buffer EB**.
- Close cap.
- Remove strip from magnetic rack.
- Vortex + pulse spin.
- Wait for **2min**.
- Move strip back to magnetic rack and wait for **2 to 5min.**
- Transfer the **15μl** of DNA to a new strip.

##### 5. PCR enrichment of adaptor-ligated DNA

Thaw NEBNext Ultra II Q5 Master Mix. Homogenize by pipetting + Pulse spin. DO NOT VORTEX. Leave on Ice.

|  | **Master Mix:** |  |
| --- | --- | --- |
|  | x 1 reaction | x 96 reactions |
| NEBNext Ultra II Q5 Master Mix | 25 µl | 2400 µl |
| Primer F (10µM) | 2 µl | 192 µl |
| Primer R (10µM) | 2 µl | 192 µl |
| Master mix to add to each tube | 29 µl |  |

With

Primer F = AATGATACGGCGACCACCG*A

Primer R = CAAGCAGAAGACGGCATACG*A

(synthesized by IDT (Integrated DNA Technologies, Inc.), HPLC Purification, * = Phosphorothioate Bond.)

###### Vortex + pulse spin master mix

- Add 29 μl of master mix to tubes containing the 15 μl of adaptor ligated DNA.
- Vortex.
- Pulse spin.
- Place in a thermocycler. **<u>Enable heated lid</u>** and run the following program:

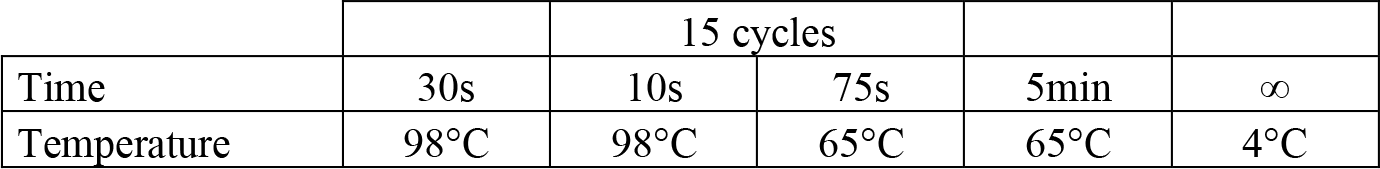

##### 6. Quantification, equimolar pooling and re-concentration

Qubit - quantify 2 μl of each library.

In our experience, a clean-up step of the librairies is not necessary. Prepare 6 pools of 16 samples in 1.5ml LoBind tubes.

Pool 100 ng of each library (you want a final quantity of 100-500ng DNA per pool after concentration to fit with the requirements of the MYbaits® protocol.)

**<u>On beads Concentration of DNA</u>** (Agencourt AMPure XP Purification system (Beckman Coulter)). Concentration of DNA is required, as MyBaits require 7 μl of starting material.

Remove Ampure beads from fridge. Make sure they are at room temperature (RT) before use.

- Vortex Ampure beads.
- Add 0.8 vol. of Ampure beads for 1 vol. of DNA.
- Vortex + pulse spin.
- Incubate for **5min** at RT.
- Move tubes to magnetic rack and wait for **2 to 5 min**.
- Remove and discard supernatant without disturbing beads.
- Do not remove tubes from magnetic rack and add **200 μl of 70 % ethanol**without disturbing beads (use freshly made 70% EtOH to ensure accurate concentration.)
- Wait for **30s** (leave cap open.)
- Remove and discard supernatant without disturbing beads.
- Add again **200 μl of 70 % ethanol**.
- Wait for **30s** (leave cap open.)
- Remove and discard supernatant without disturbing beads.
- **Make sure to remove all traces of Ethanol** (use a 10μl micropipette if necessary.)
- Dry for **5 min** (leave cap open.)
- Do not remove tubes from magnetic rack and add **9μl**of buffer EB.
- Close cap.
- Remove tubes from magnetic rack.
- Vortex + spin pulse.
- Wait for **2min**.
- Move strip back to magnetic rack and wait for **2 to 5min**.
- Transfer the **9μl**of DNA to a new 1.5ml LoBind tube.

Qubit quantify 1 μl of each library (dilution 1:2).

Check library profile on an Agilent Bioanalyzer with a High Sensitivity DNA Analysis Kit (load 1μl of library dilution 1:2).

##### 7. Capture of UCEs

UCEs were captured on each pool using the probes designed by Faircloth et al. (2015) and synthetisized by MYbaits®. Capture was performed with the MYbaits® kit following manufacturer’s instruction (http://www.mycroarray.com/pdf/MYbaits-manual-v3.pdf).

##### 8. PCR Enrichment of captured libraries and final clean-up

At this step of the process we have obtained libraries and Streptavidin beads mixed in 30 μl of 10mM Tris-Cl, 0.05% TWEEN®-20 solution (pH 8.0 – 8.5).

Here we want to release the bead-bound UCE-enriched library from the baits and amplif the resulting fragments.

PCR enrichment of the captured fragment is performed on beads with the following mix. We performed 2 PCR reactions for each pool of samples.

| **Post-capture PCR Master Mix:** | Final concentration | x1 |
| --- | --- | --- |
| H2O up | - | 5µl |
| 2X KAPA Hifi HotStart ReadyMix | 1X | 25µl |
| Primer F (10µM) | 500nM | 2.5µl |
| Primer R (10µM) | 500nM | 2.5µl |
| Enriched Library (on-bead) | - | 15µl |
| Master mix to add to each well |  | 50µl |

With

Primer F = AATGATACGGCGACCACCG*A

Primer R = CAAGCAGAAGACGGCATACG*A

(synthesized by IDT (Integrated DNA Technologies, Inc.), HPLC Purification, * = Phosphorothioate Bond)

| **Program:** |  | 14 cycles |  |  |  |  |
| --- | --- | --- | --- | --- | --- | --- |
| Step | Activation | Denaturation | Annealing | Extension | Final Extension | End |
| Time | 2min | 20sec | 30sec | 45sec | 5min | ∞ |
| Temperature | 98°C | 98°C | 65°C | 72°C | 72°C | 10°C |

###### Remove Streptavidin beads

Pool the 2 PCR products obtained for each pool of samples in DNA LoBind tubes (1.5ml. Move the 6 tubes to a magnetic rack and wait for 2 to 5min.

Transfer supernatant to new tubes.

**Final Cleanup** (Agencourt AMPure XP Purification system (Beckman Coulter)

Remove Ampure beads from fridge. Make sure they are at room temperature (RT) before use.

- Vortex Ampure beads.
- Add 1V of Ampure beads for 1V of DNA (ie 100 μl of beads for 100 μl of DNA.)
- Vortex + pulse spin.
- Incubate for **5min**at RT.
- Move tubes to magnetic rack and wait for **2 to 5min**.
- Remove and discard supernatant without disturbing beads.
- Do not remove tubes from magnetic rack and add **200 μl of 70 % ethanol** without disturbing beads (use freshly made 70% EtOH to ensure accurate concentration.)
- Wait for **30s** leave cap open.)
- Remove and discard supernatant without disturbing beads.
- Add again **200 μl of 70 % ethanol**.
- Wait for **30s** (leave cap open.)
- Remove and discard supernatant without disturbing beads.
- **Make sure to remove all traces of ethanol** (use a 10μl micropipette if necessary.)
- Dry for **5 min** (leave cap open.)
- Do not remove tubes from magnetic rack and add **25μl of buffer EB**.
- Close cap.
- Remove tubes from magnetic rack.
- Vortex + pulse spin.
- Wait for **2min**.
- Move strip back to magnetic rack and wait for **2 to 5min**.
- Transfer the **25μl** of DNA to new LoBind tubes.

##### 9. Final quantification and equimolar pooling prior to sequencing

- Qubit Quantify 2μl of each.
- Check library profile on an Agilent Bioanalyzer with a High Sensitivity DNA Analysis Kit (1μl undiluted library).
- Perform KAPA quantification.
- Equimolar pooling of the 6 librairies.

### Additional file 3

Adapters (Ultramer Oligos, TruGrade synthesis, Standard Desalting) were synthesized by IDT (Integrated DNA Technologies, Inc.) /5Phos/ = 5’ Phosphorylation; * = Phosphorothioate Bond, F=forward, R=reverse).

#### Adapter were hybridized as follows

**25μM oligo preparation:**

- Thaw 100μM oligos. Vortex and spin well.
- Pool each pair of oligos (F/R) in DNA LoBind tubes (1.5ml) as follows:

- Vortex and spin well the oligo’s pools and split each pool in tow PCR tubes (0.2ml)

|  | Initial Concentration | Final Concentration |
| --- | --- | --- |
| P1 F or P2 F | 100μM | 25μM |
| P1 R or P2 R | 100μM | 25μM |
| AB Buffer | 10X | 1X |
| Ultra Pure H2O | | QS 100μl |

#### Adapters hybridization

- Place the PCR tubes in a thermocycler:

- Vortex and spin well the tubes.
- Qubit quantify 2μl of diluted adapters (1/20e.)
- Adjust adapters concentration to 1.5μM if necessary and make aliquots.
- Store the aliquots at −20°C.

| Program: |  | 15 cycles (with -1.7°C/cycle) | 40 cycles (with -1°C/cycle) |  |
| --- | --- | --- | --- | --- |
| Time | 2min | 1min | 1min | ∞ |
| Temperature | 95°C | 95°C | 69°C | 10°C |

Adapters (1.5μM) were stored in tube strips for usage comfort.

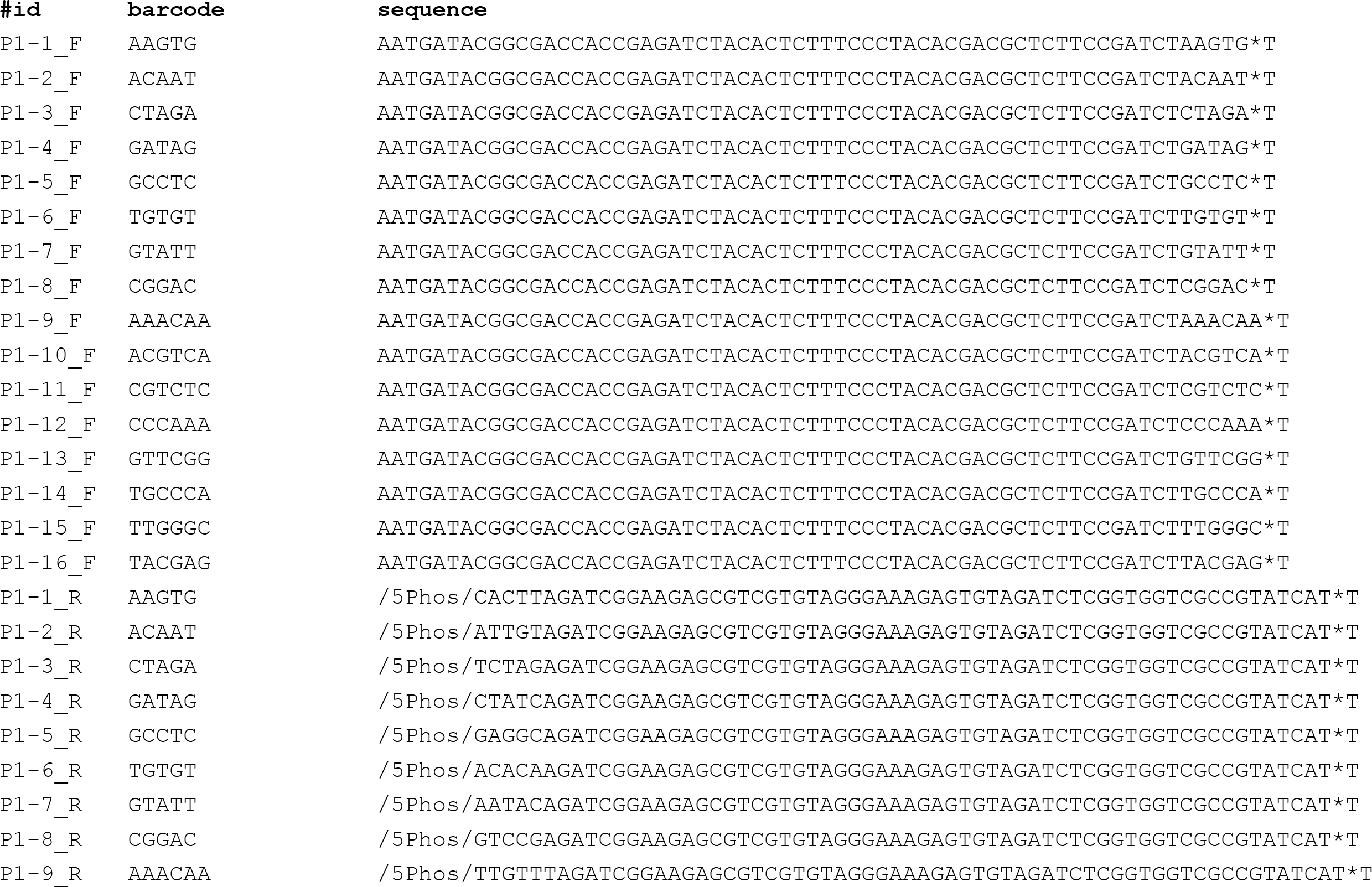

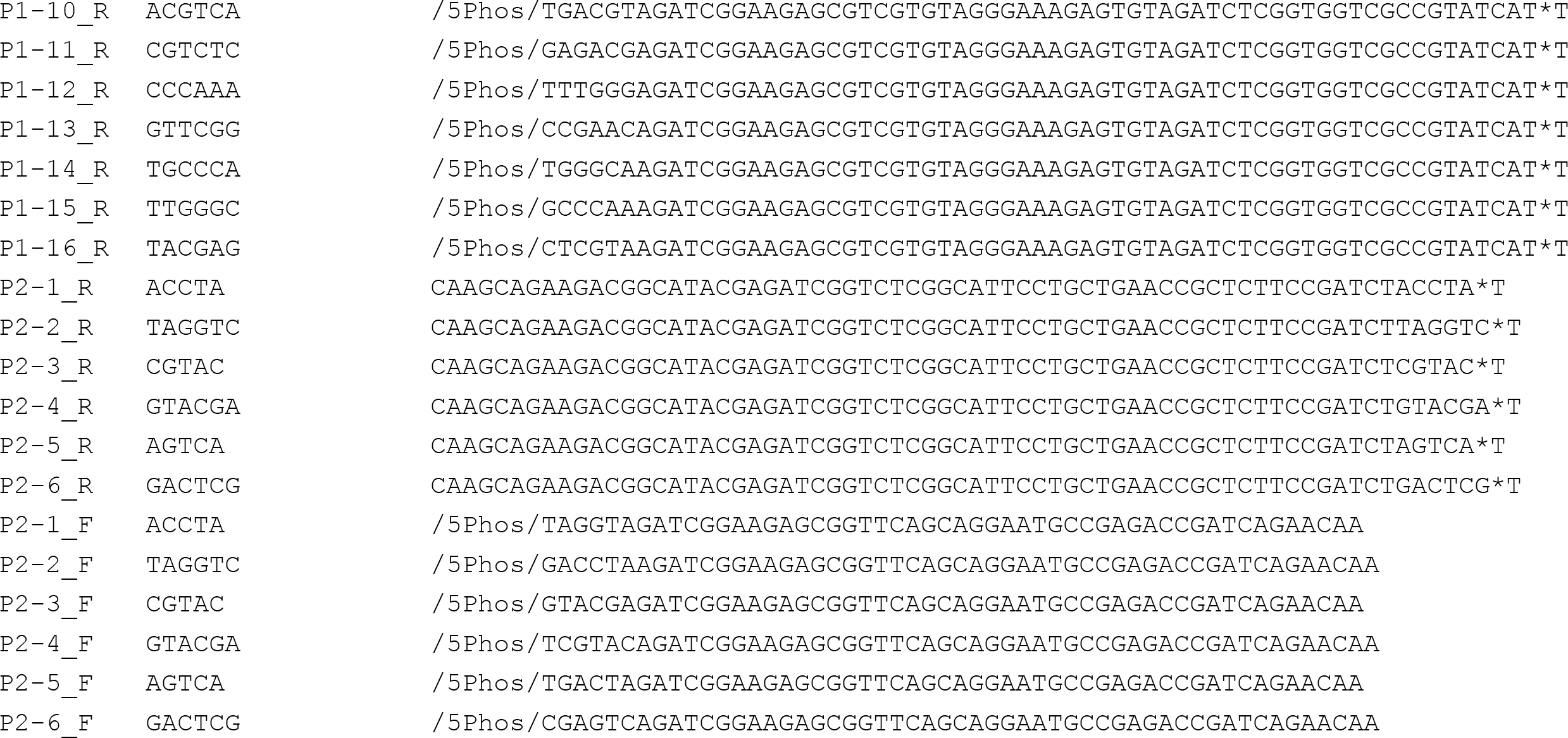

### Additional file 4

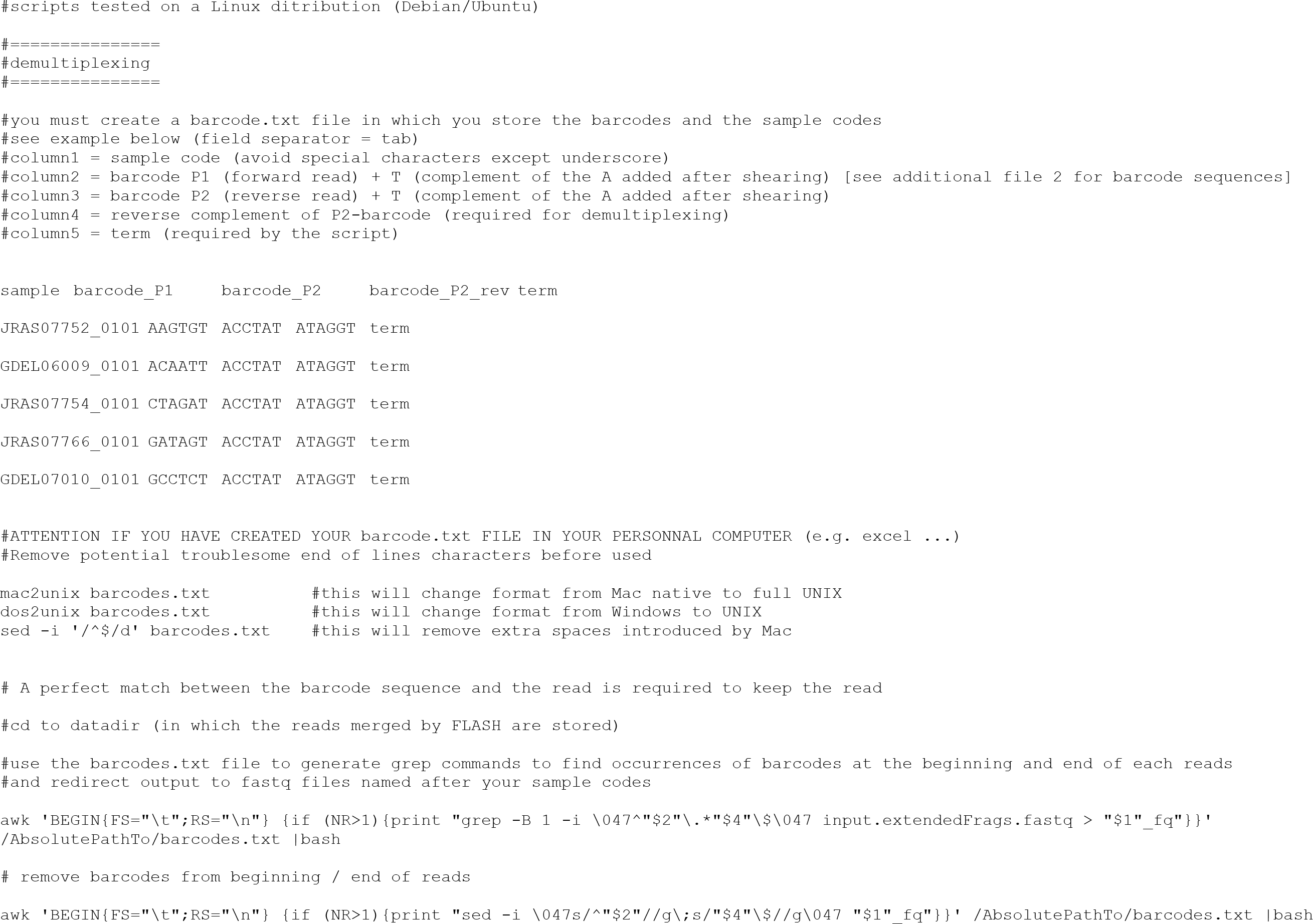

### Additional file 5

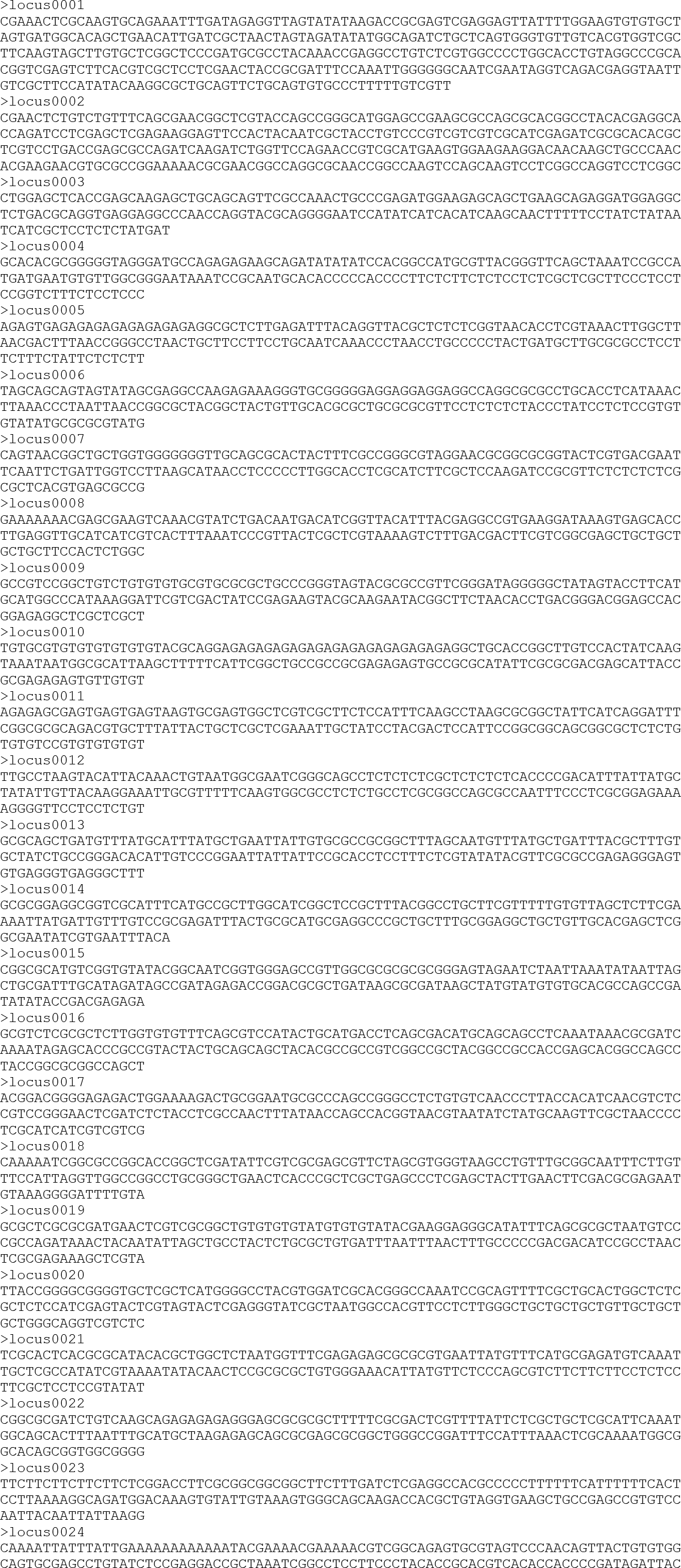

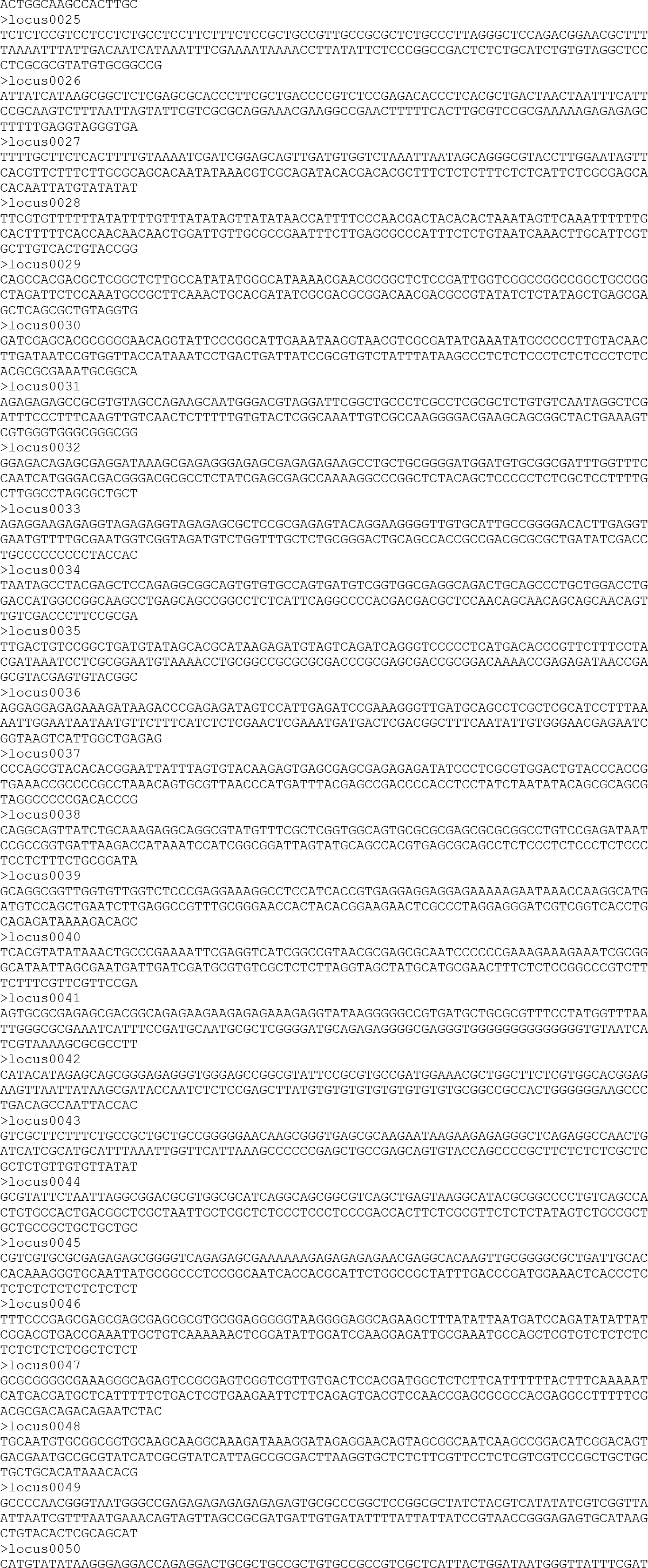

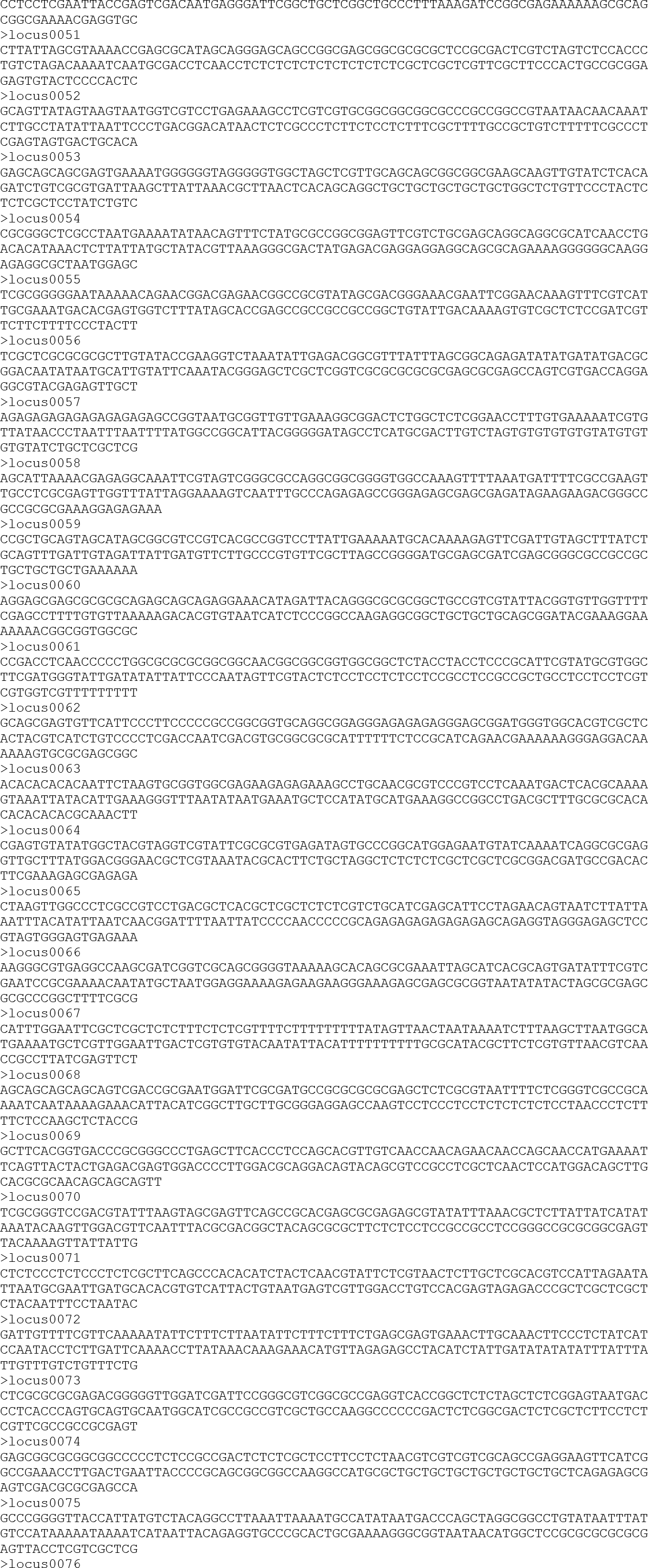

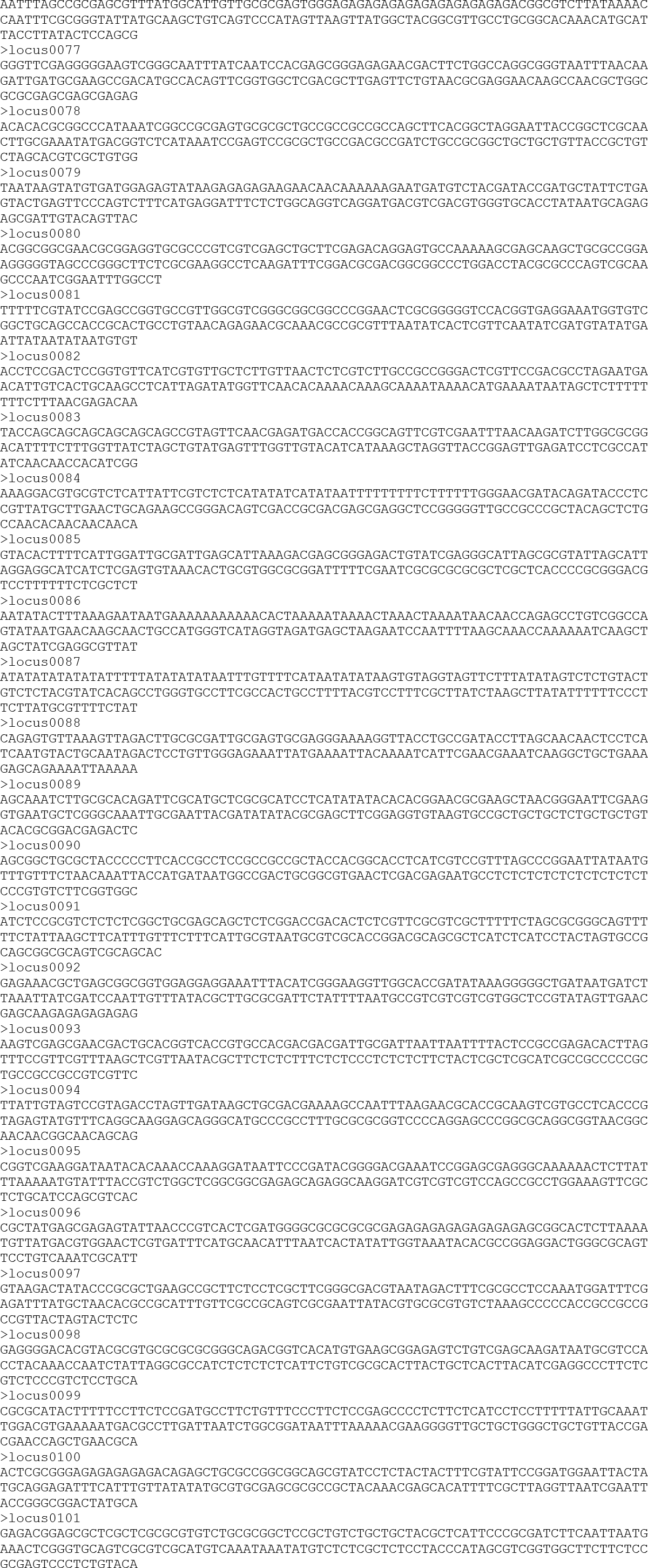

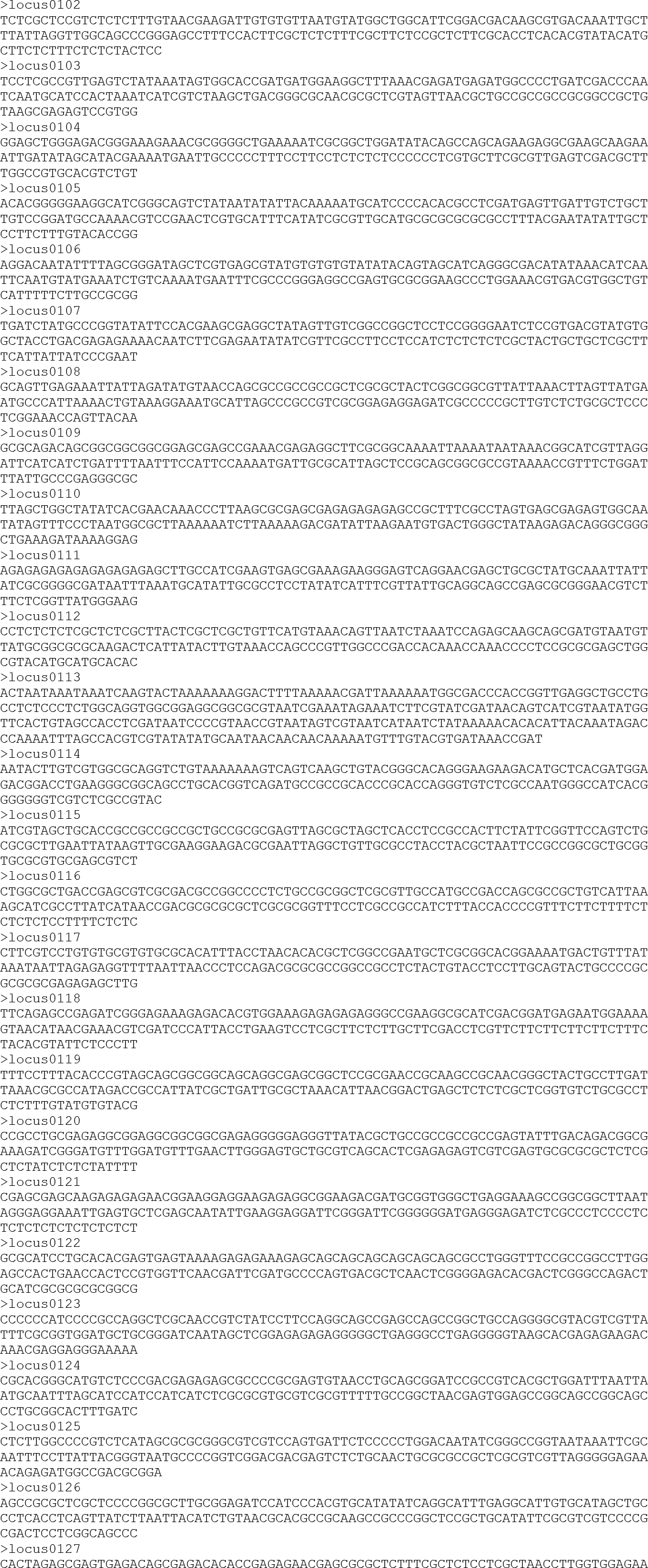

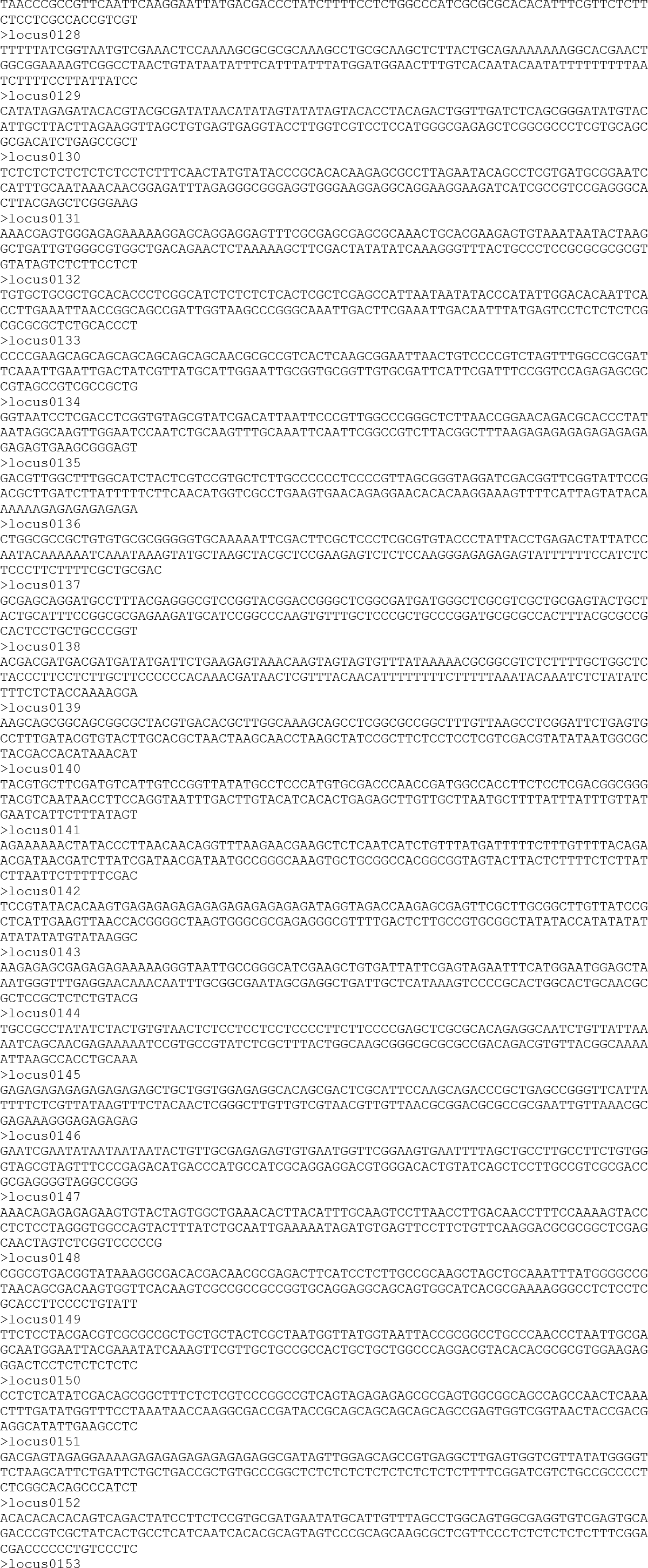

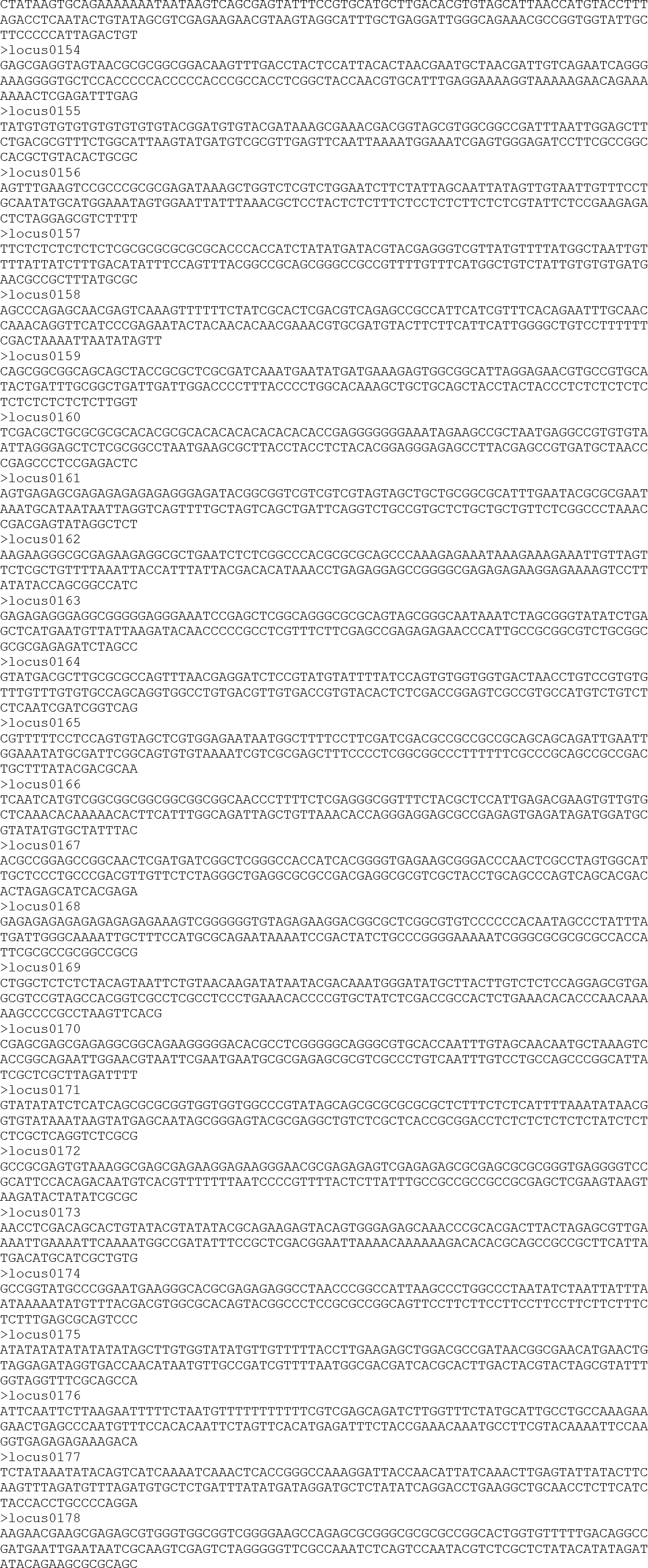

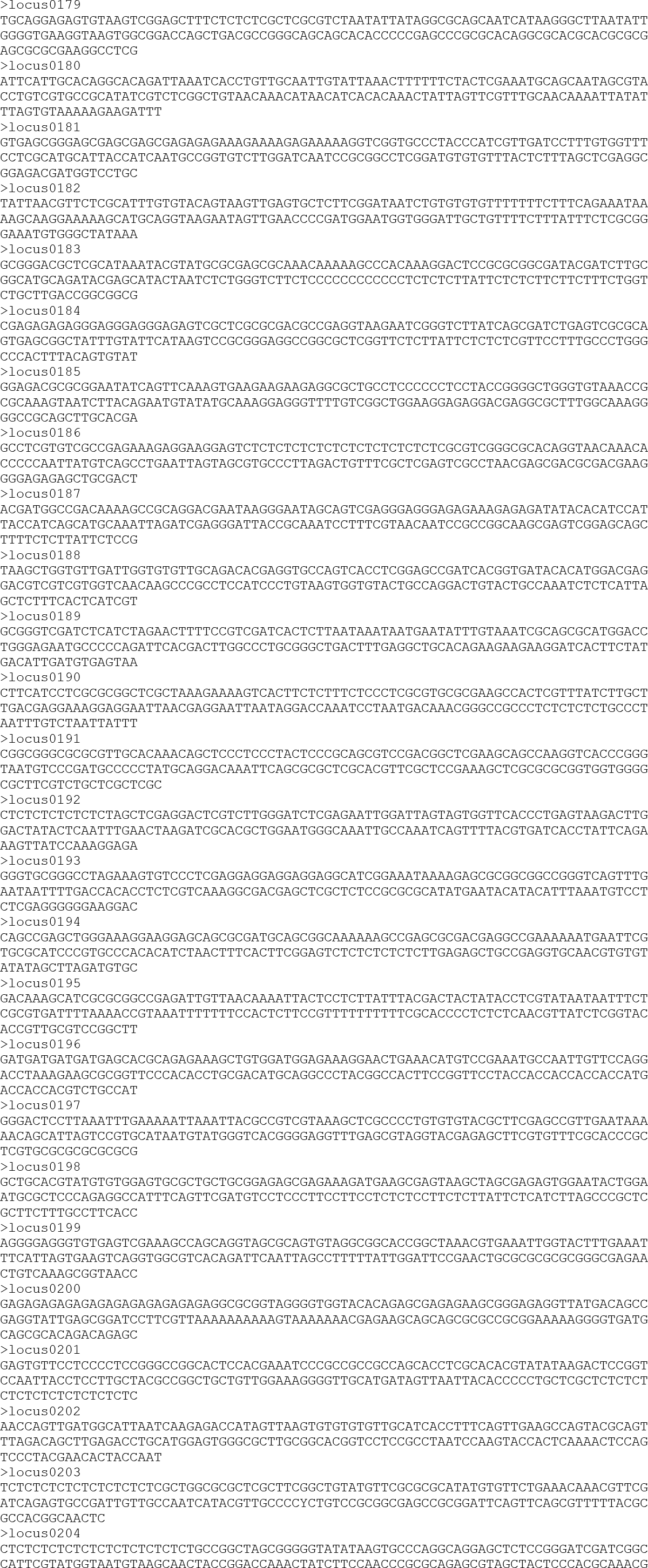

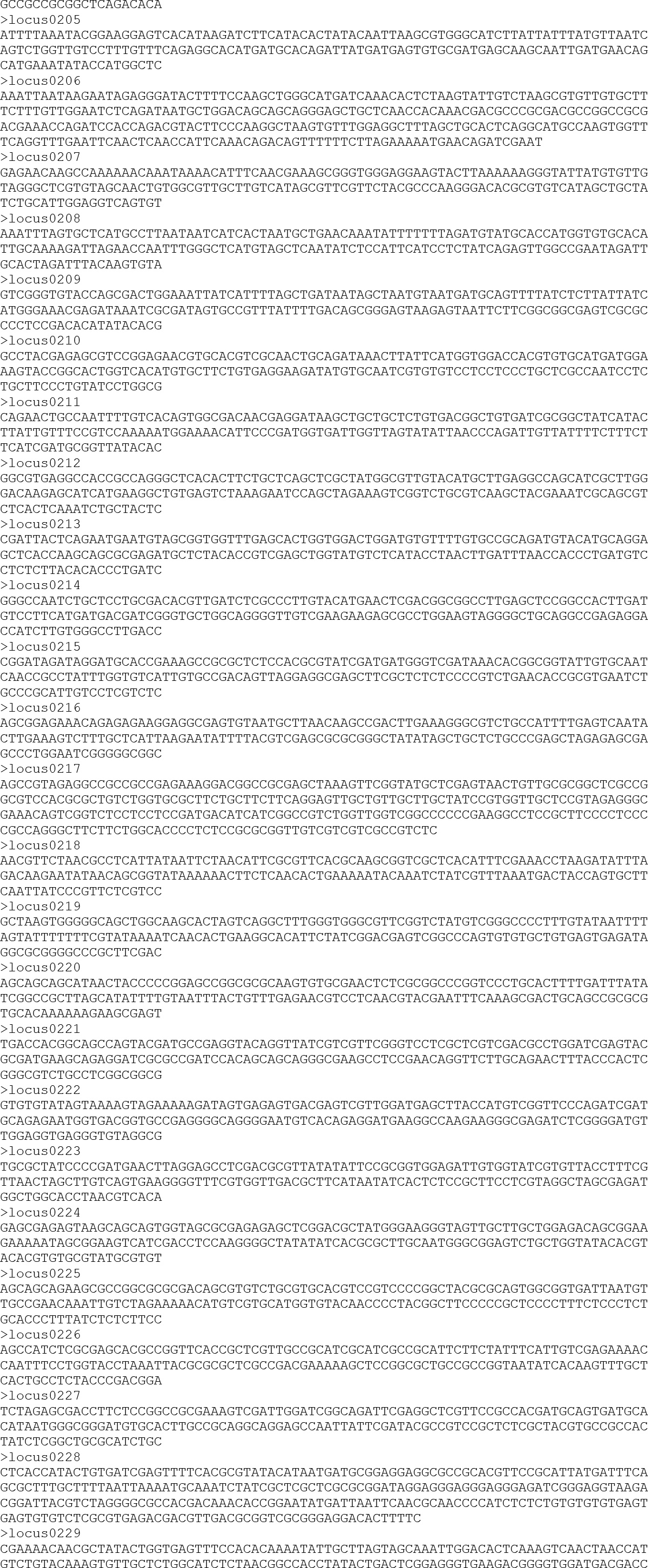

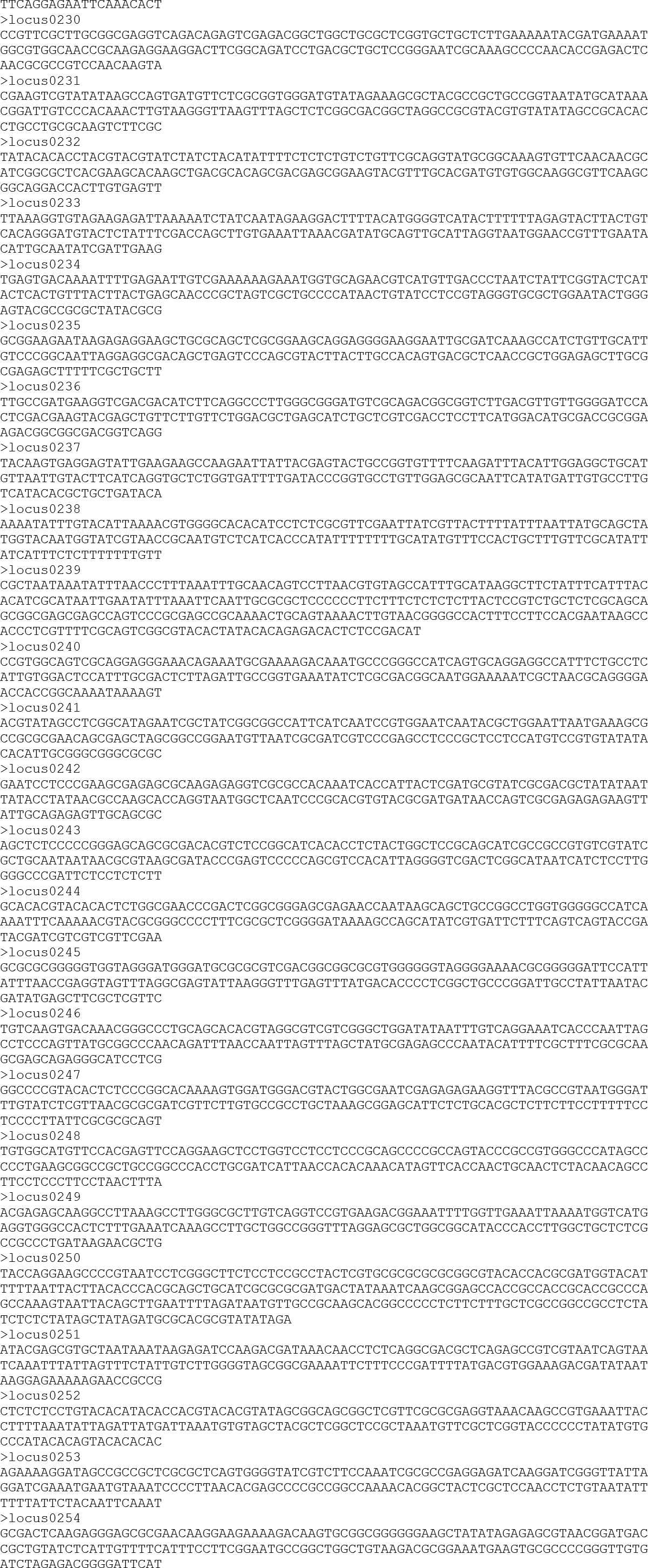

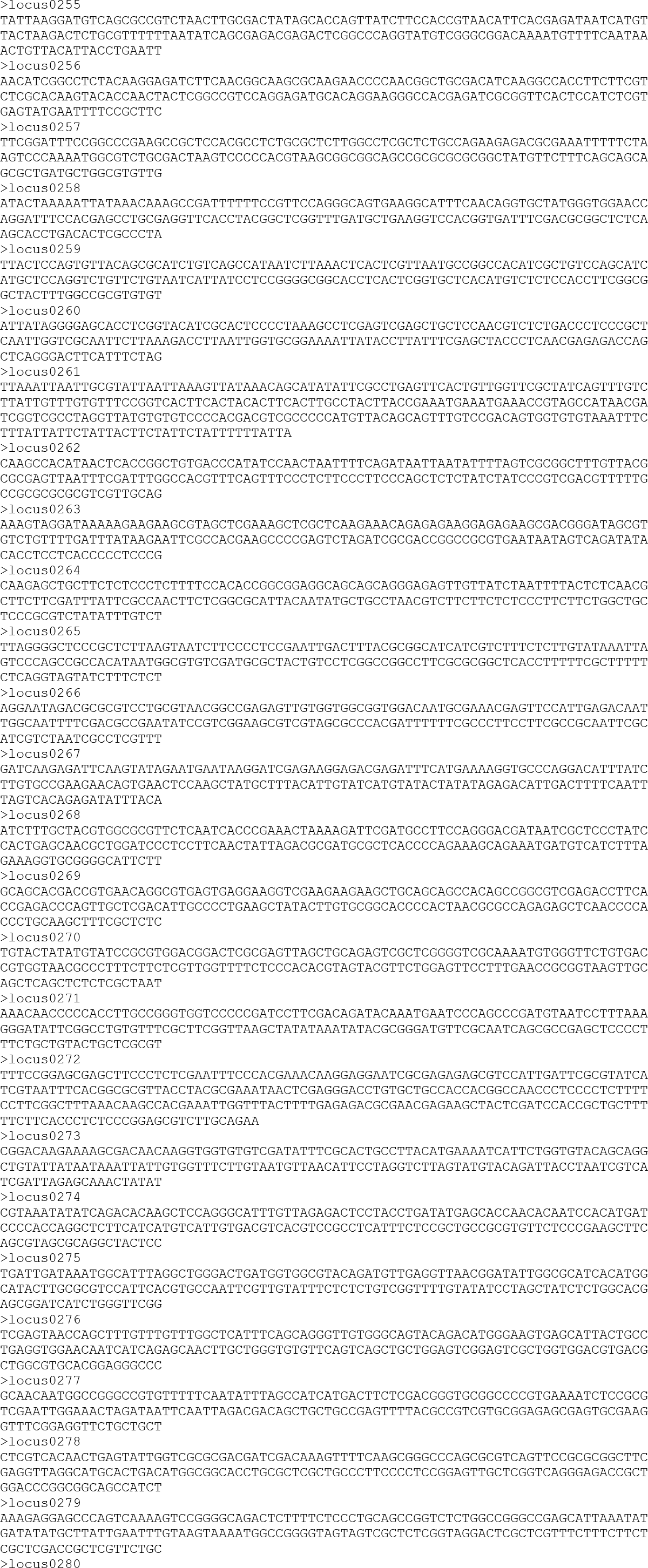

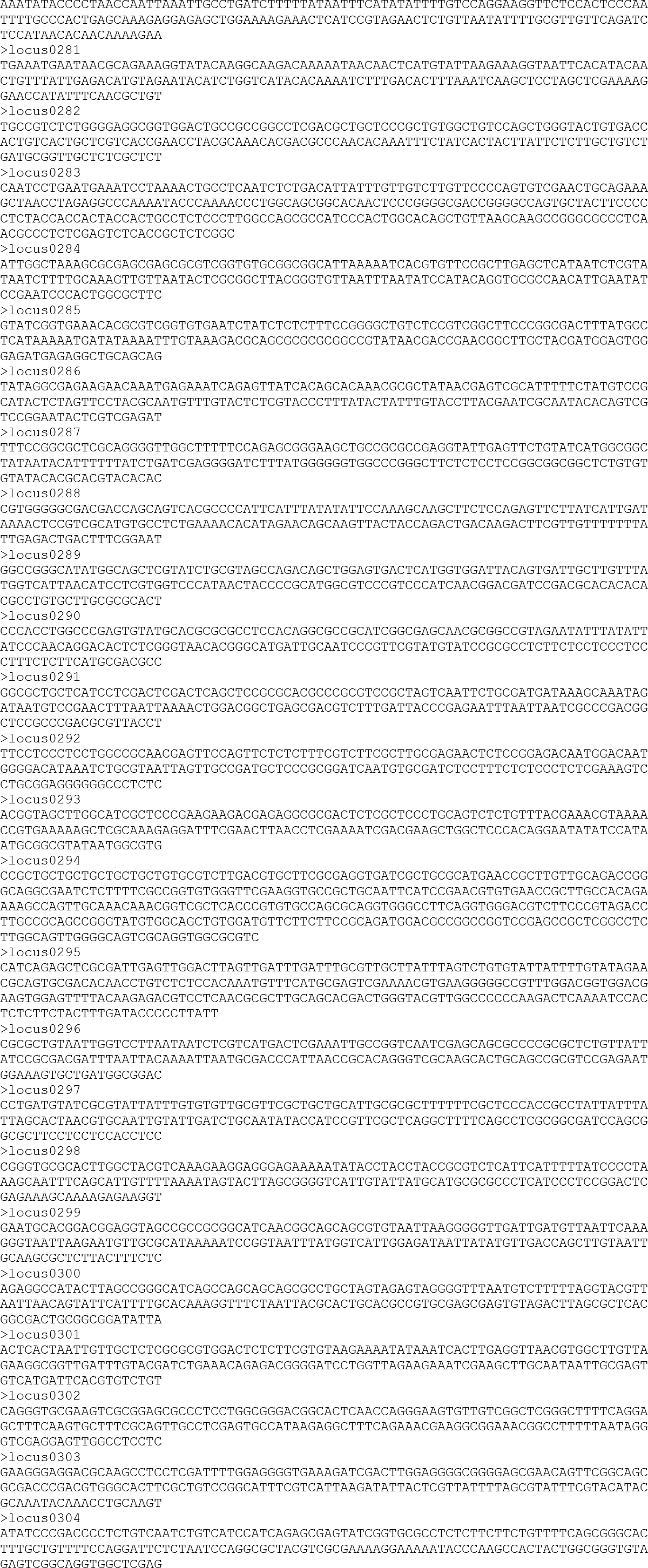

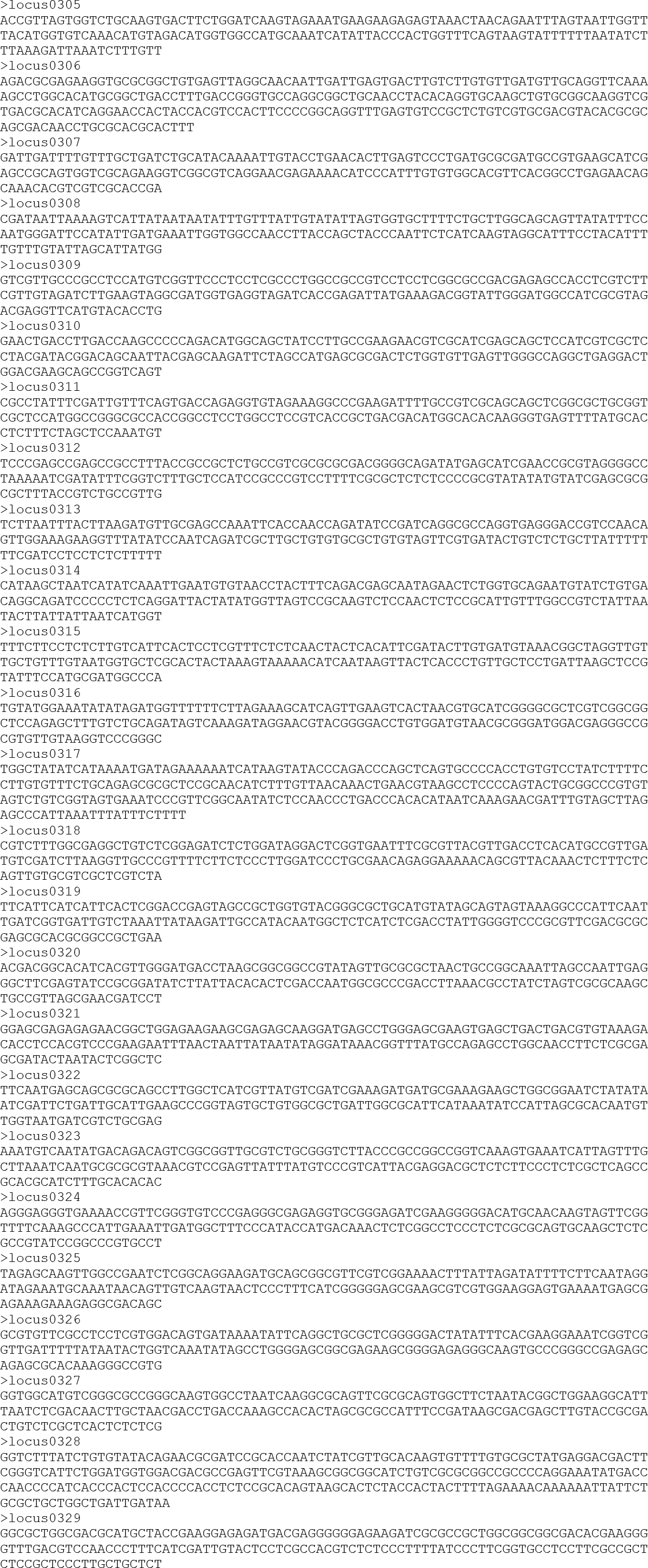

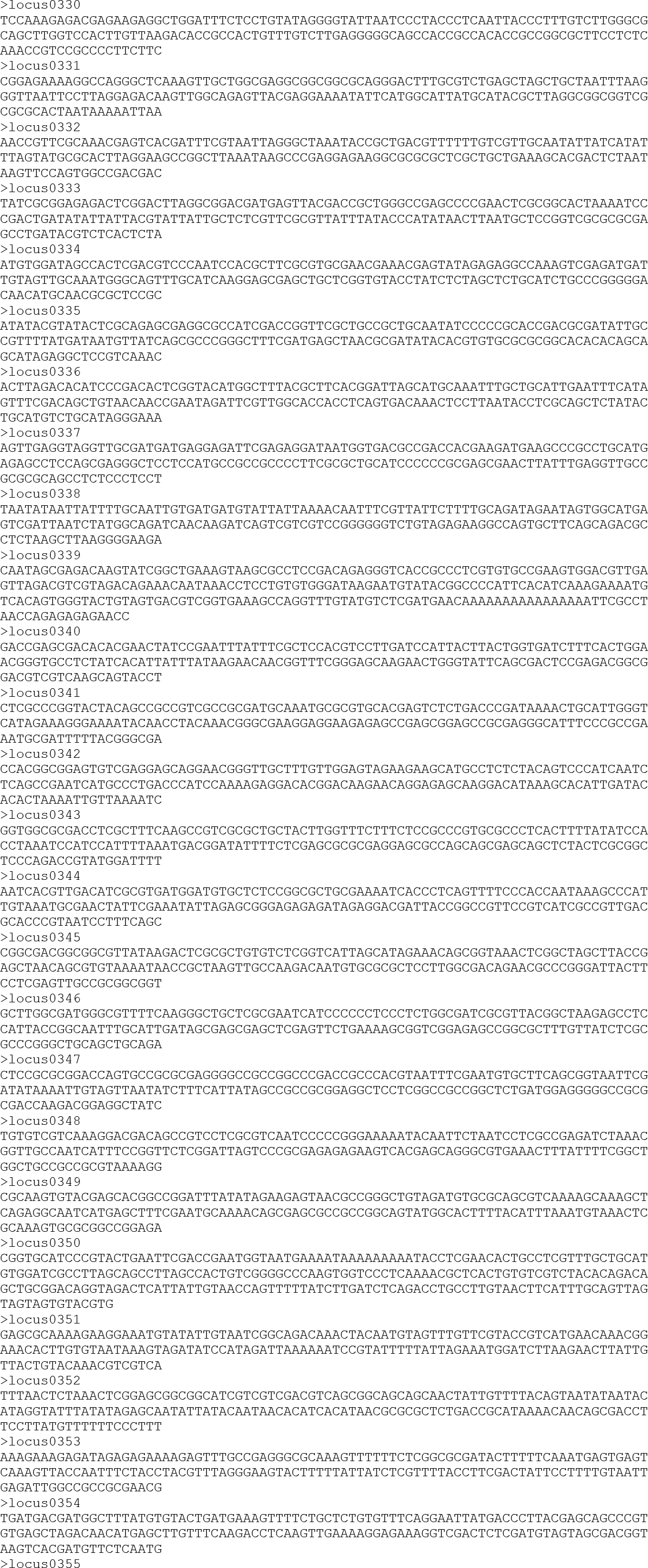

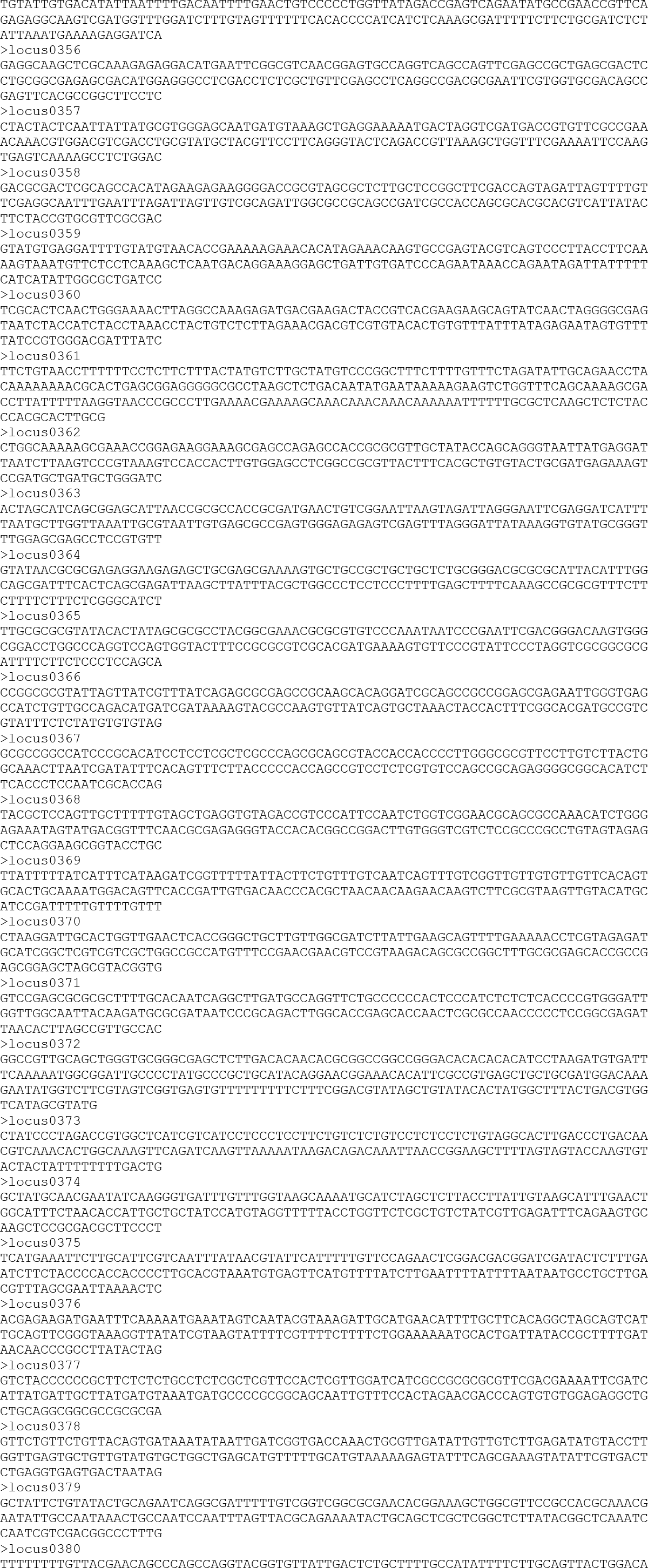

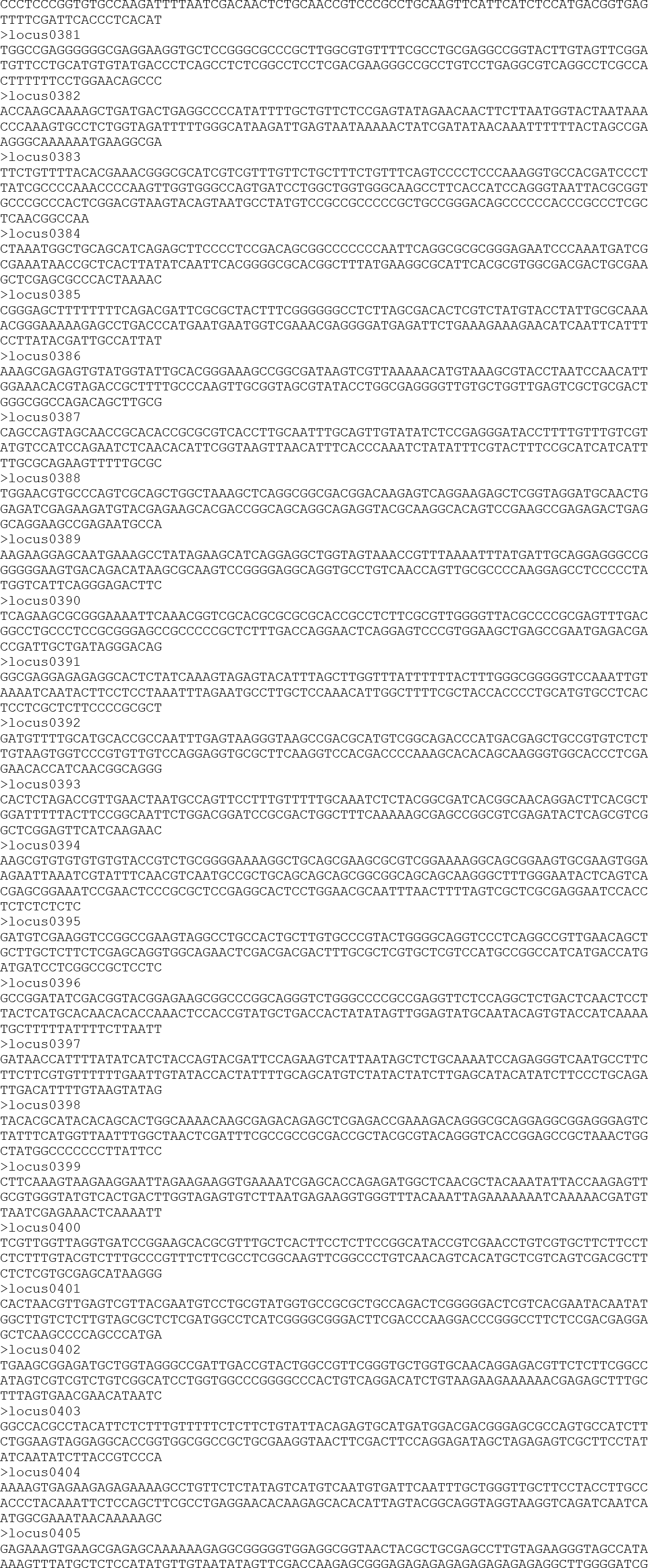

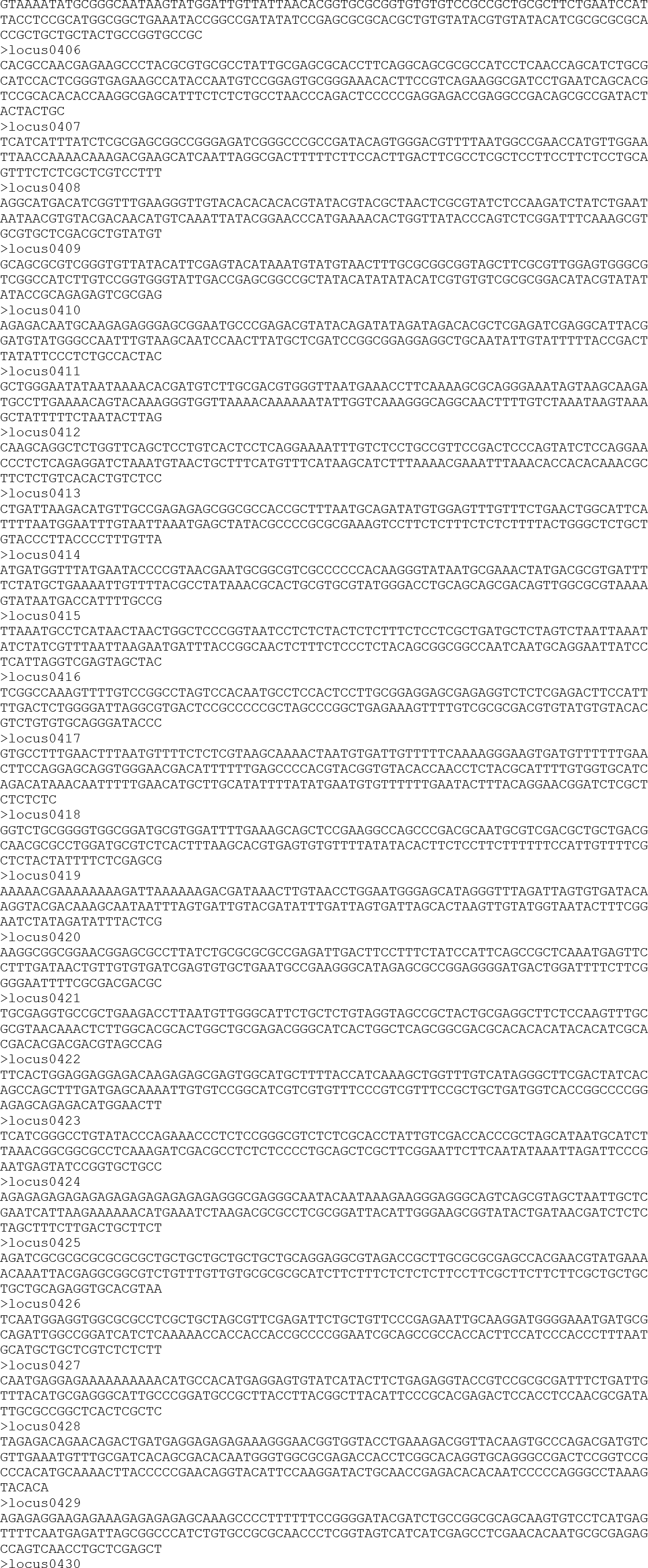

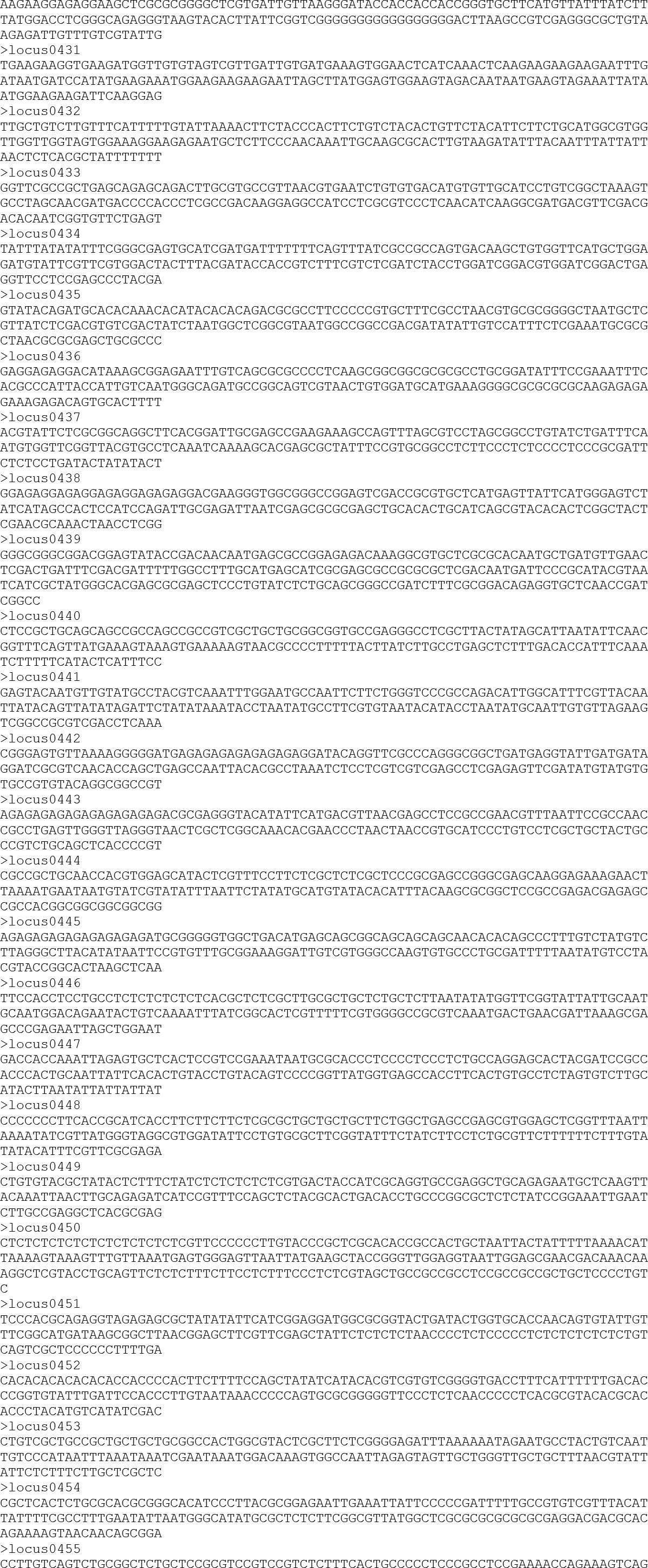

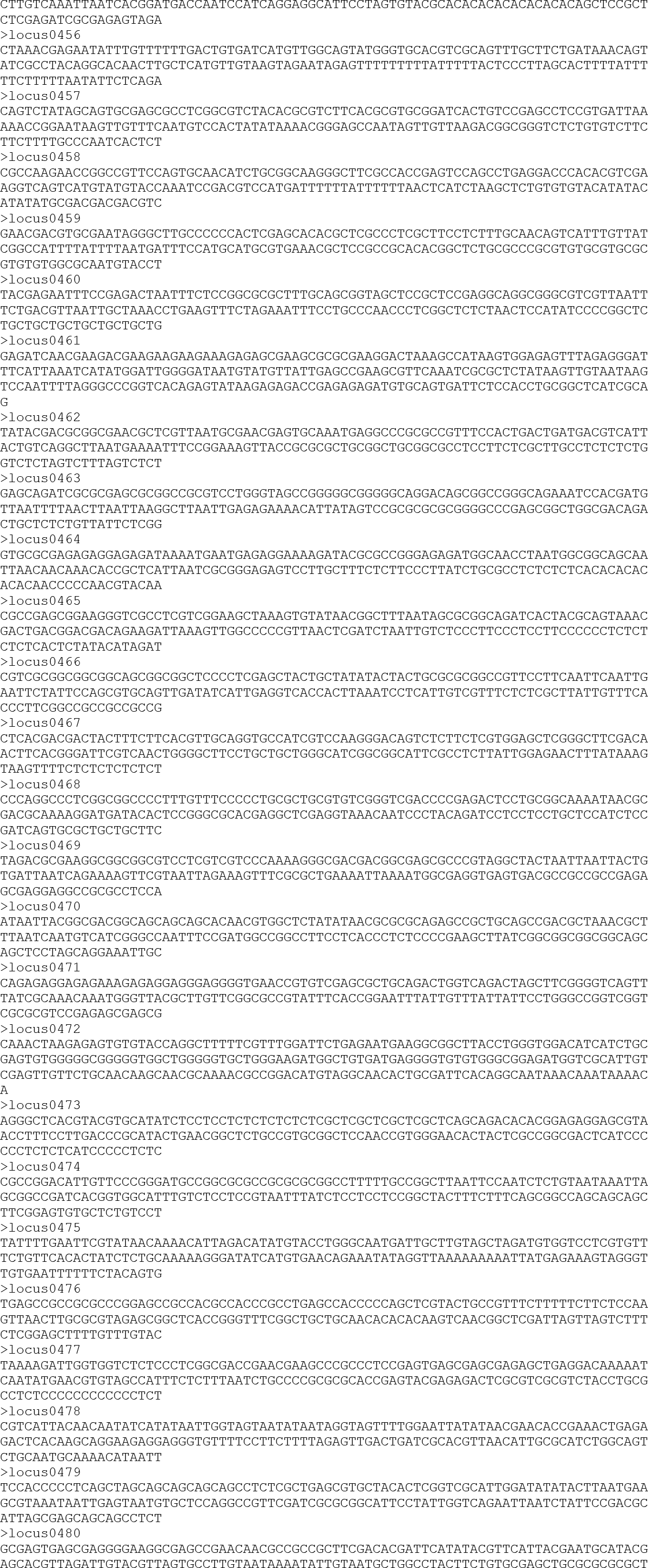

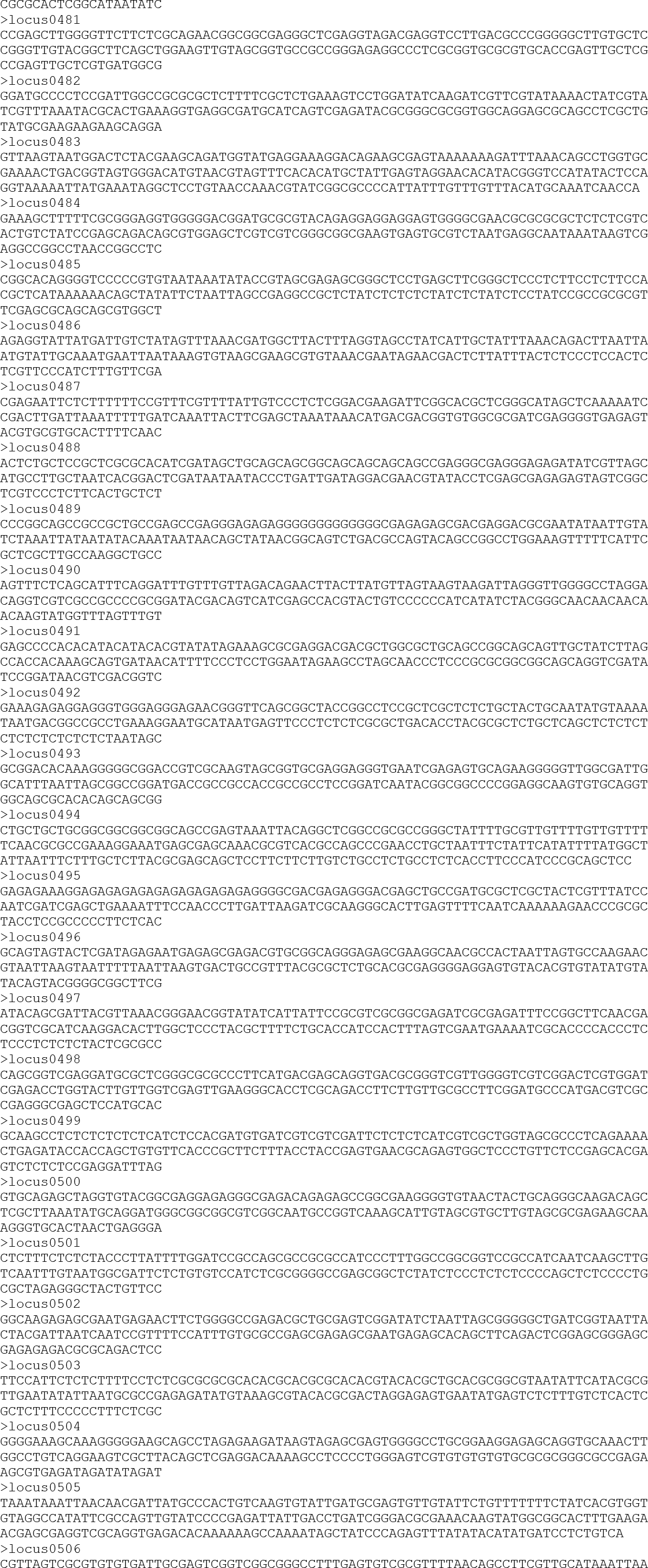

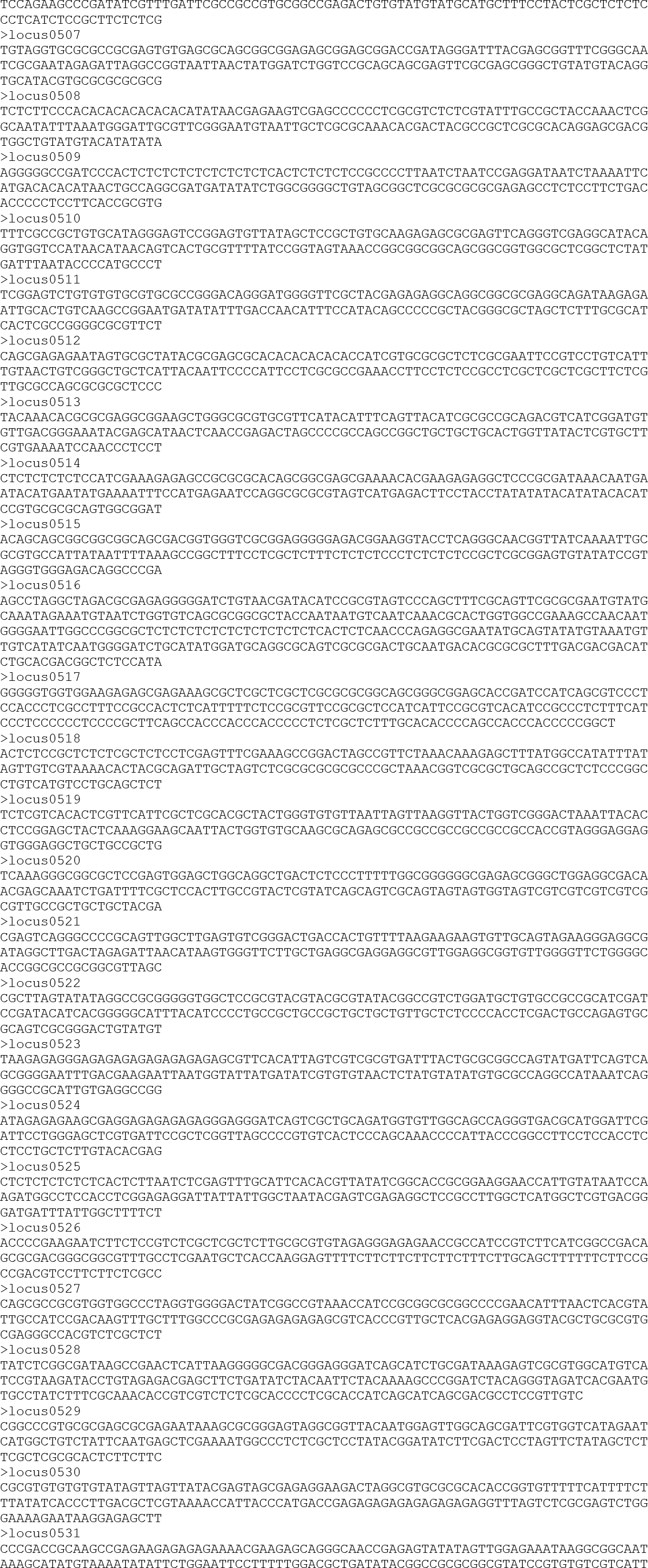

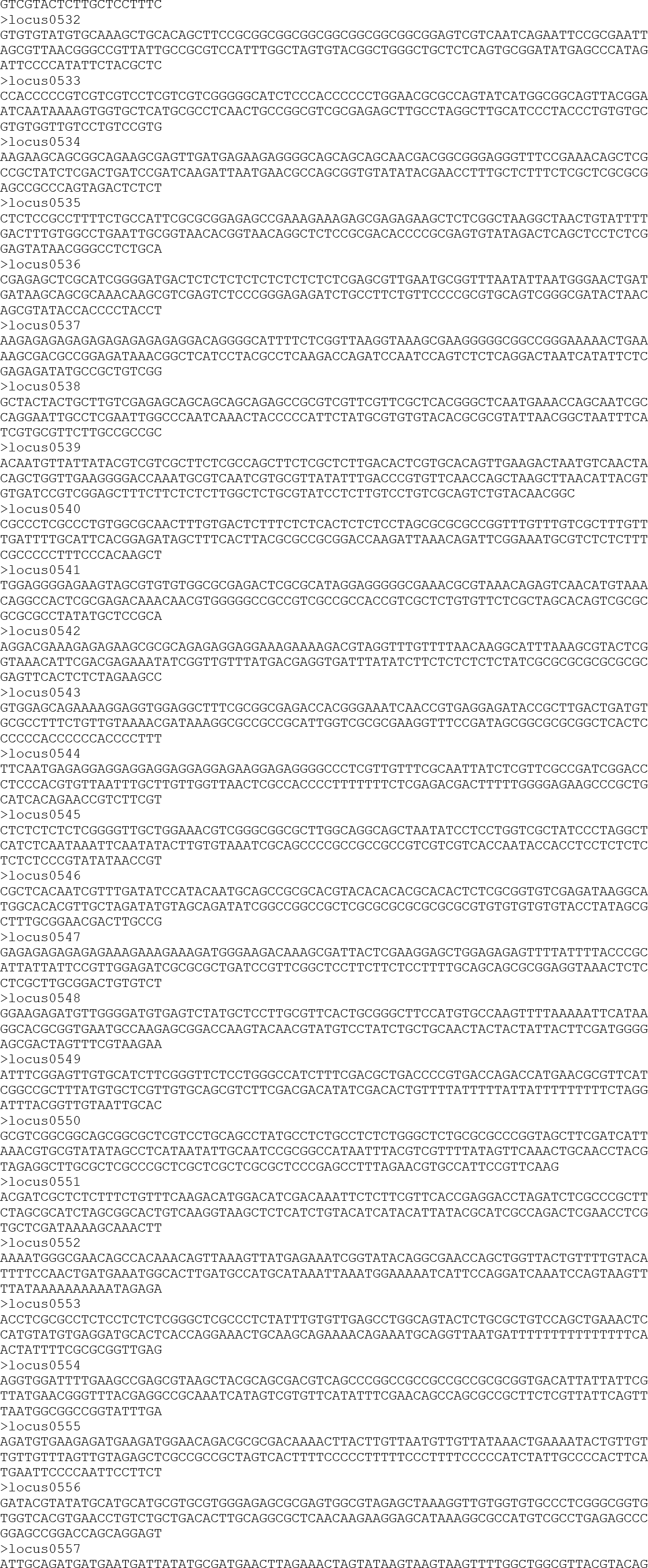

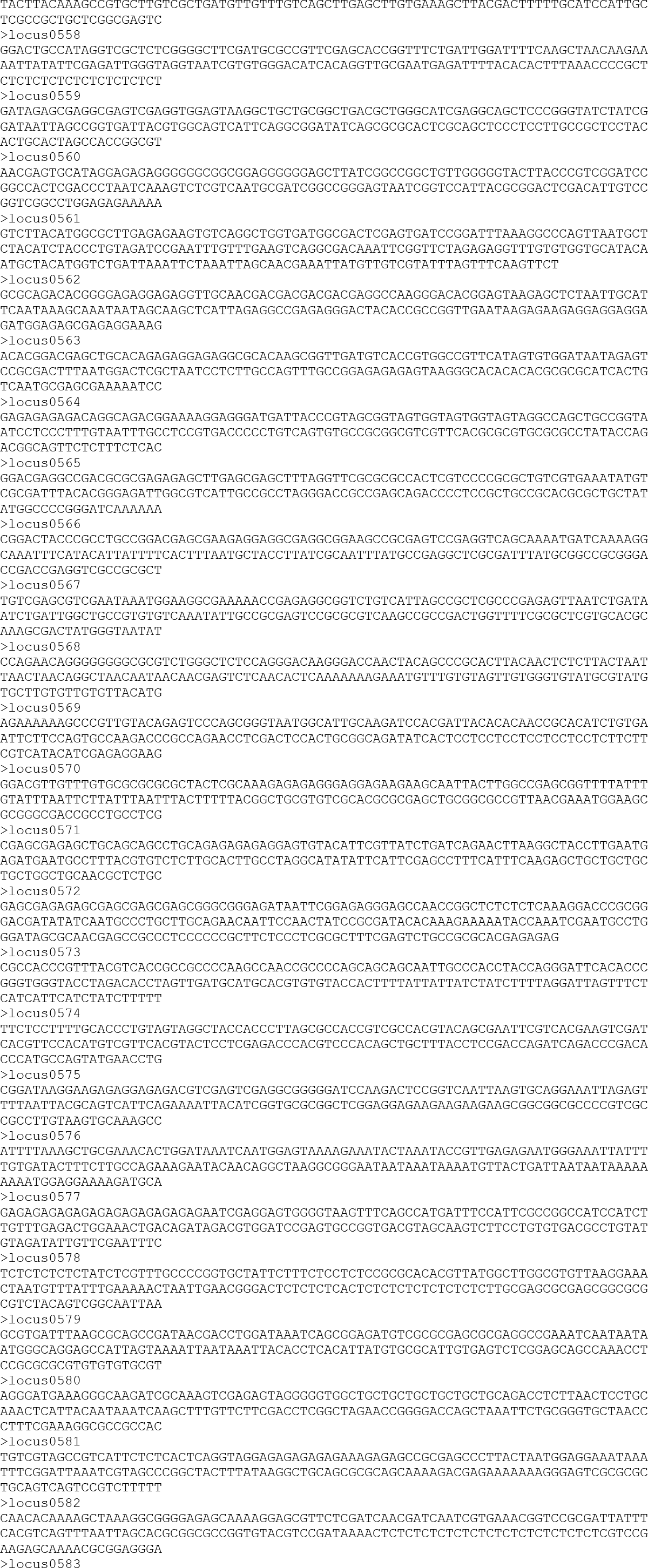

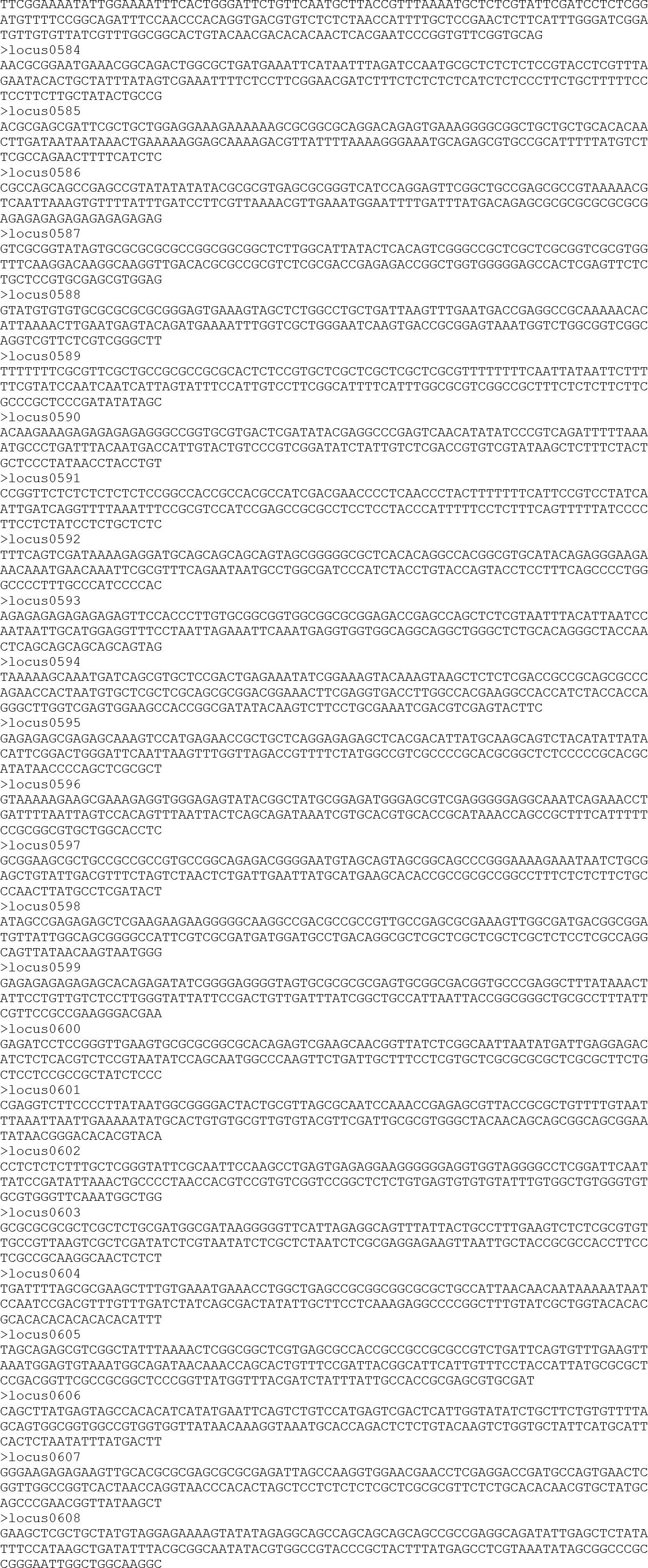

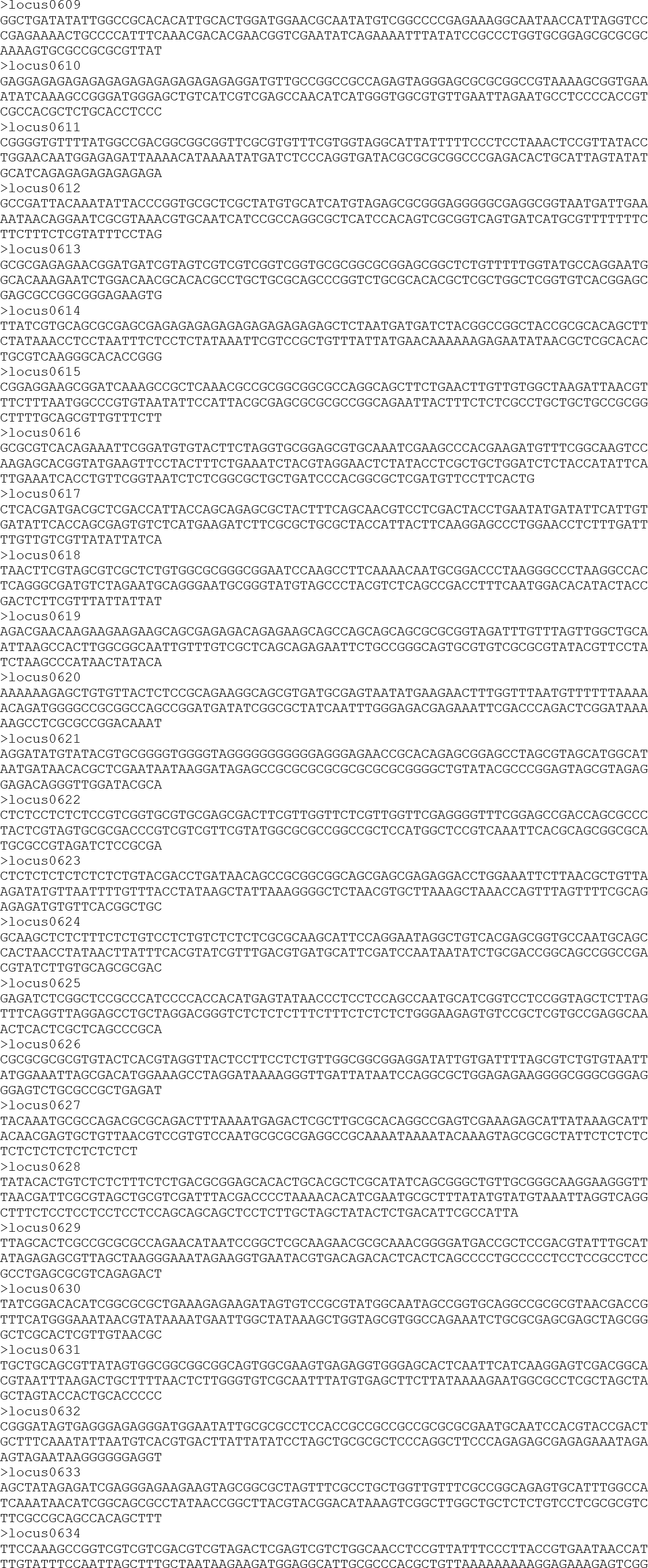

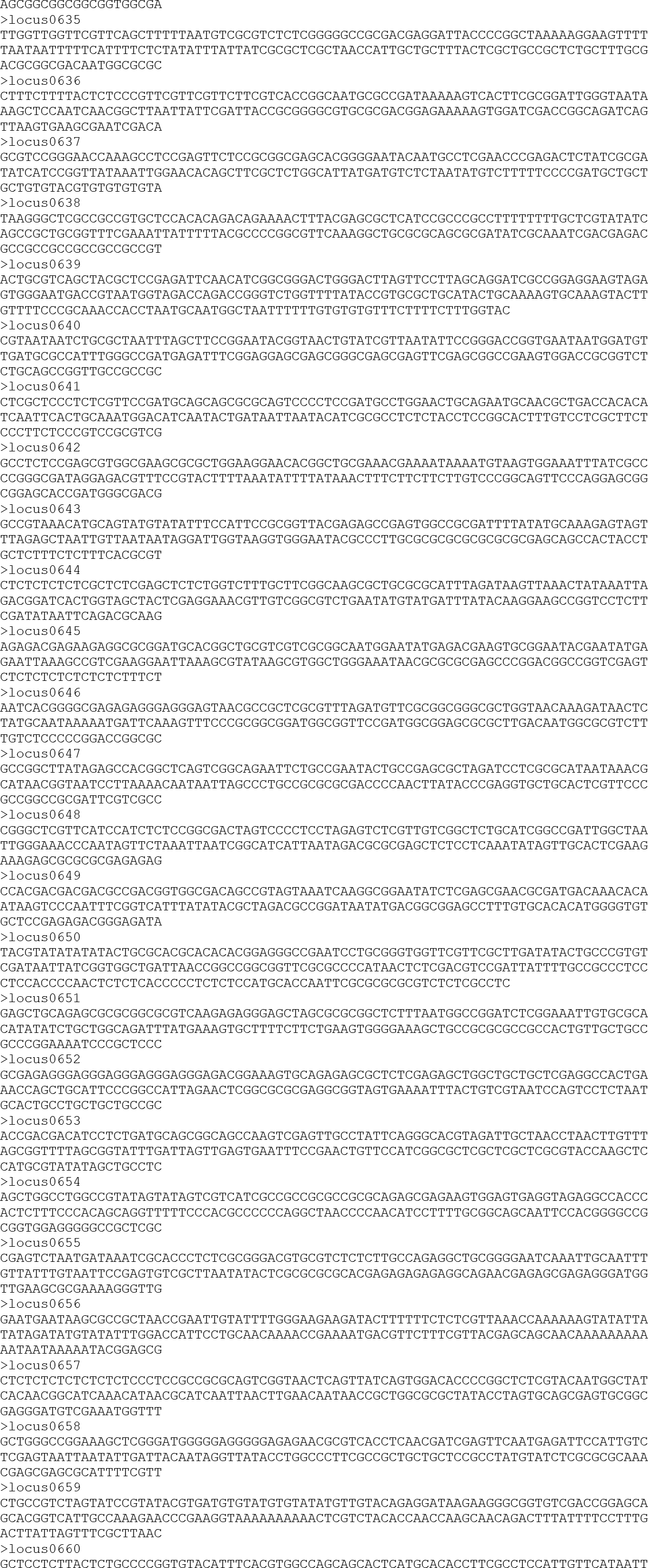

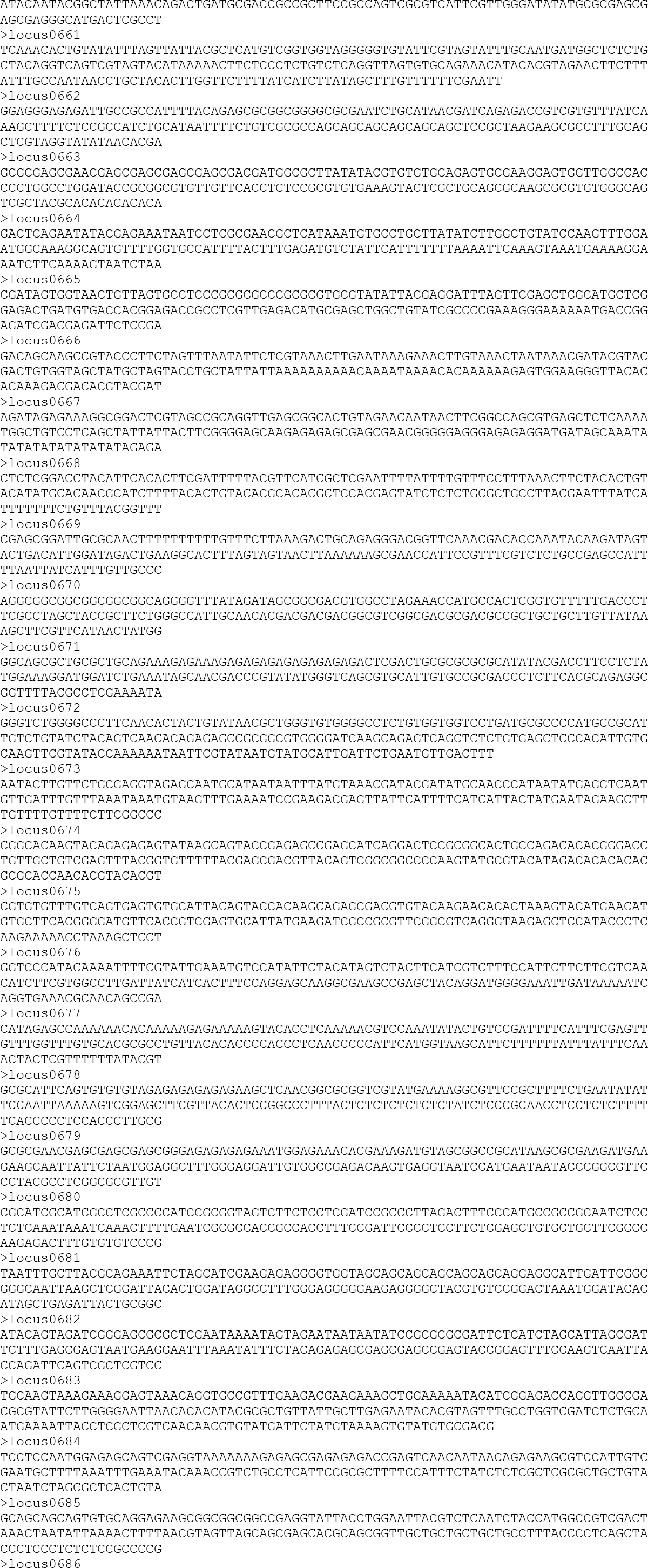

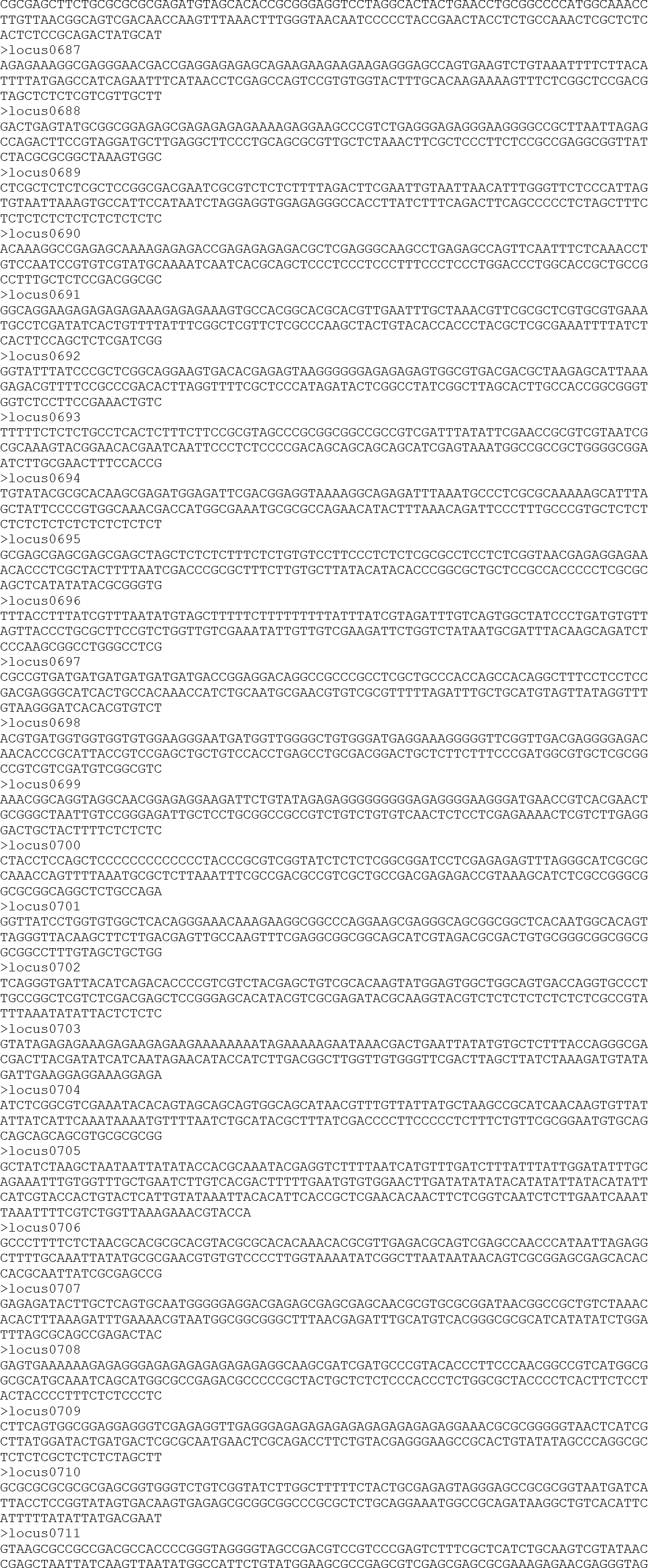

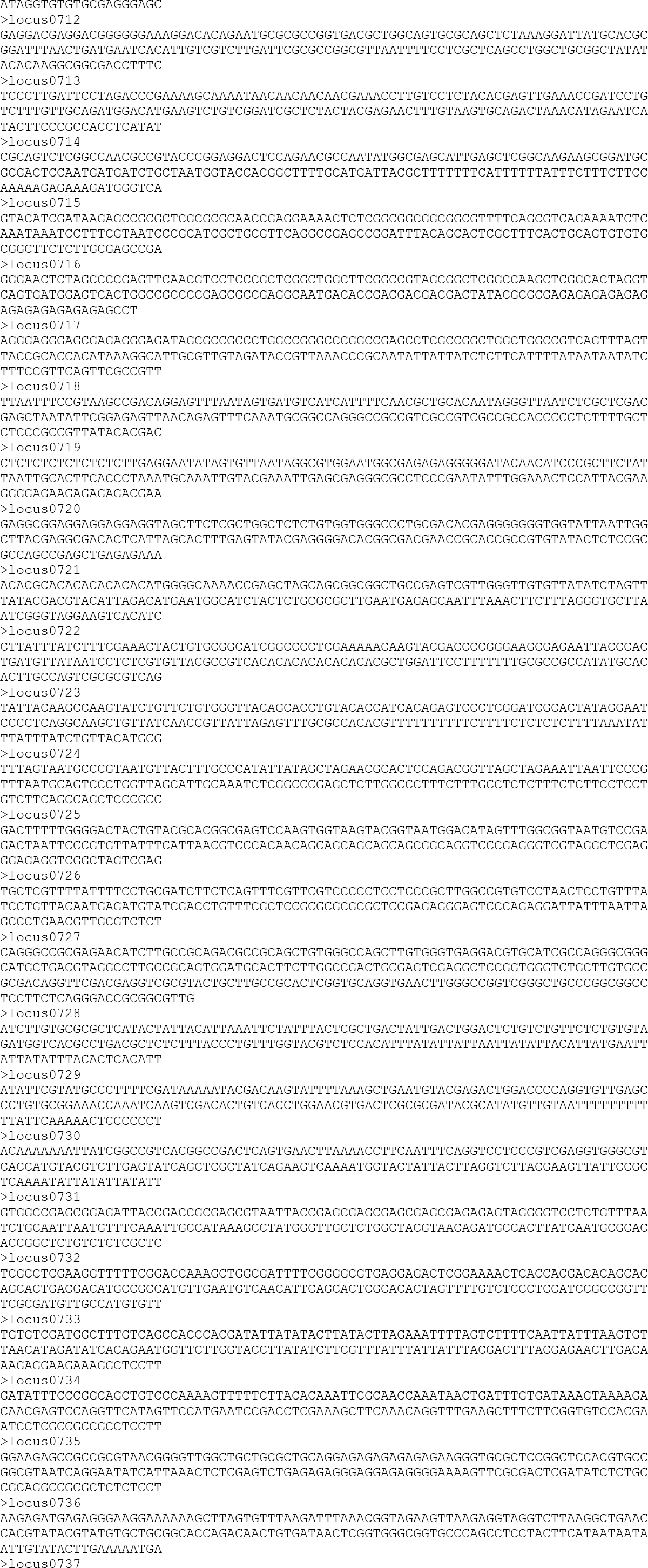

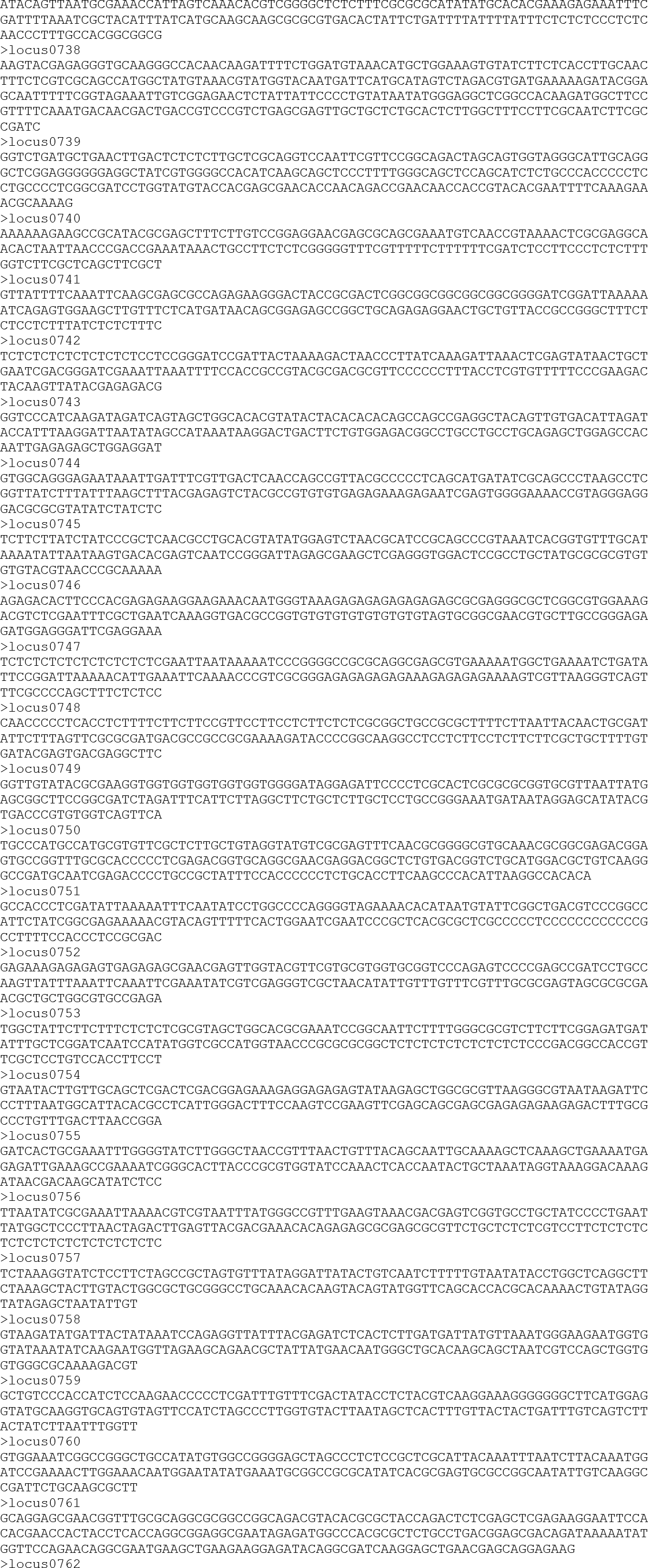

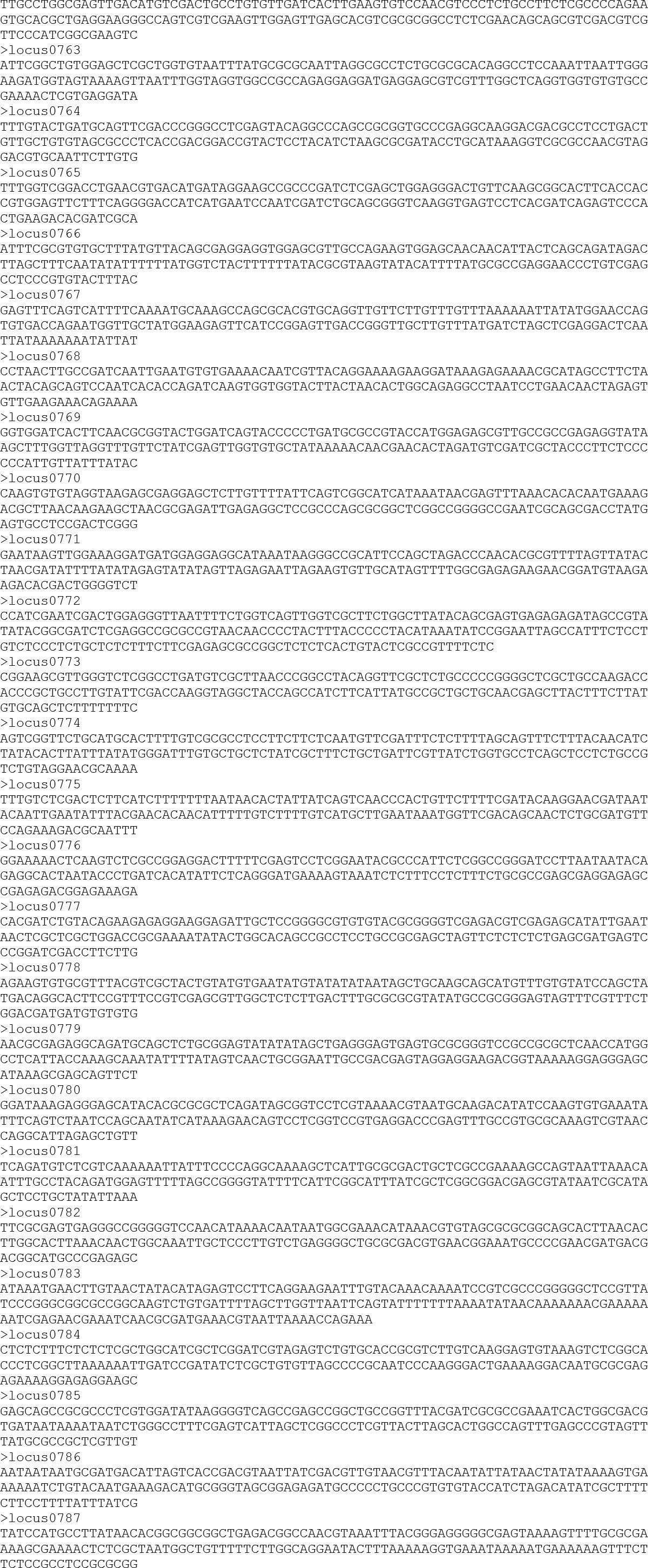

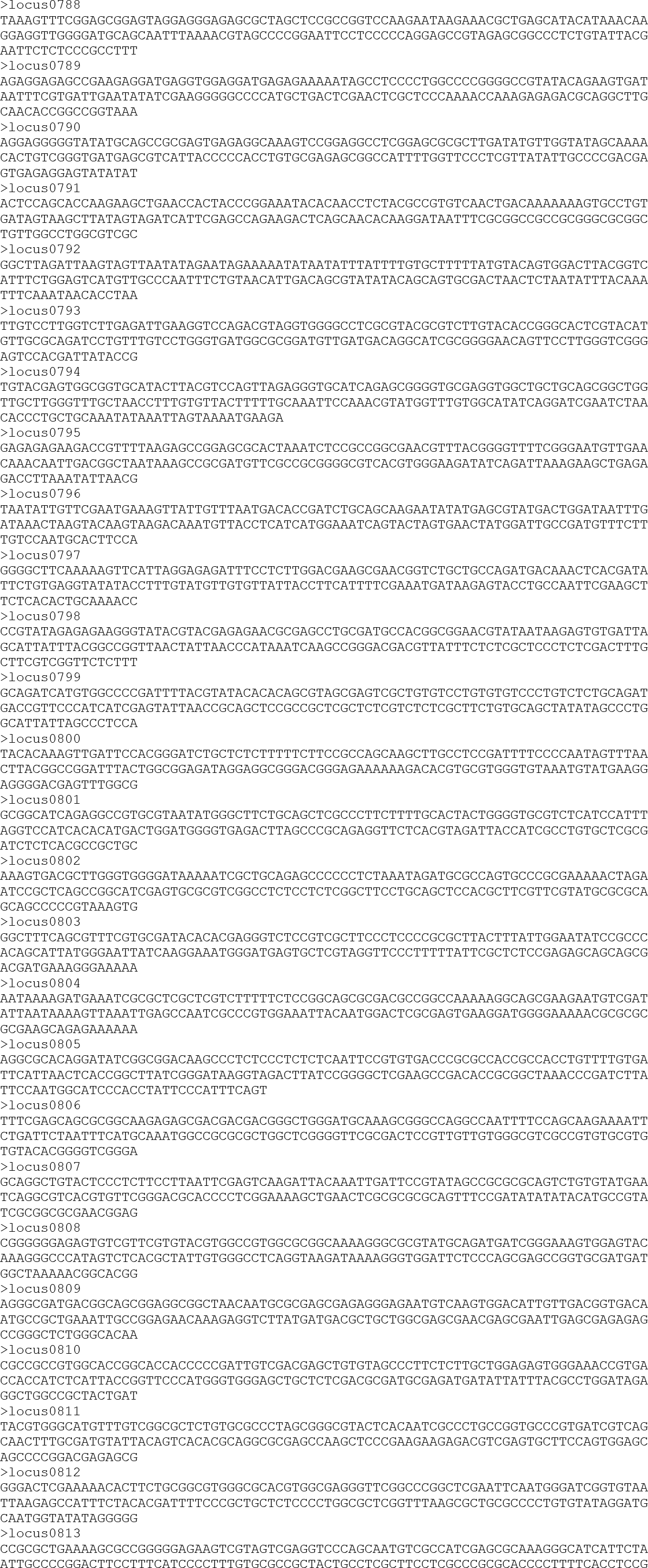

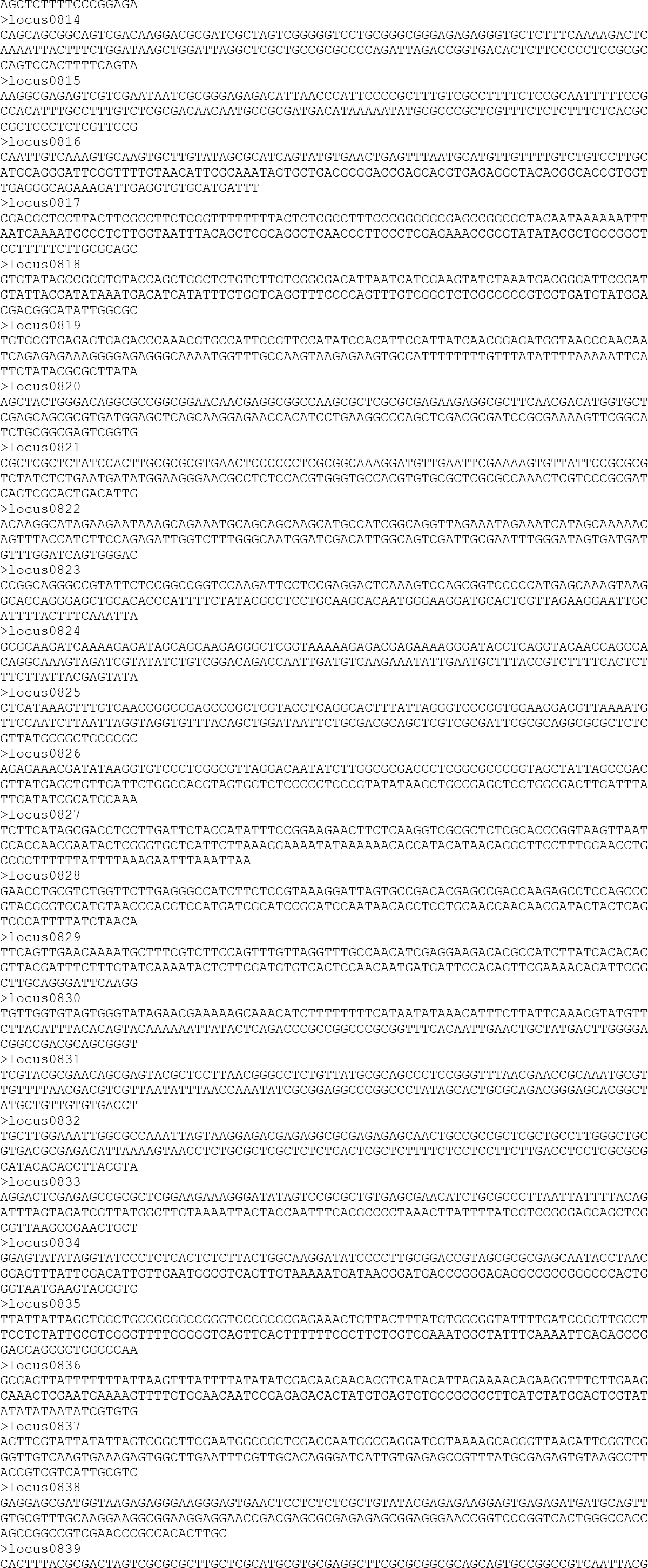

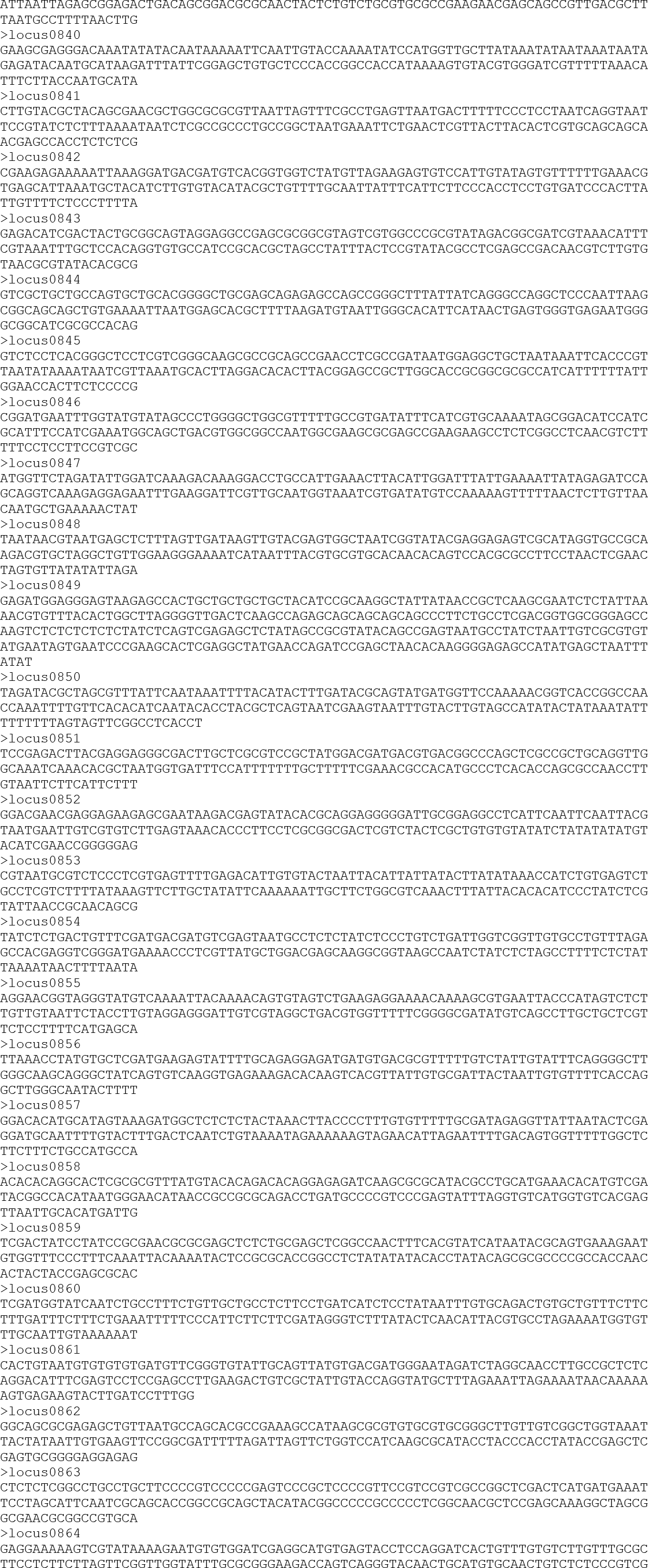

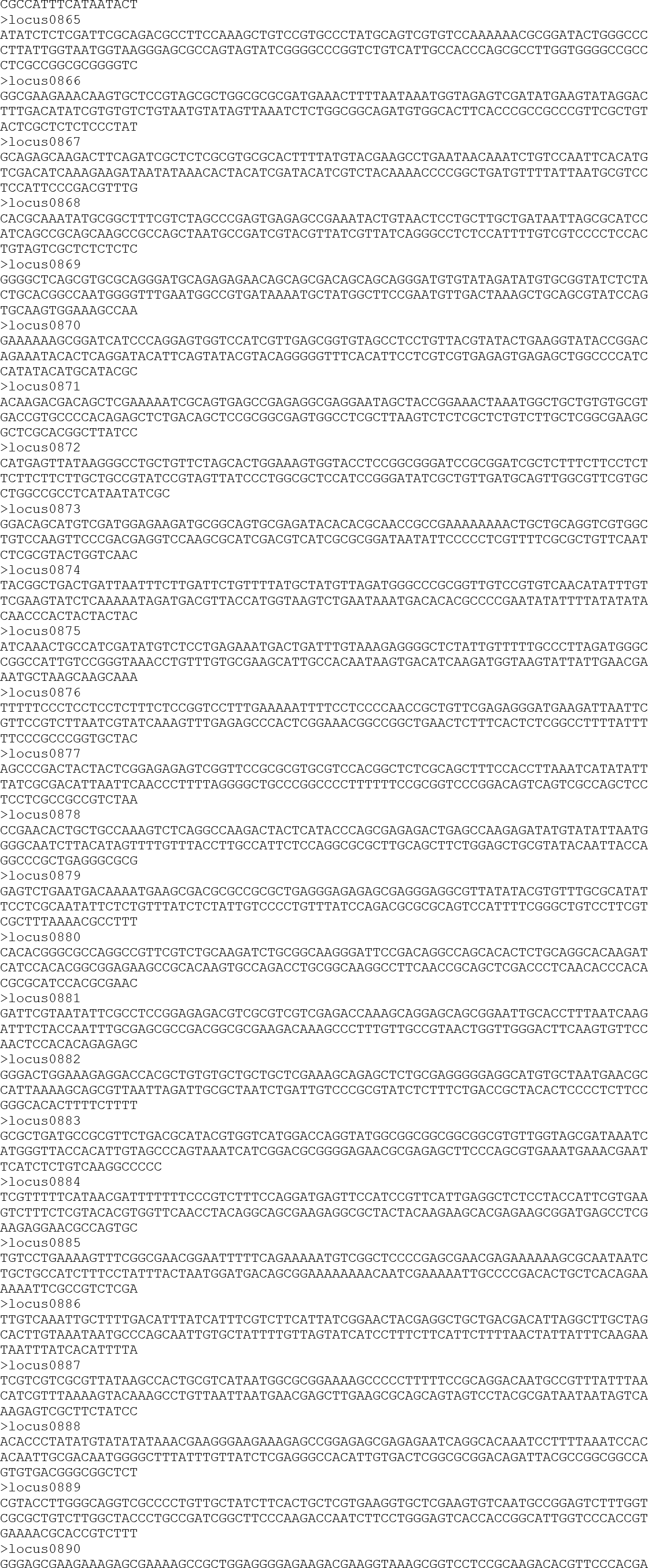

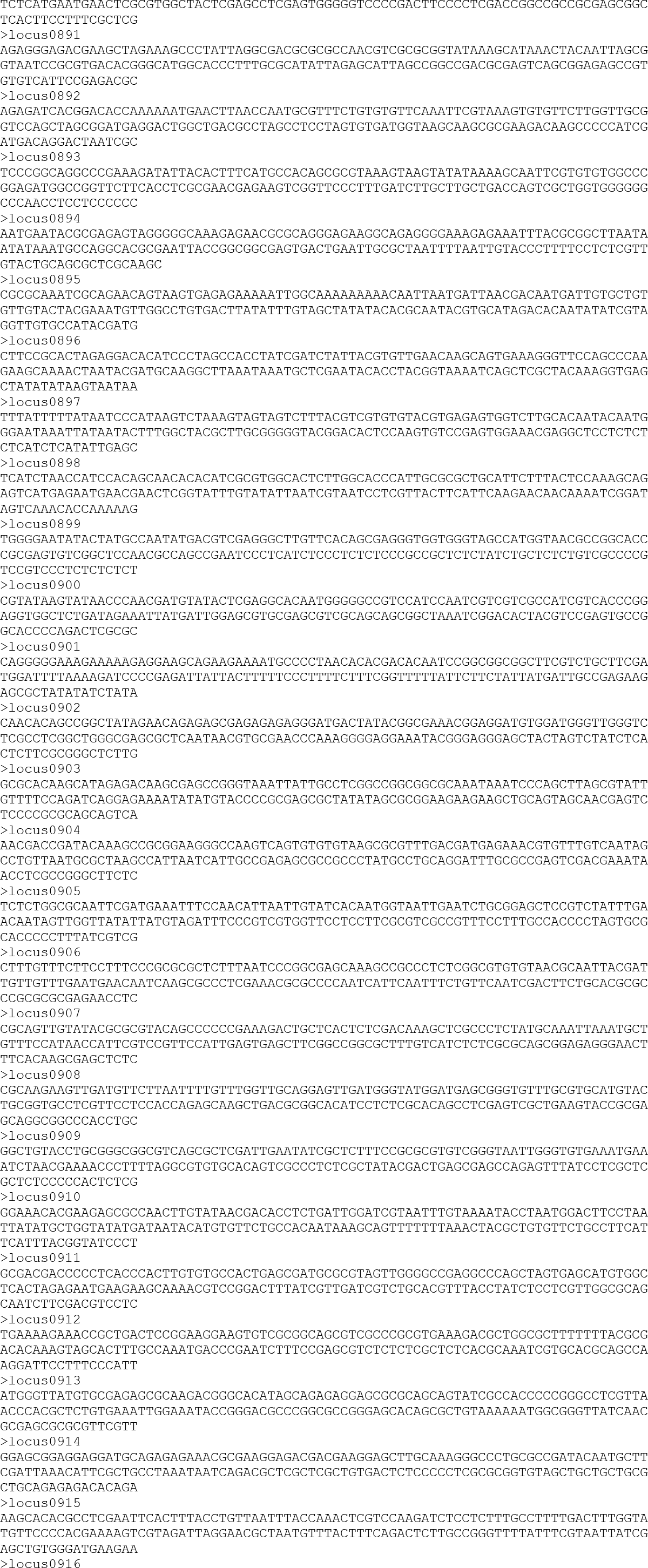

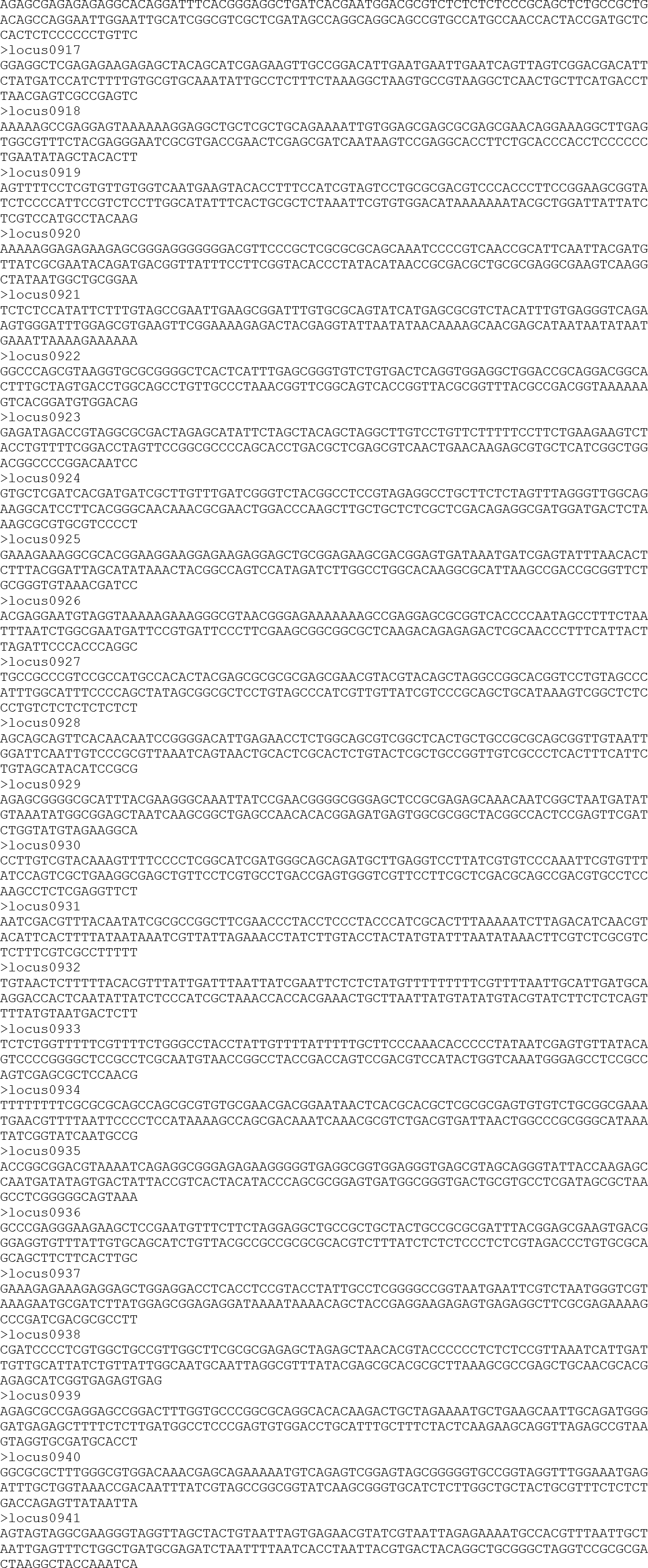

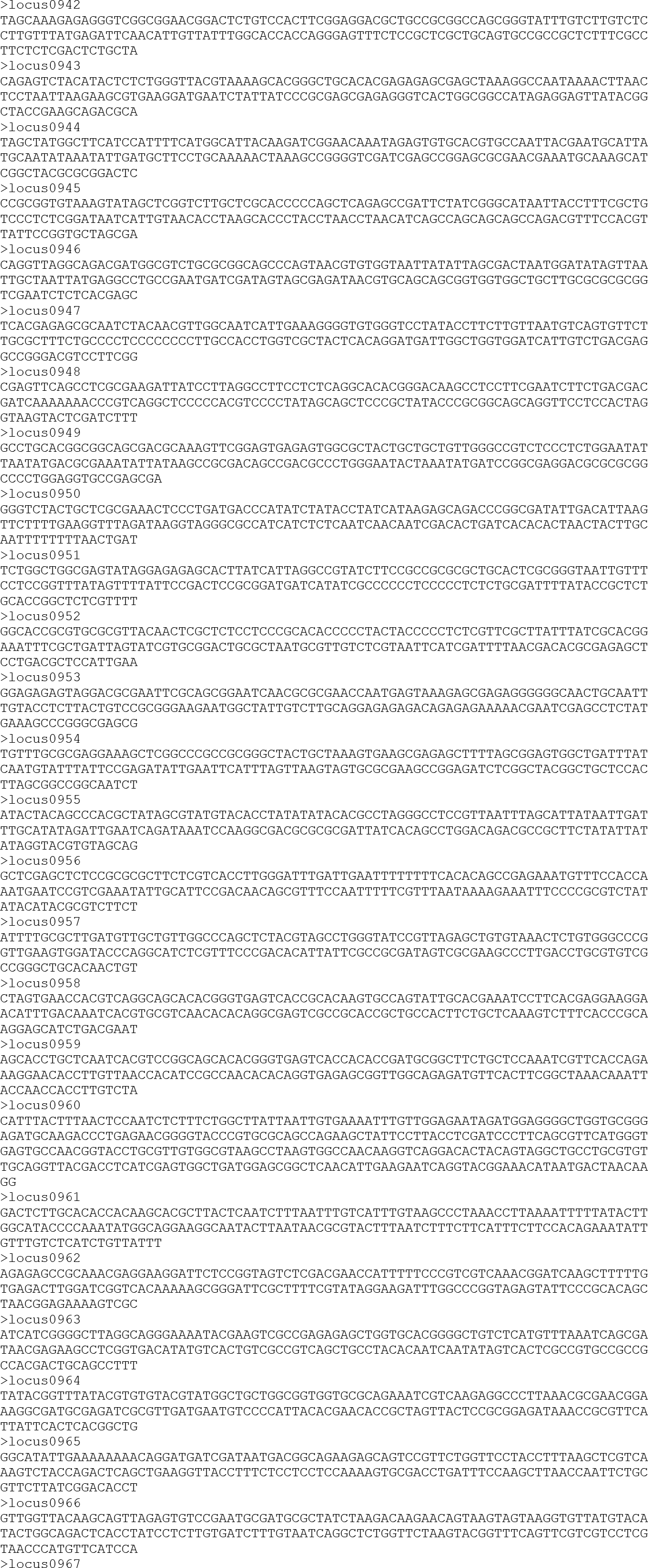

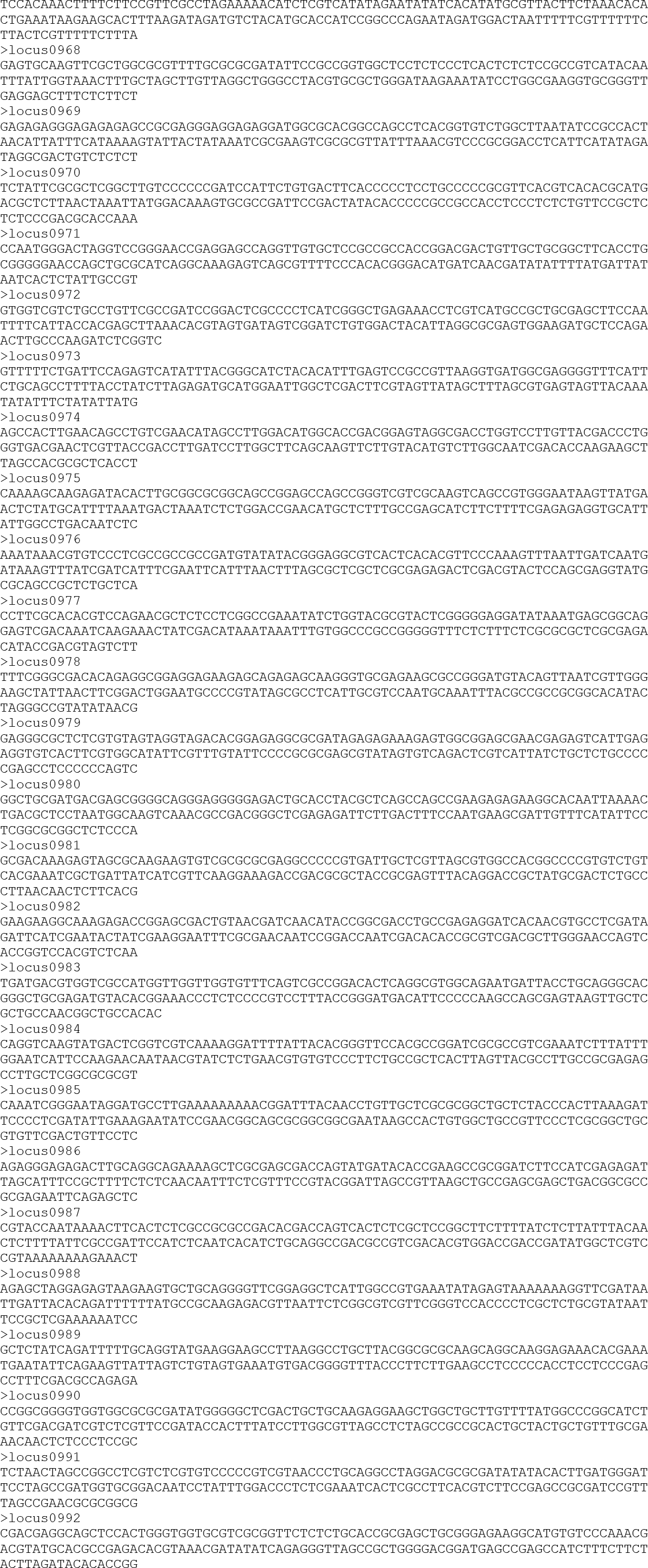

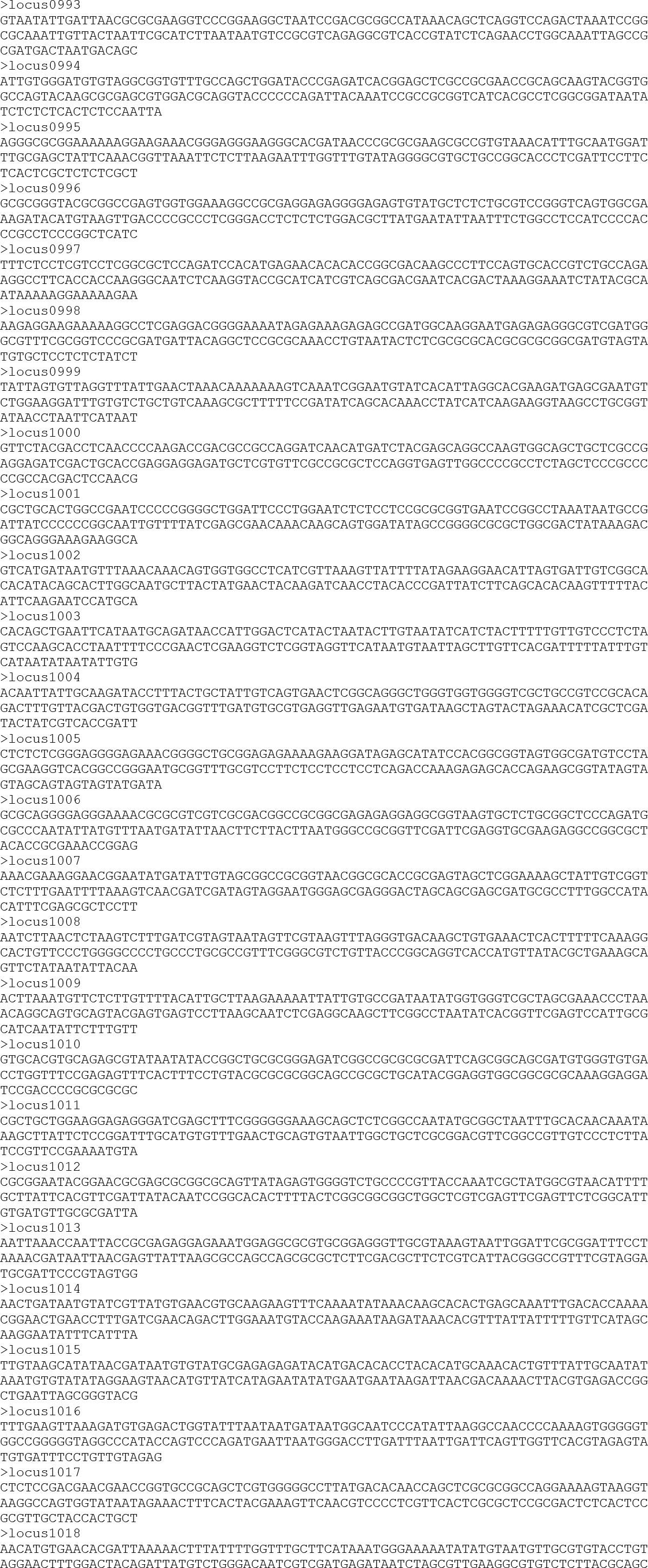

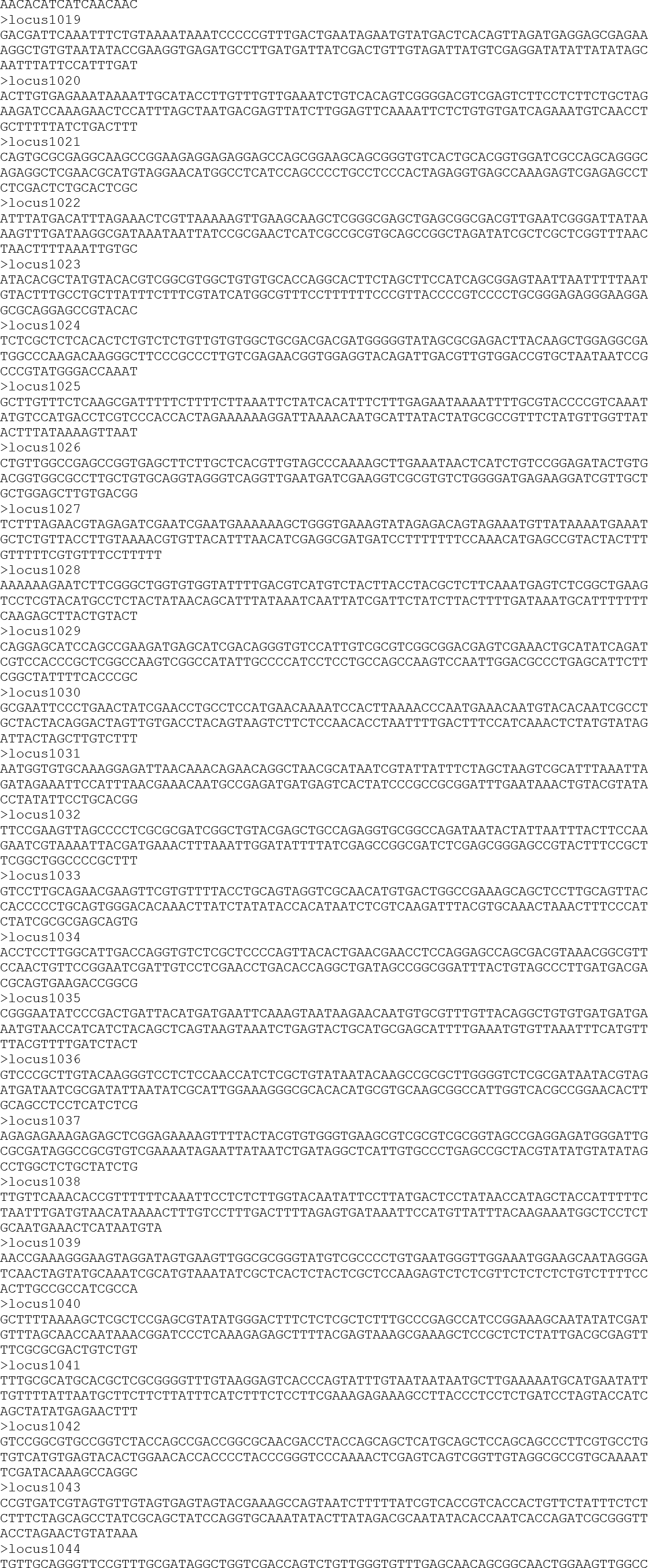

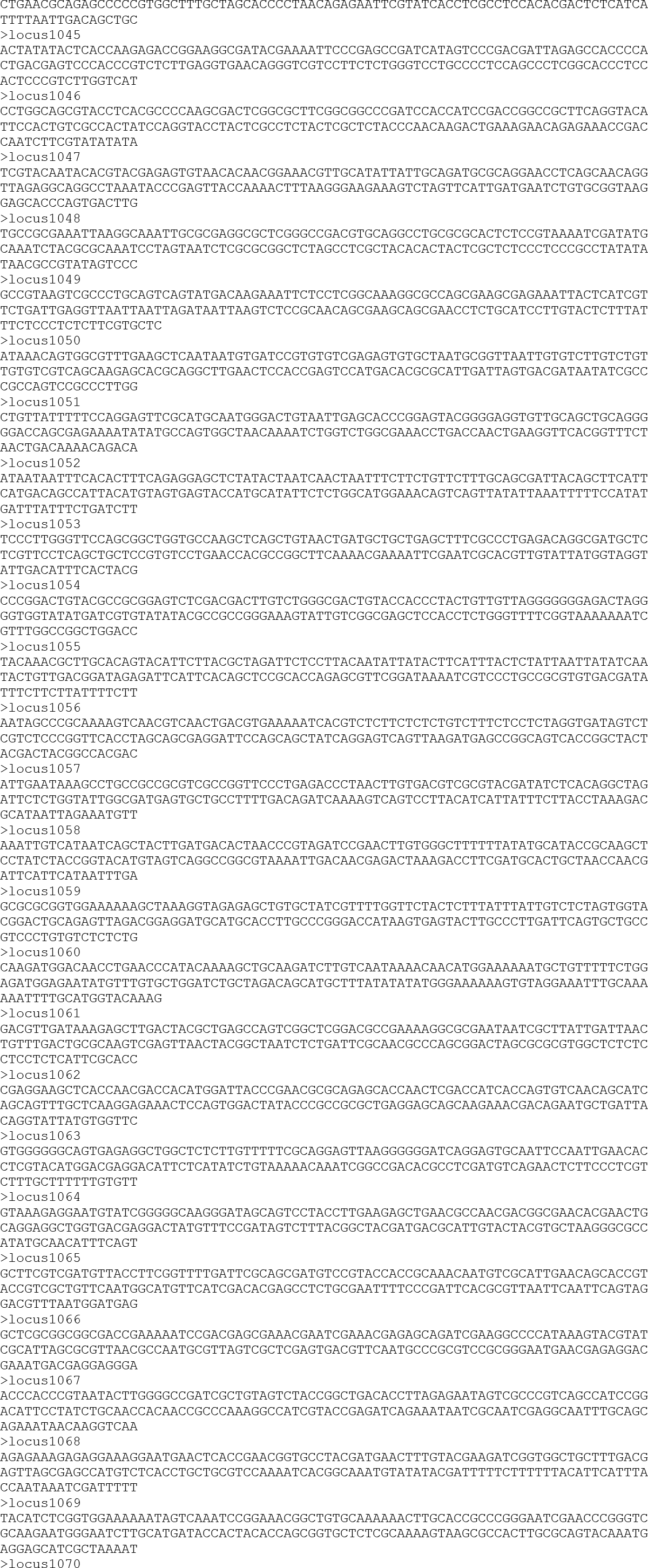

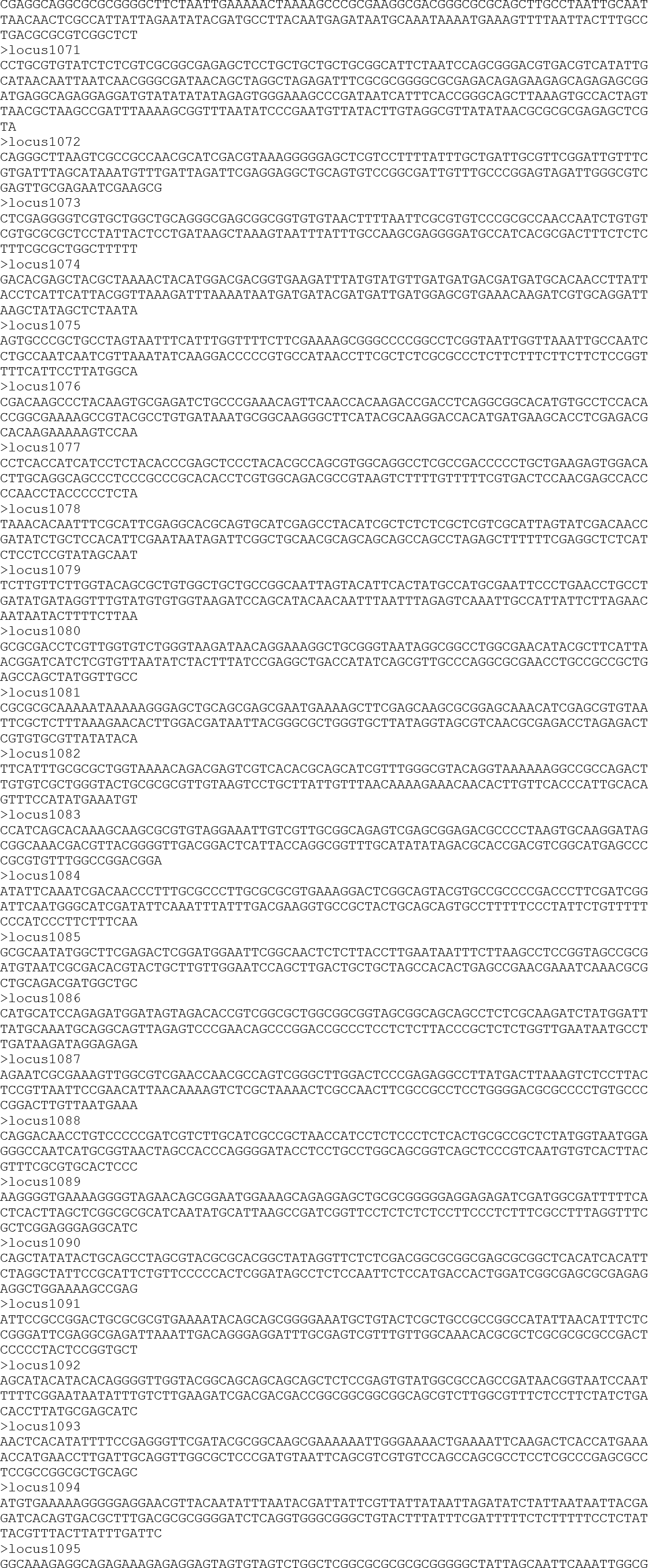

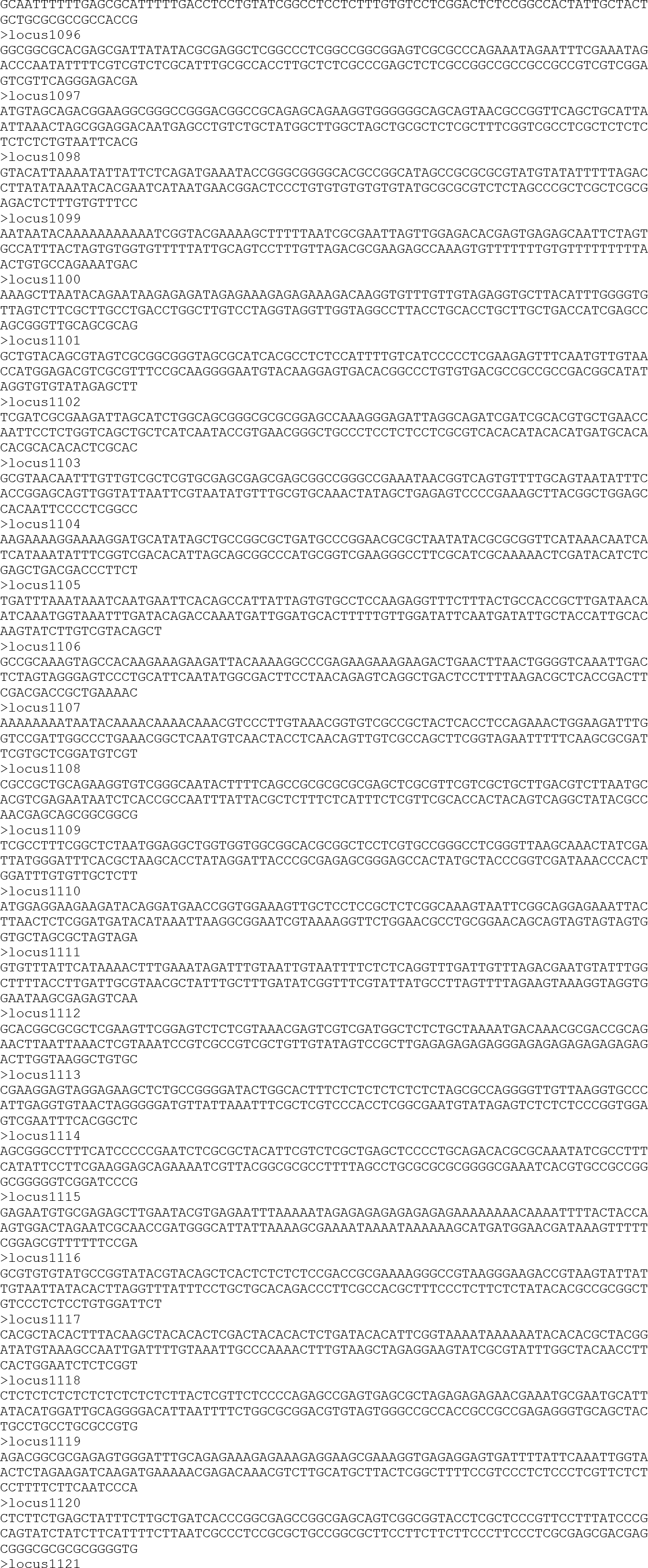

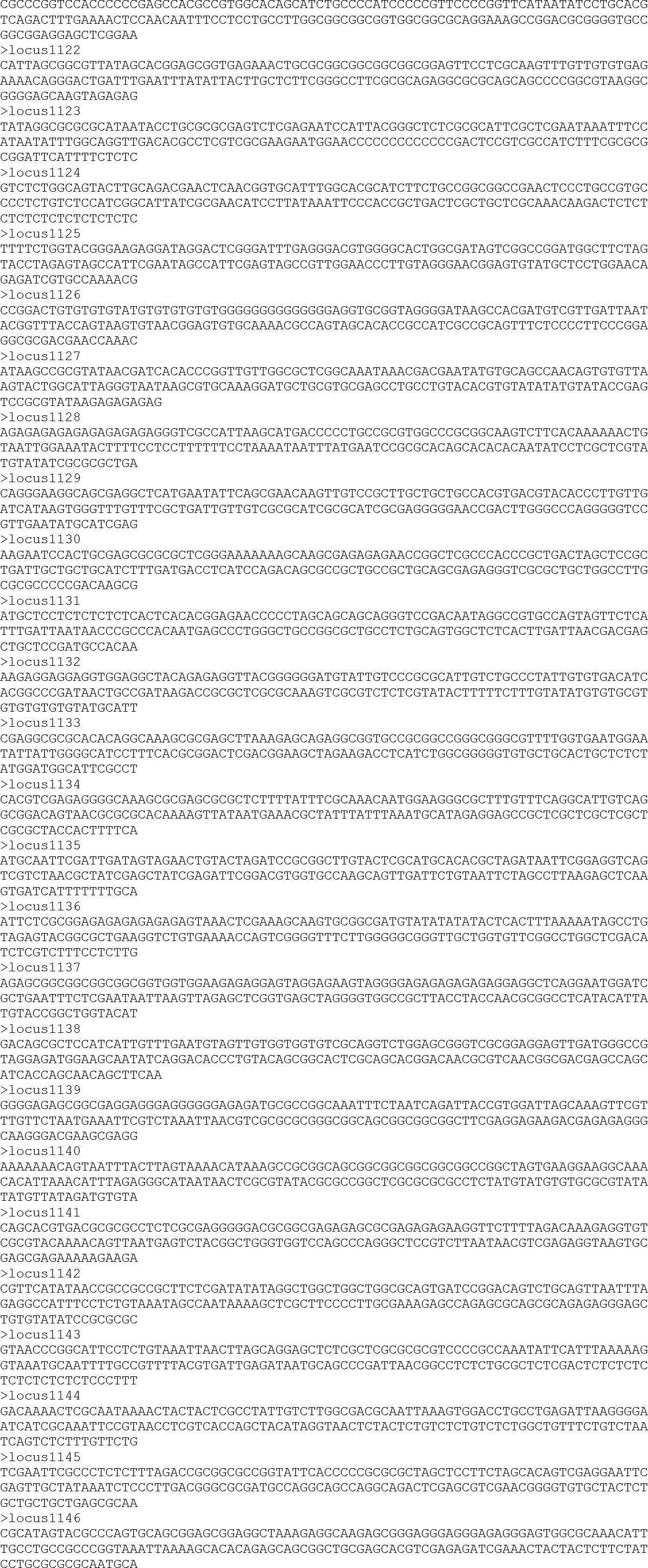

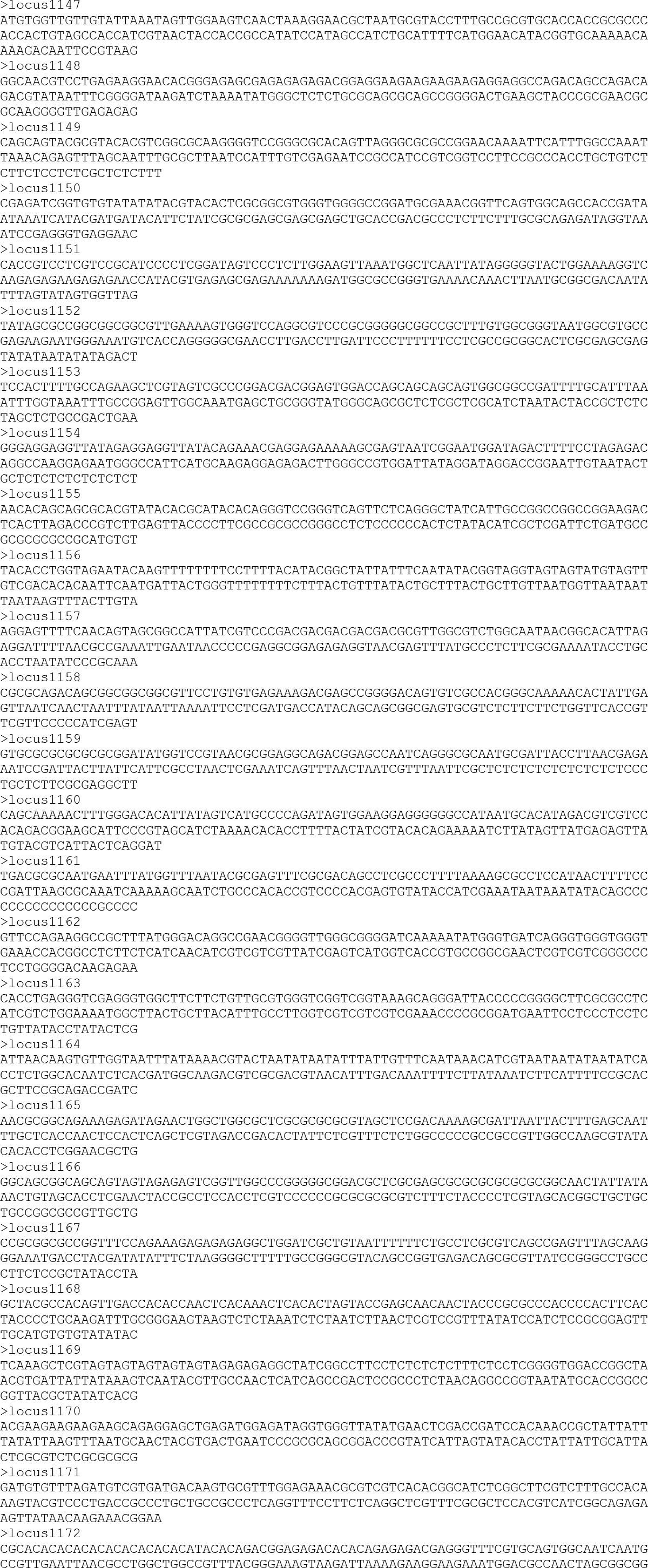

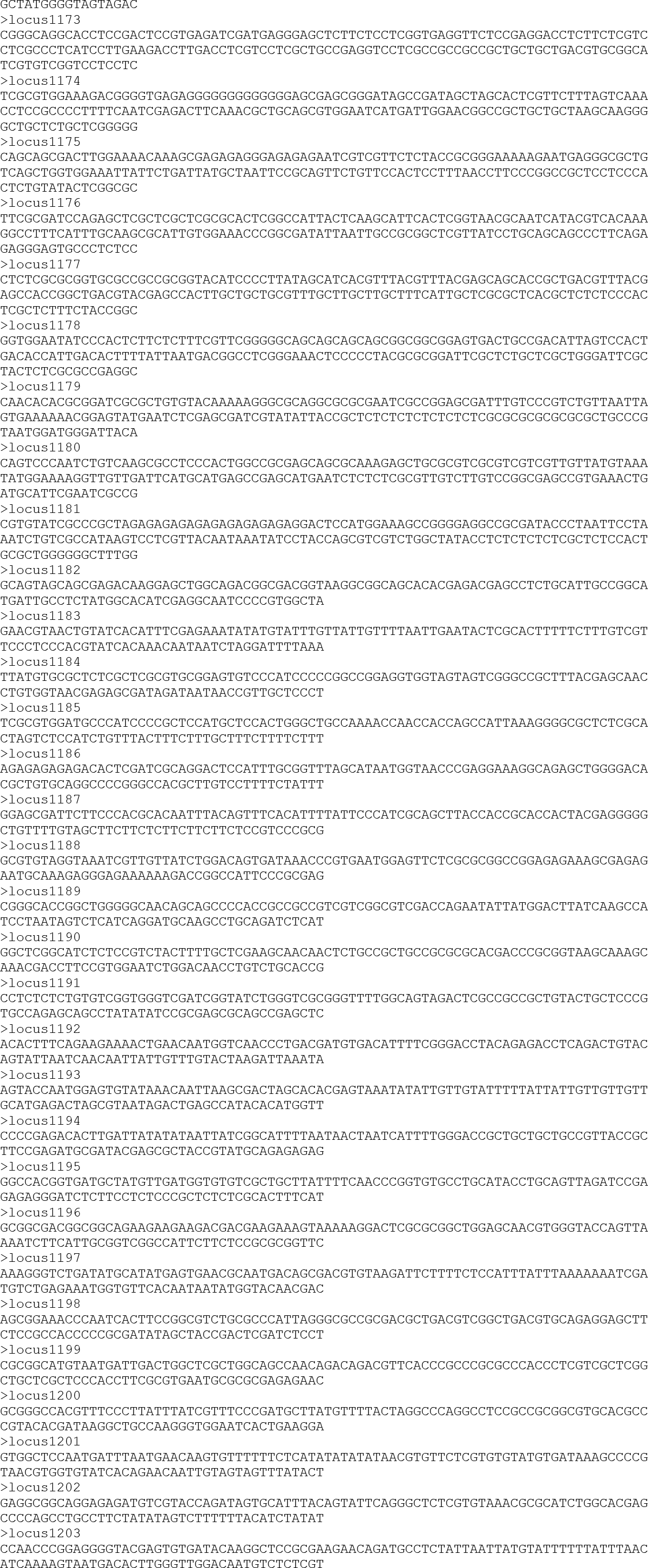

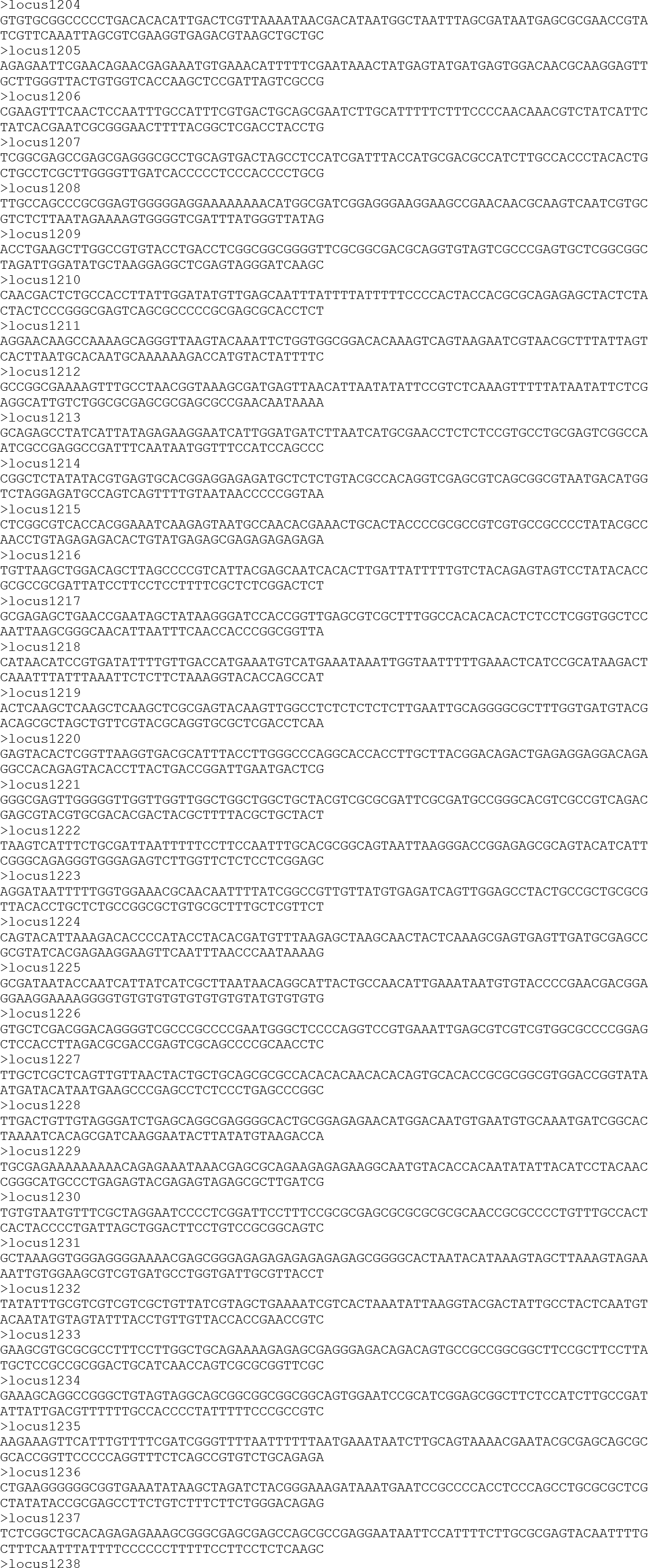

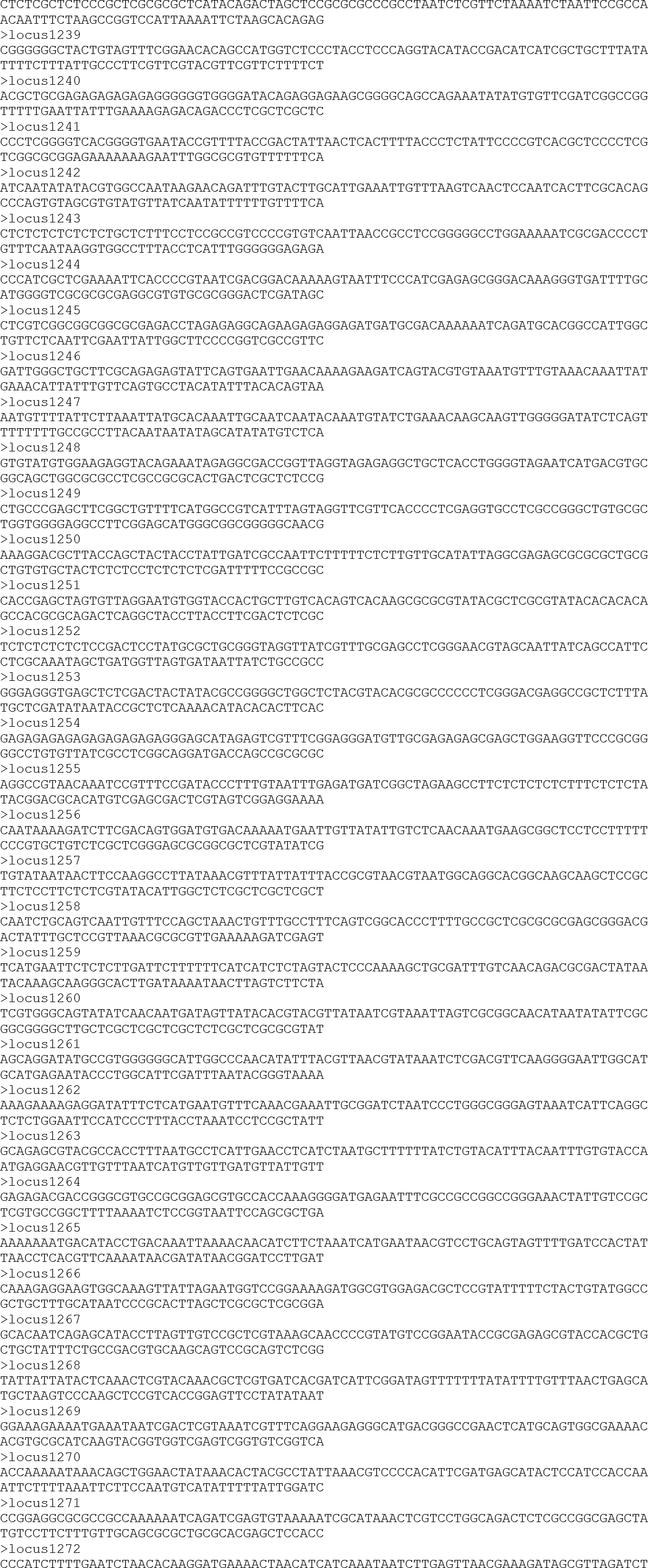

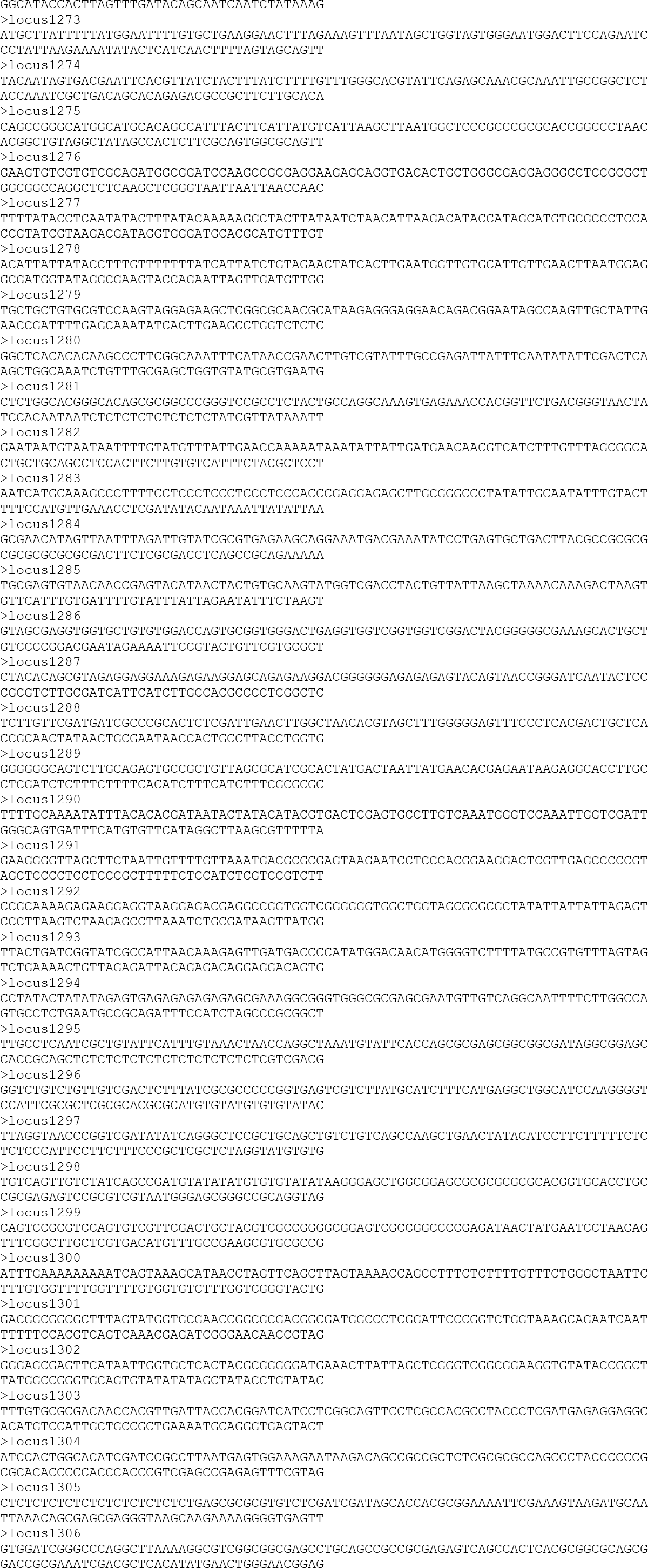

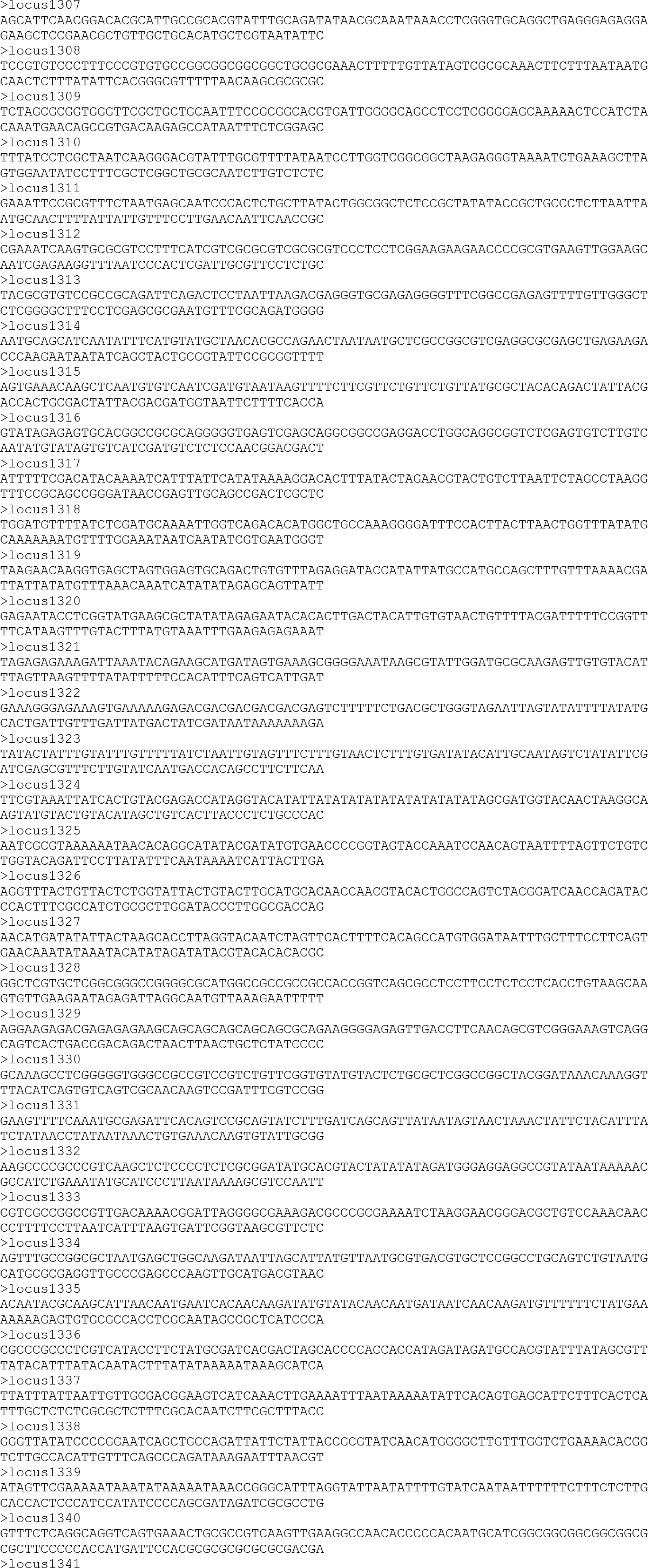

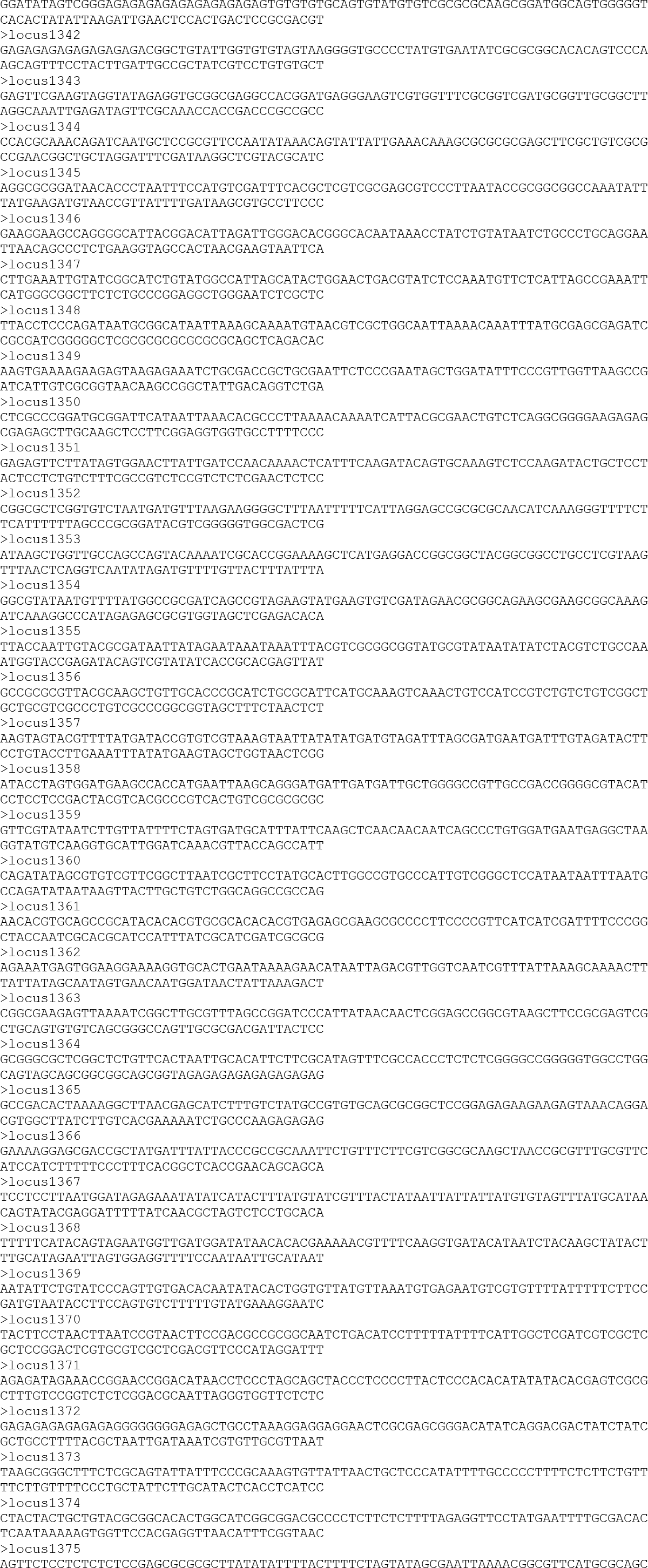

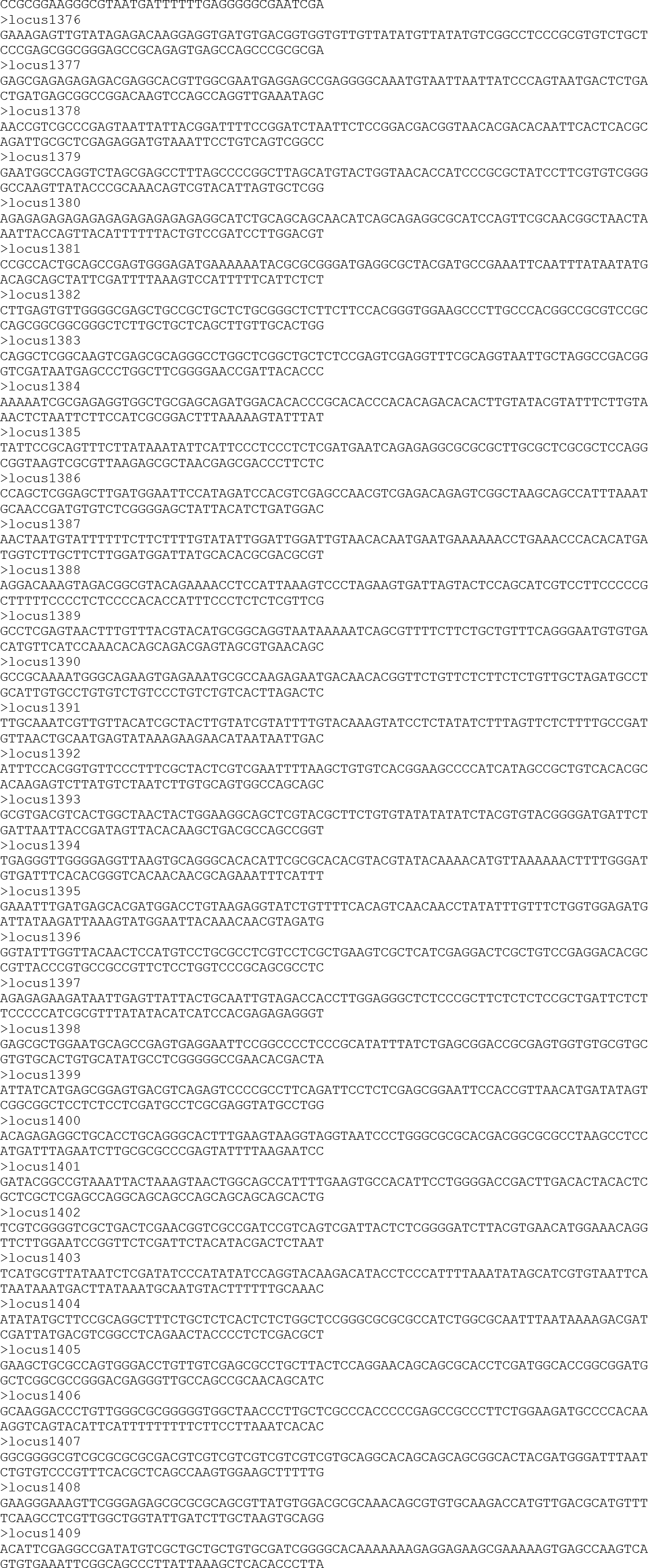

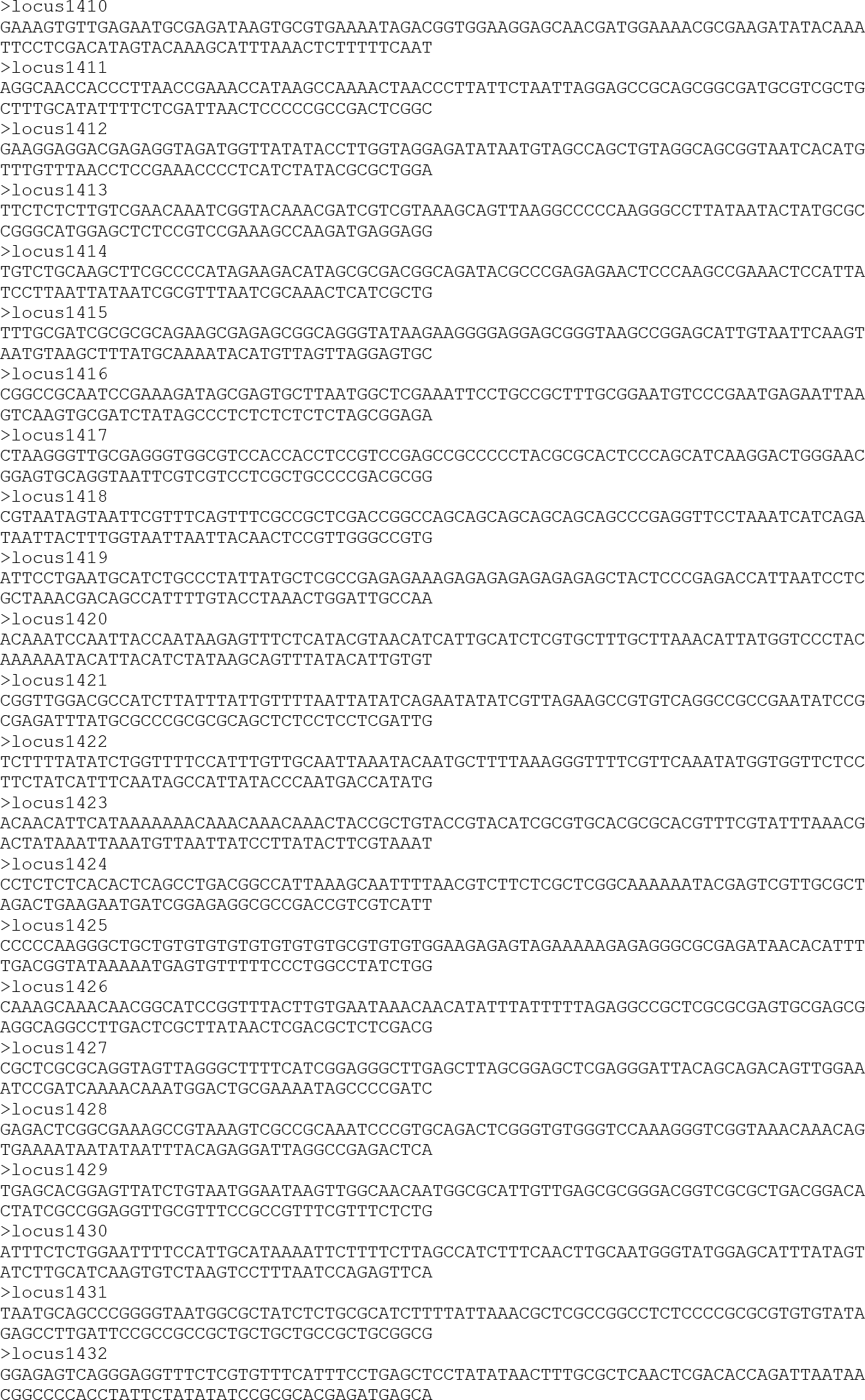

### Additional file 6

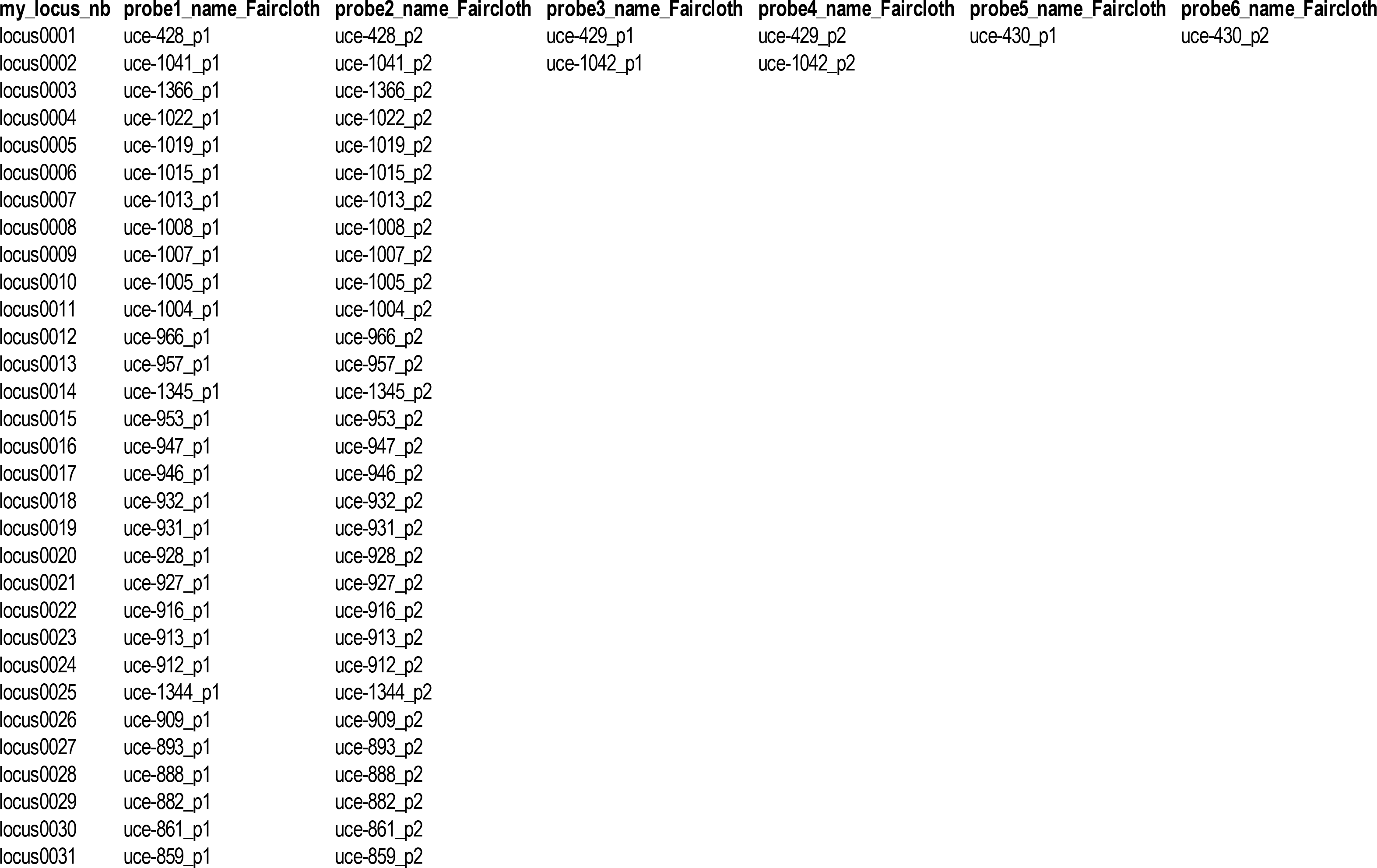

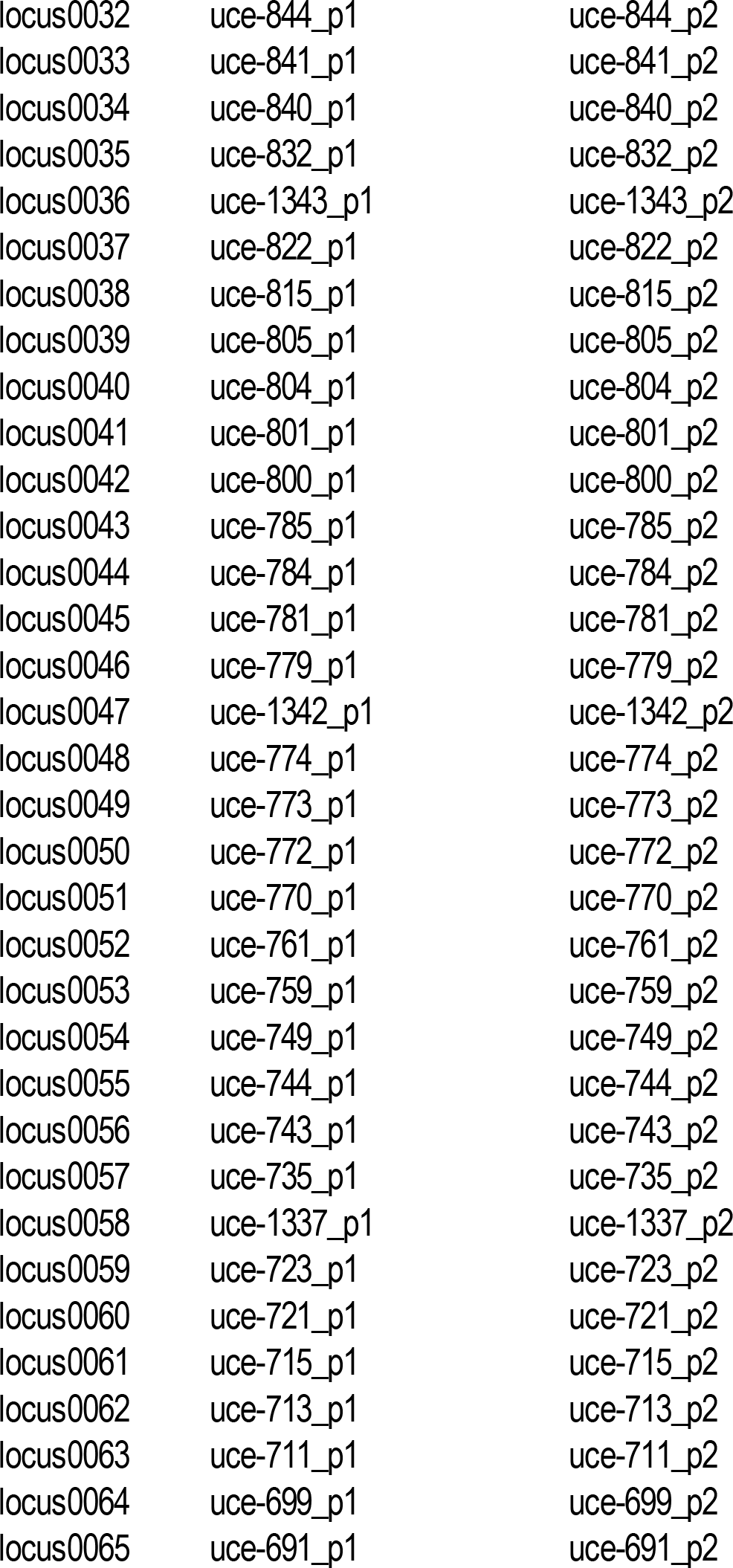

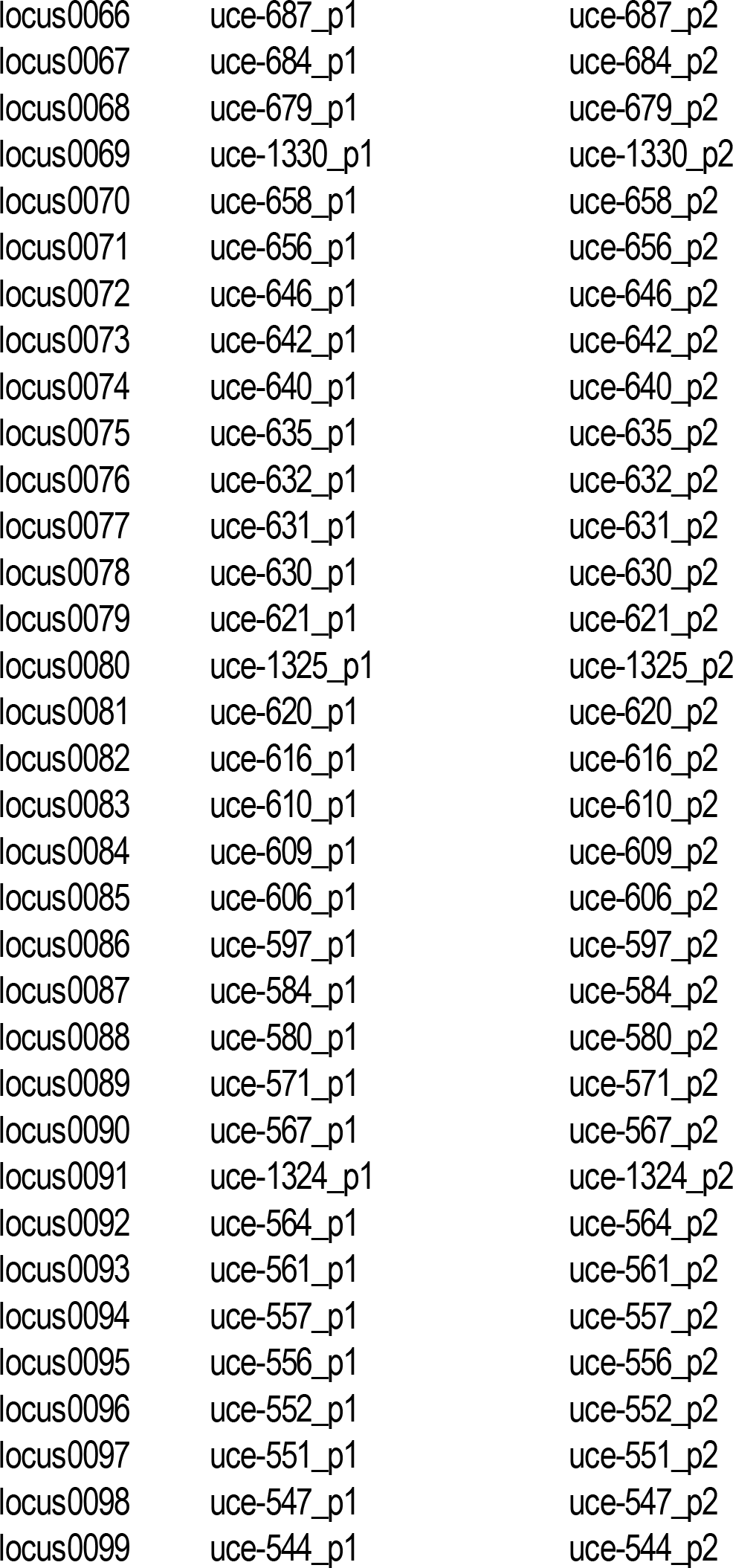

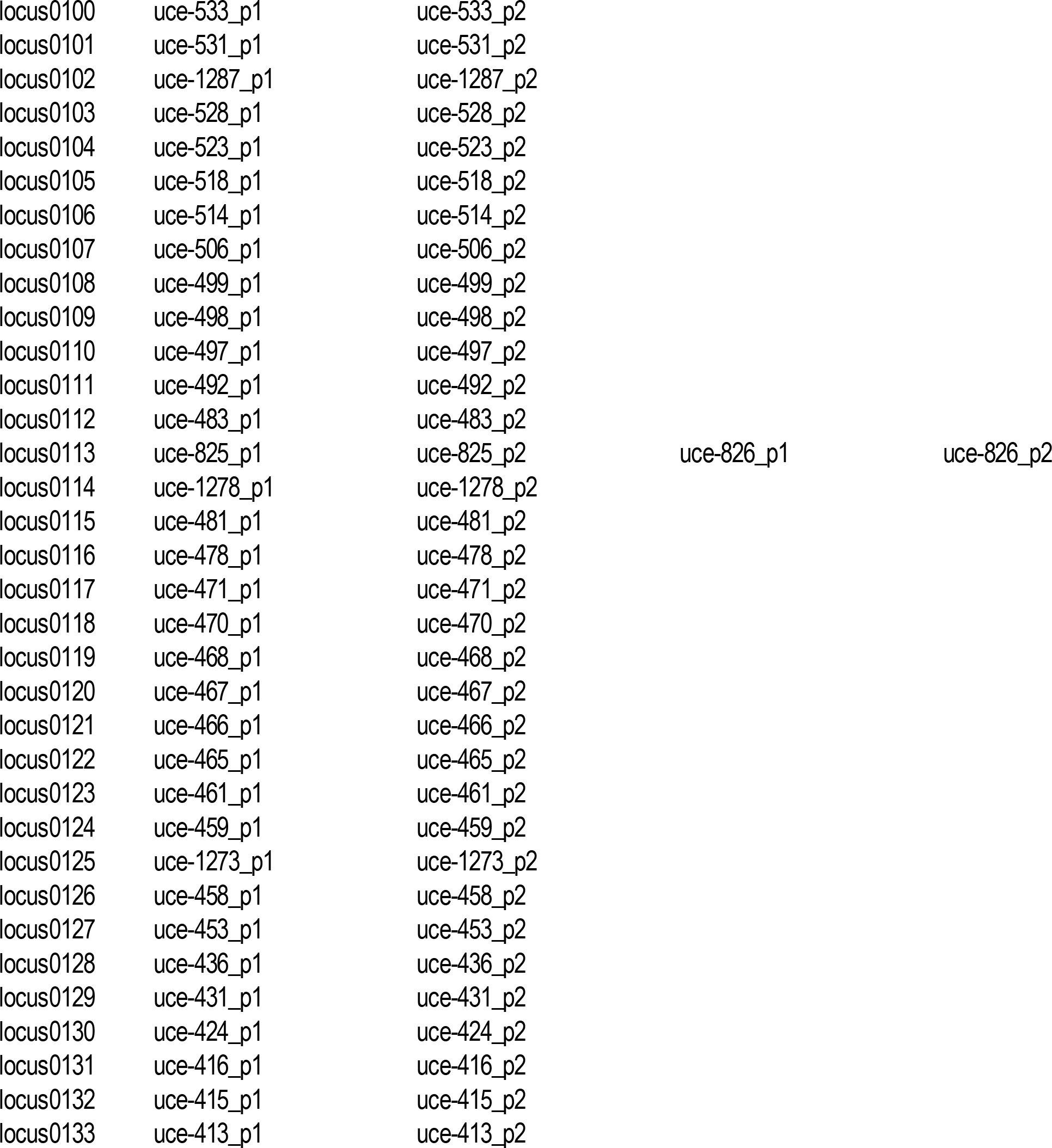

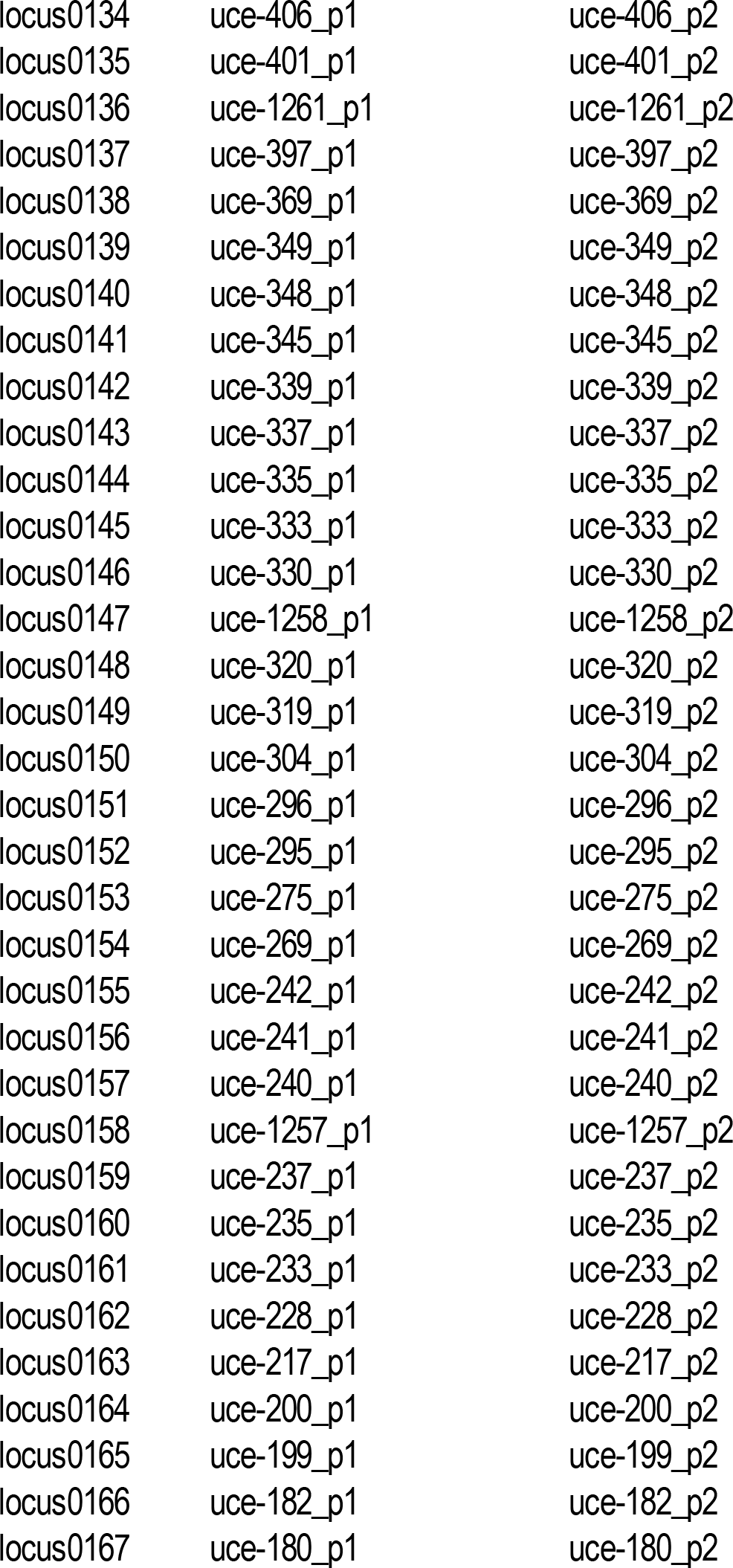

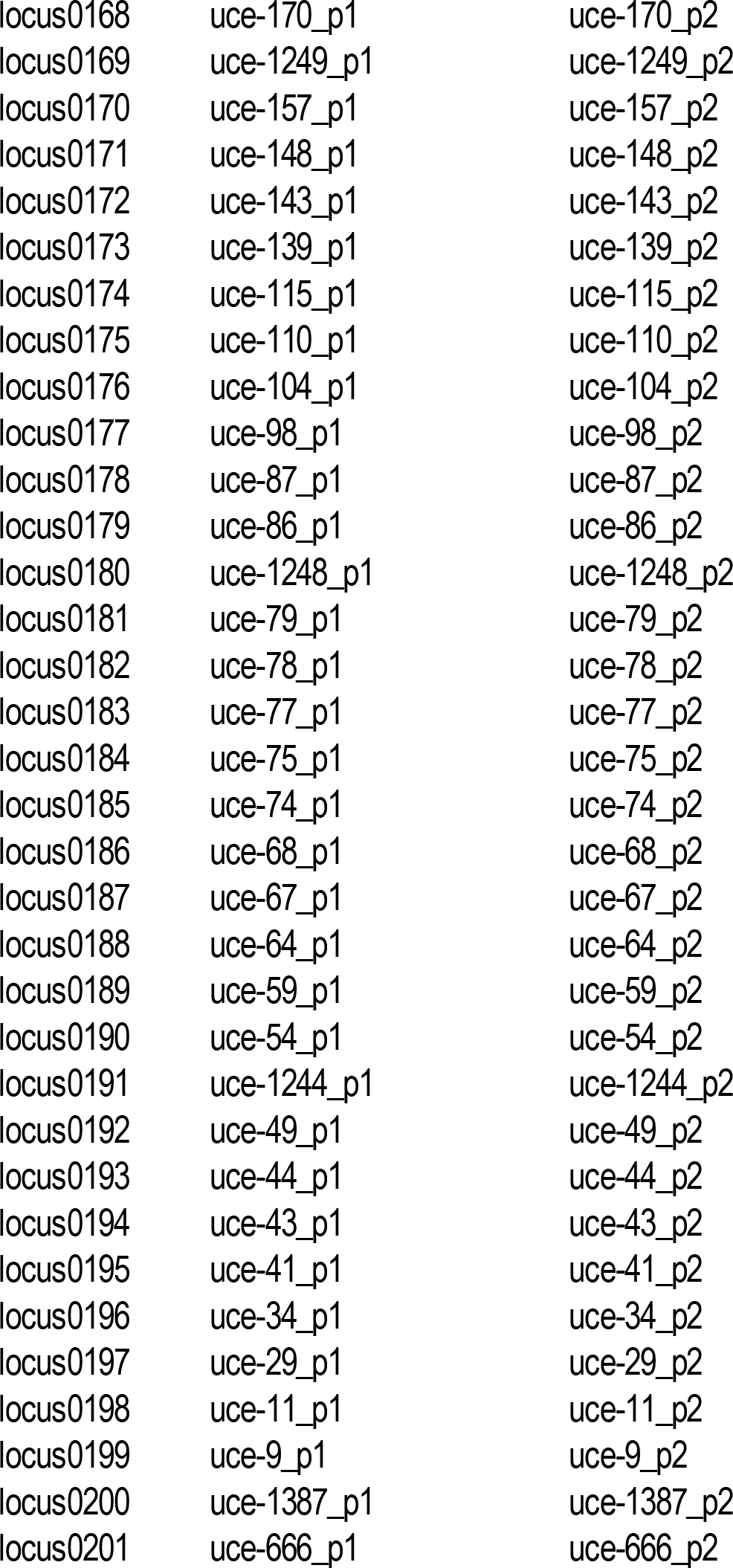

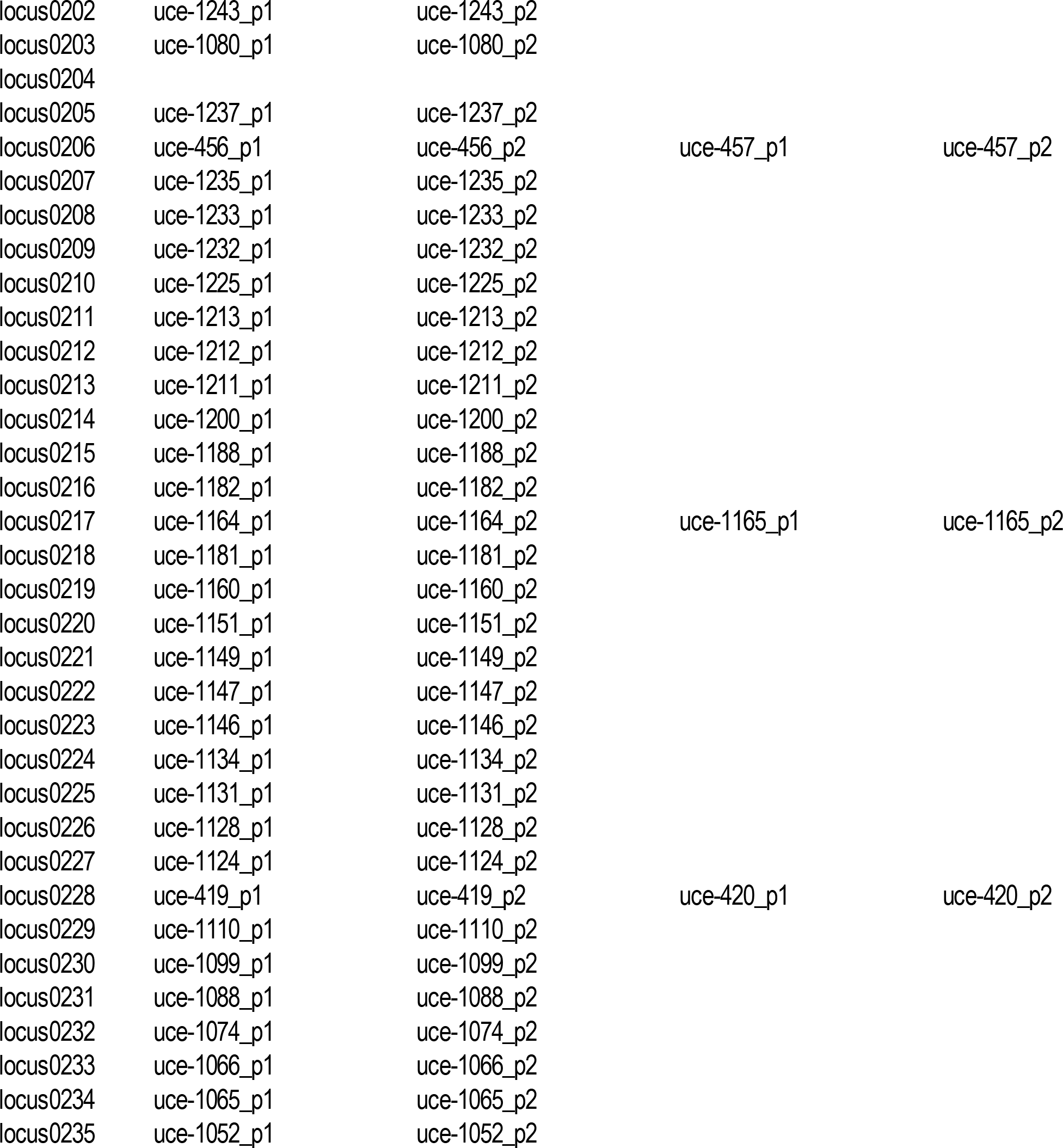

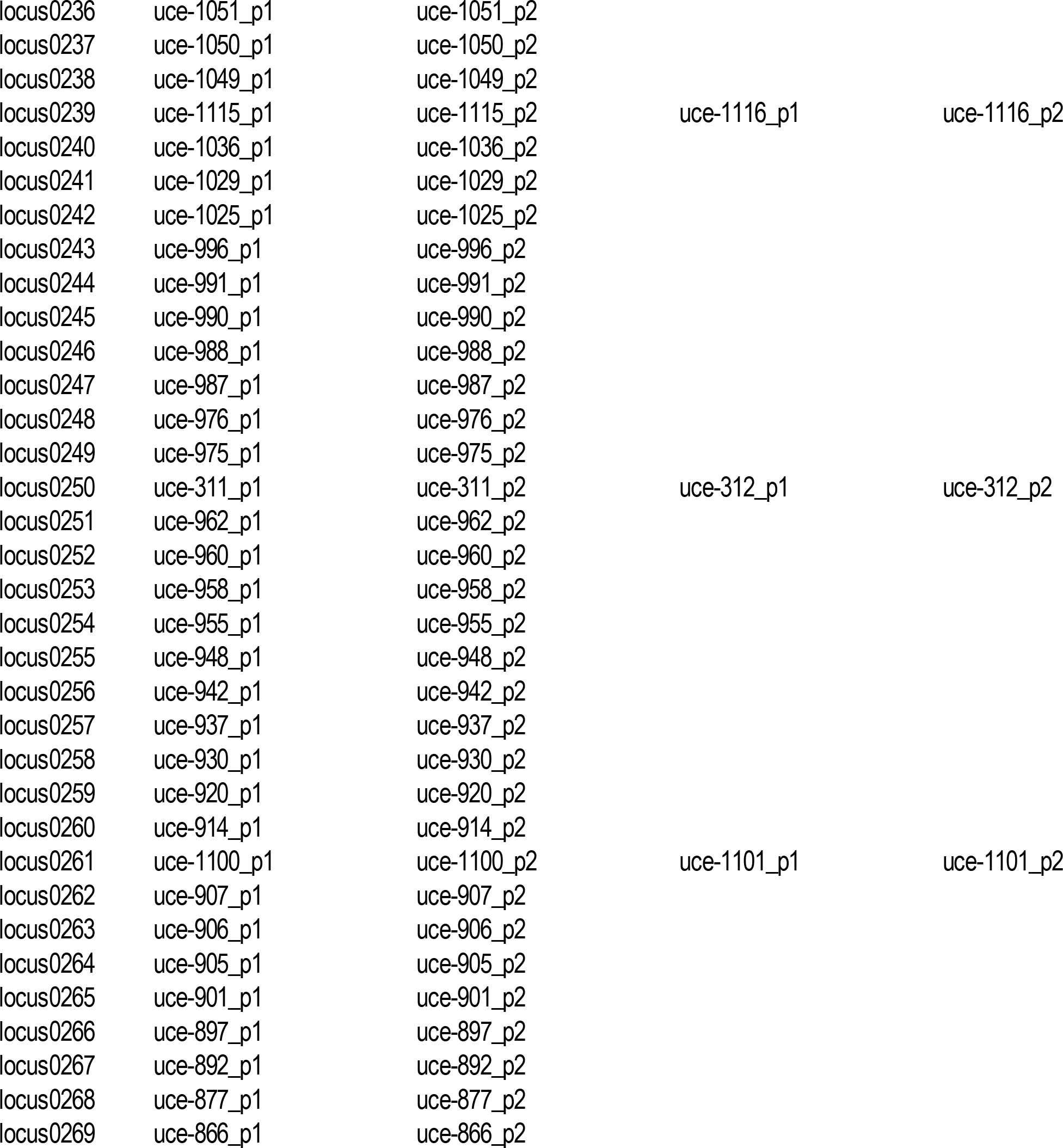

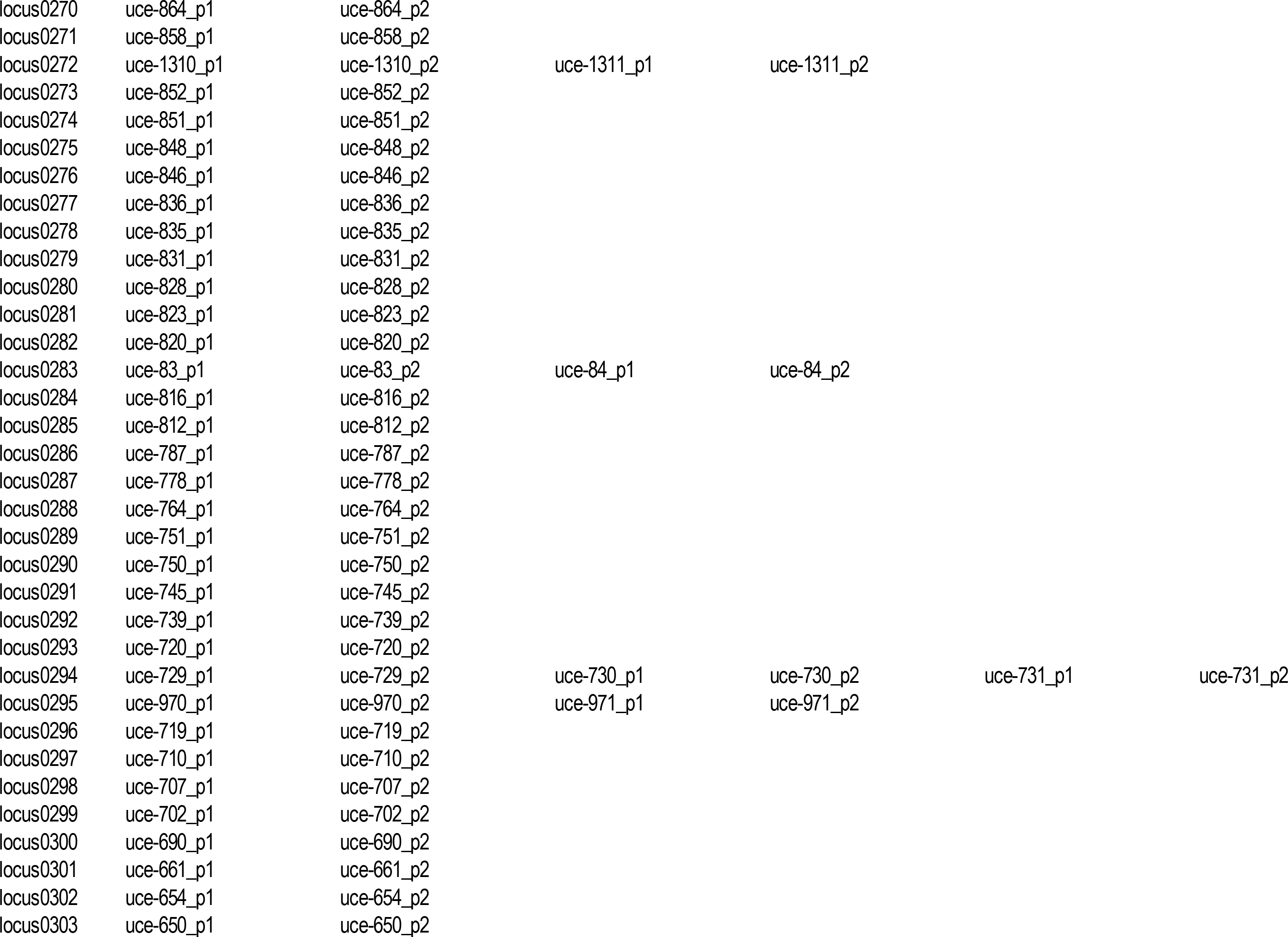

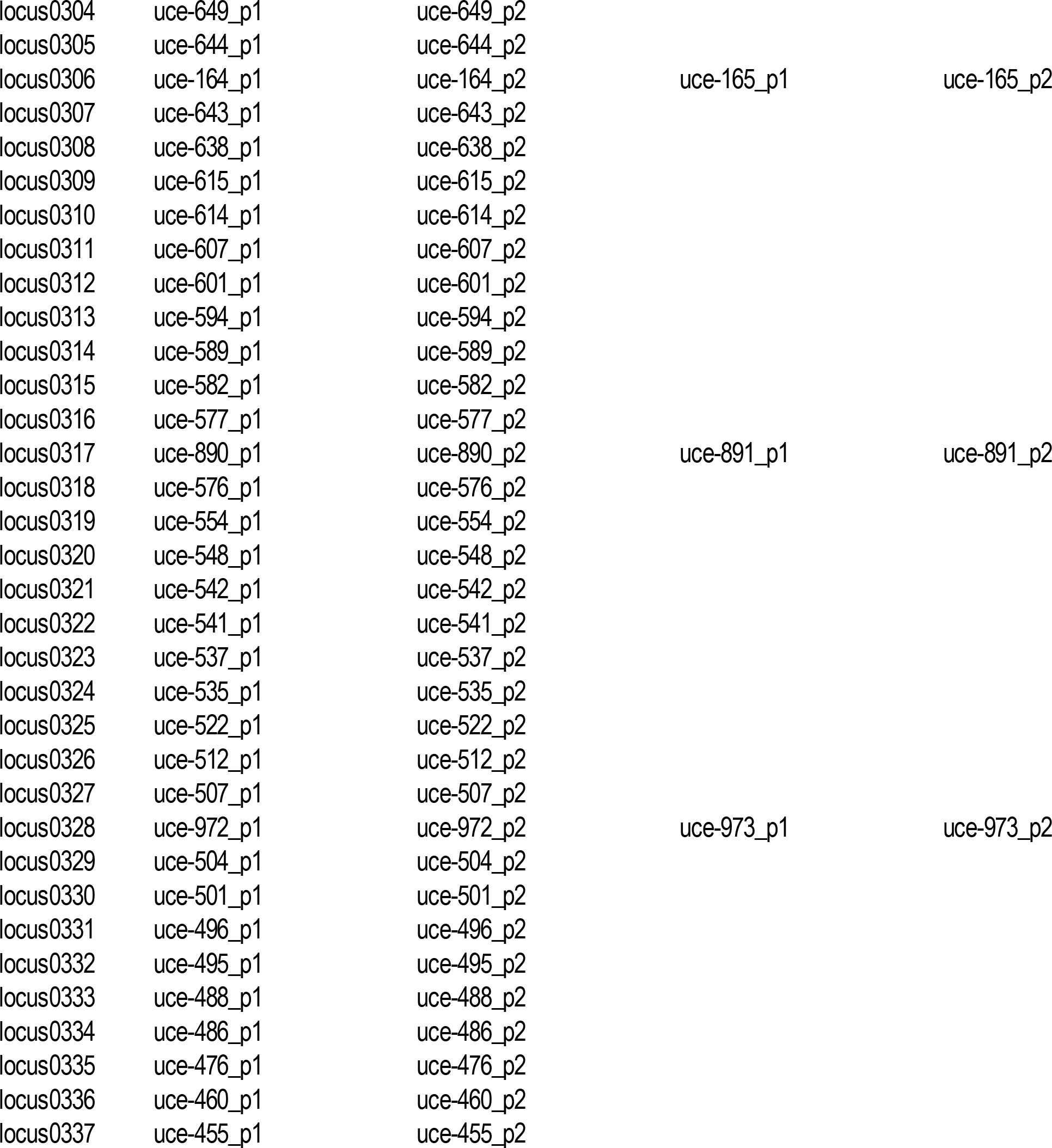

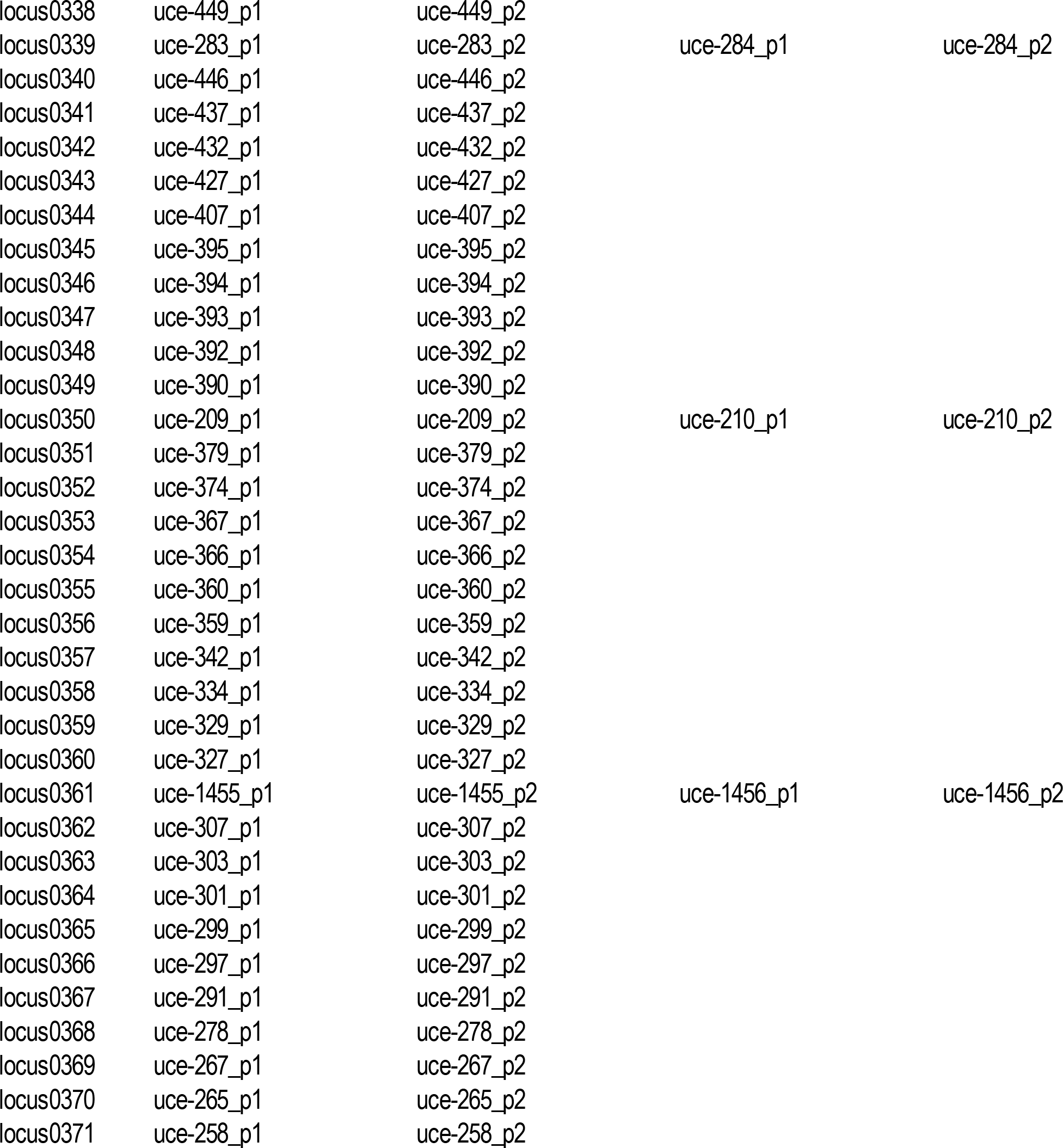

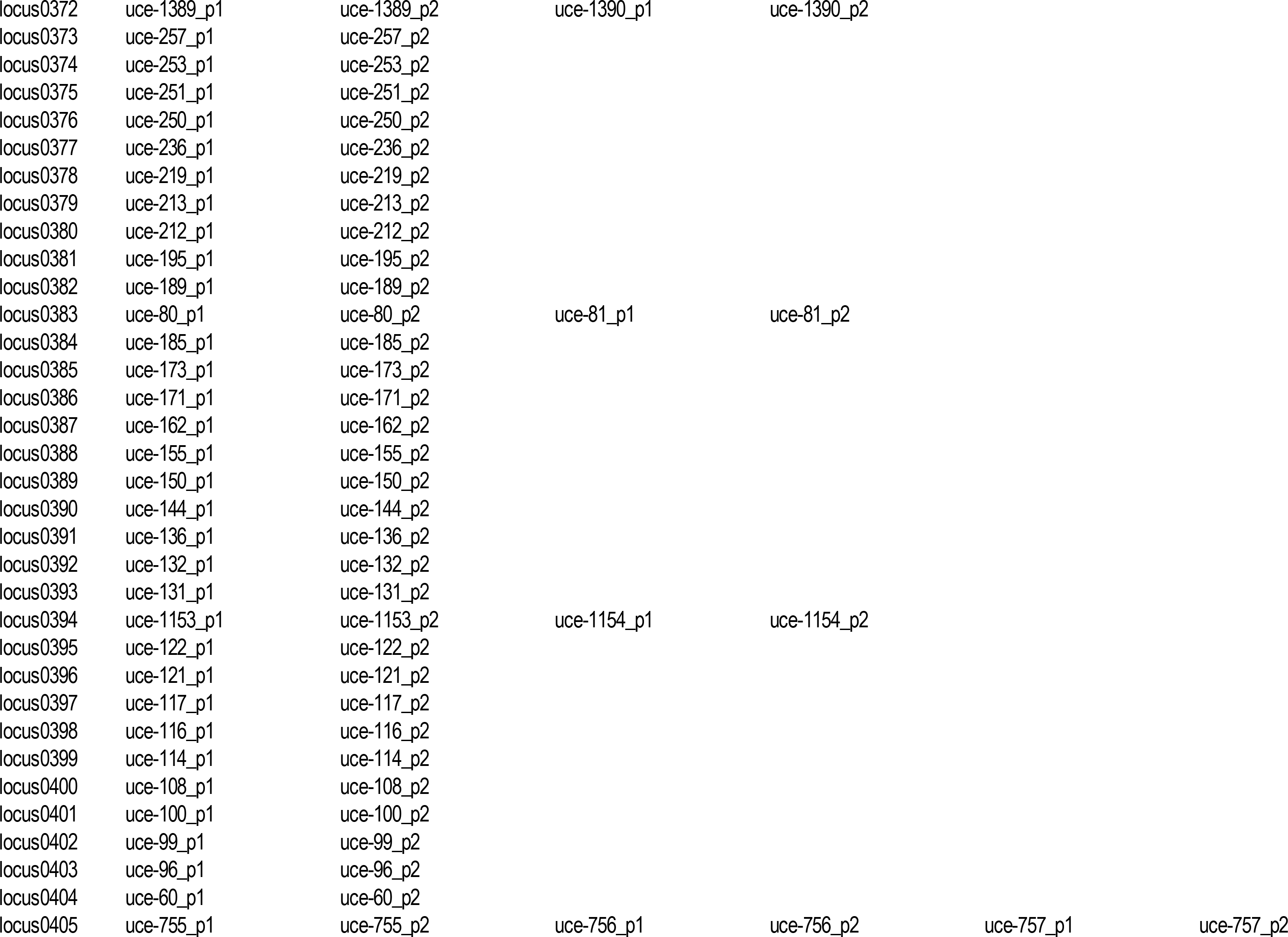

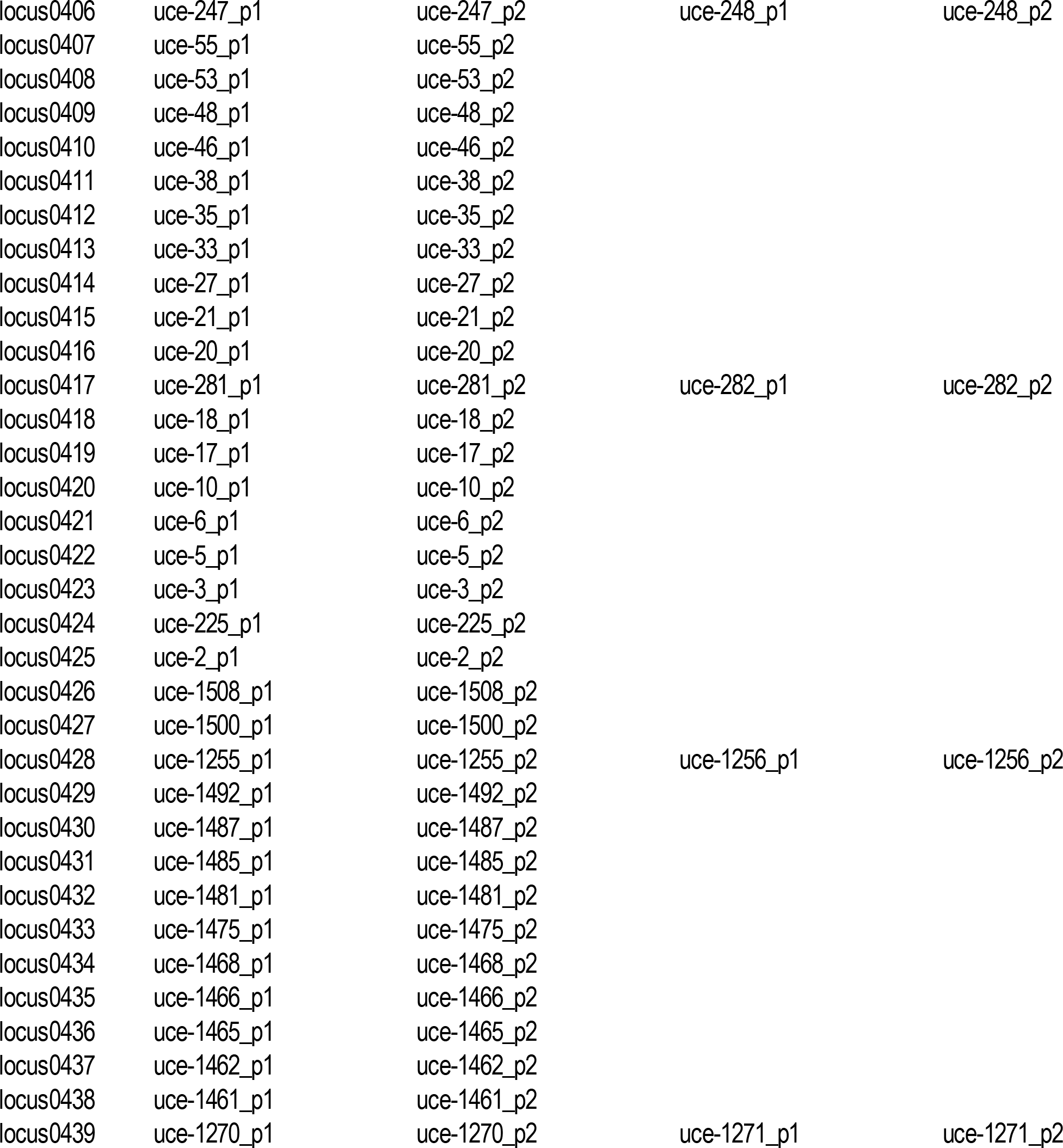

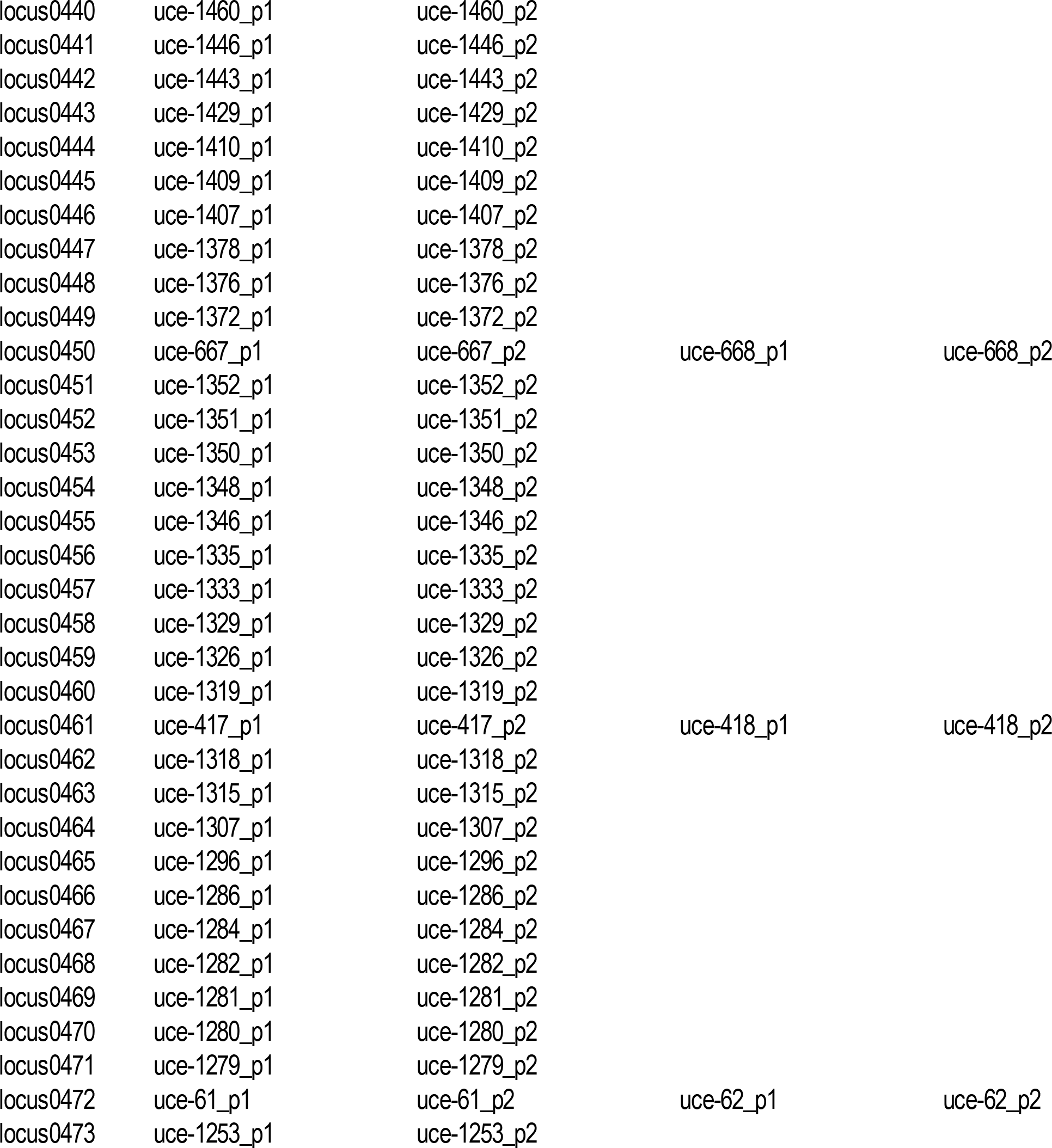

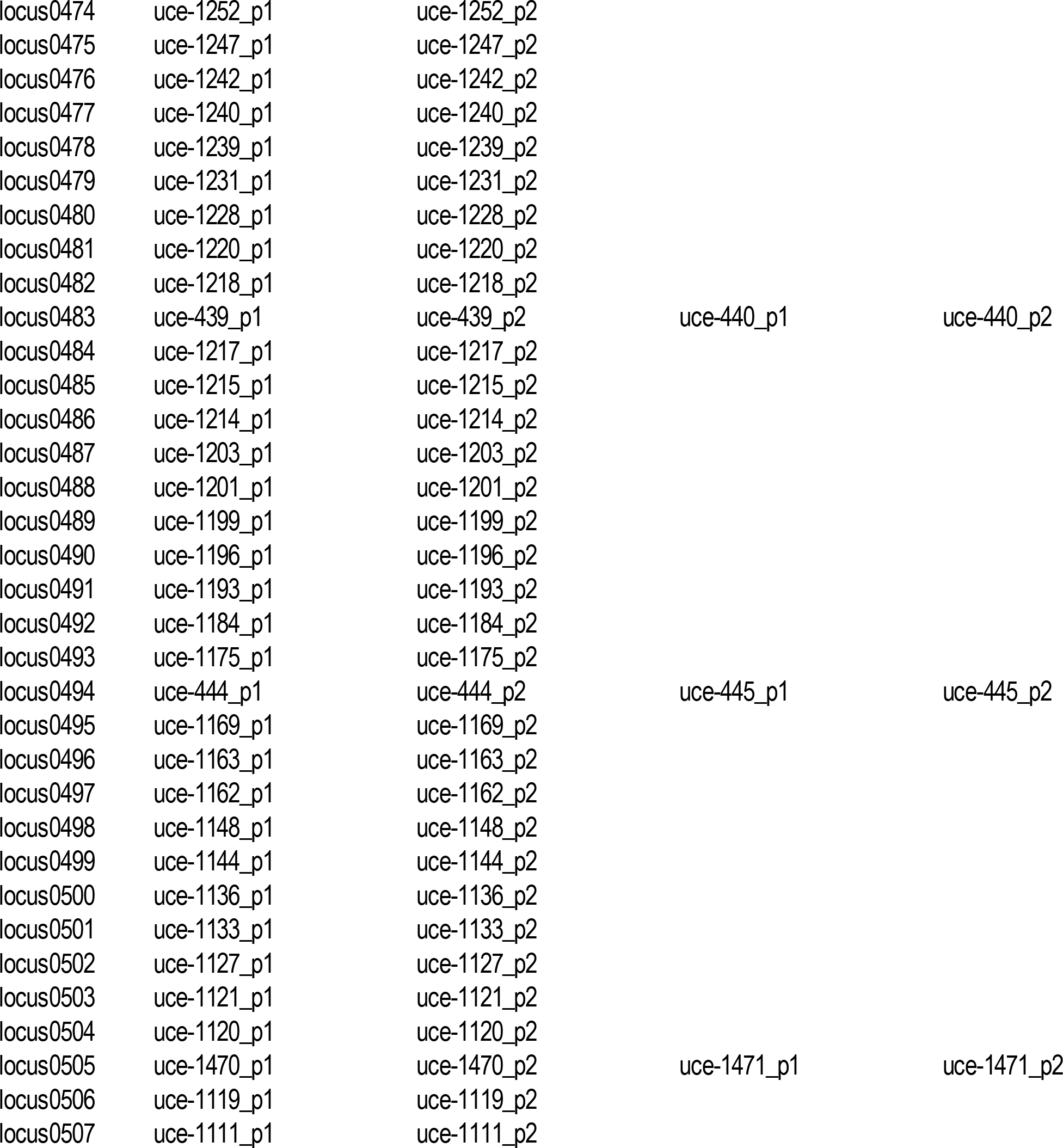

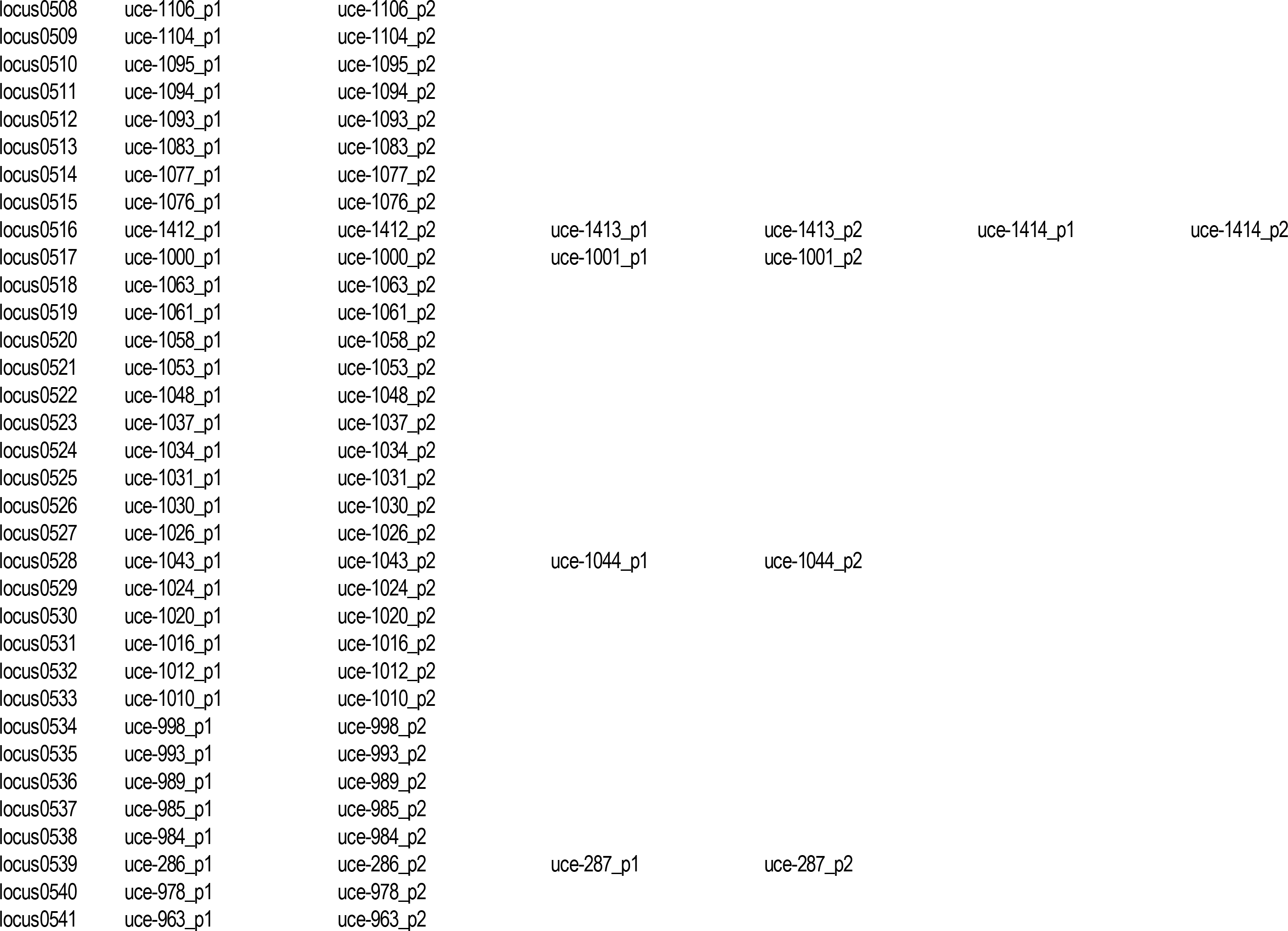

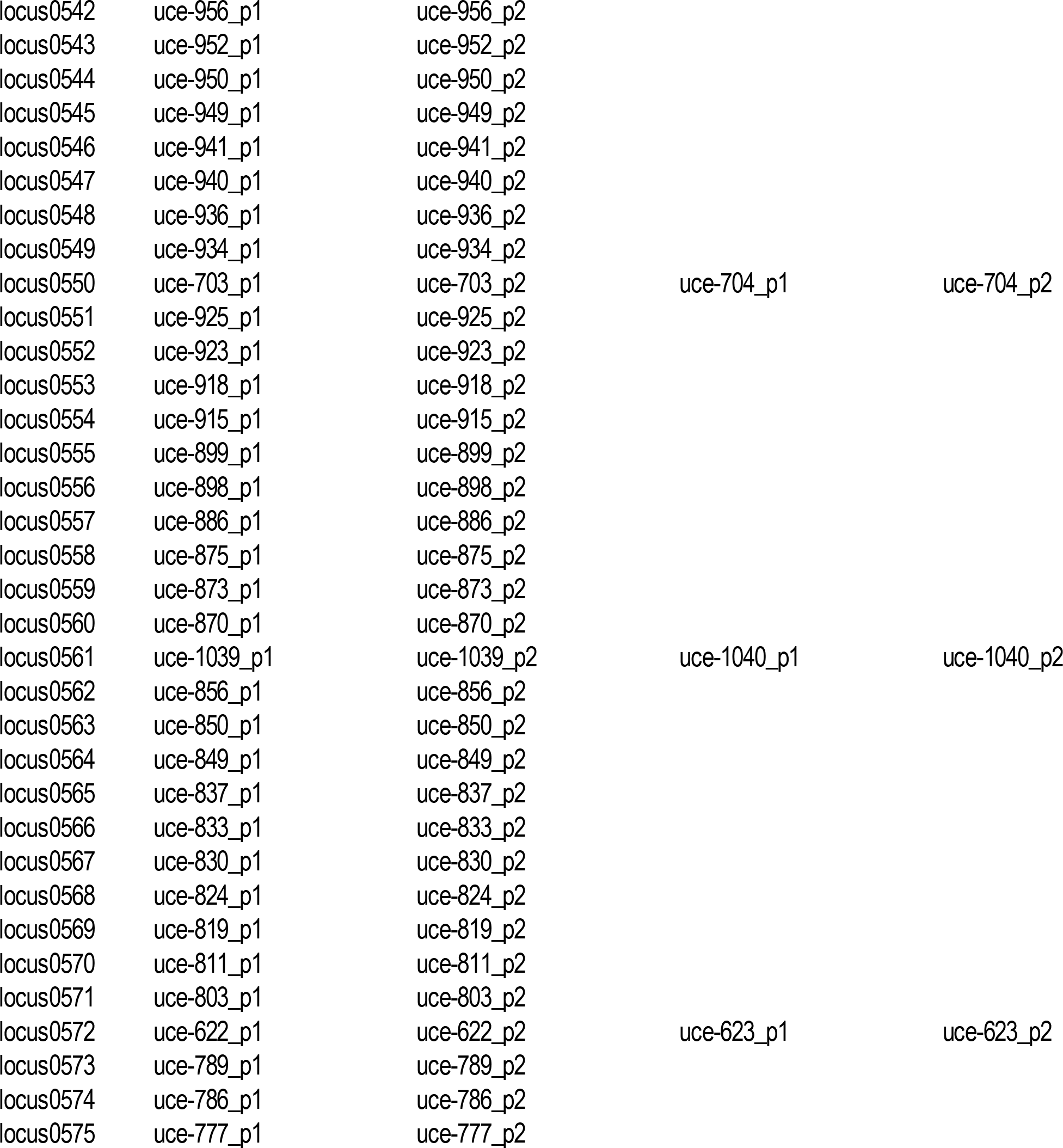

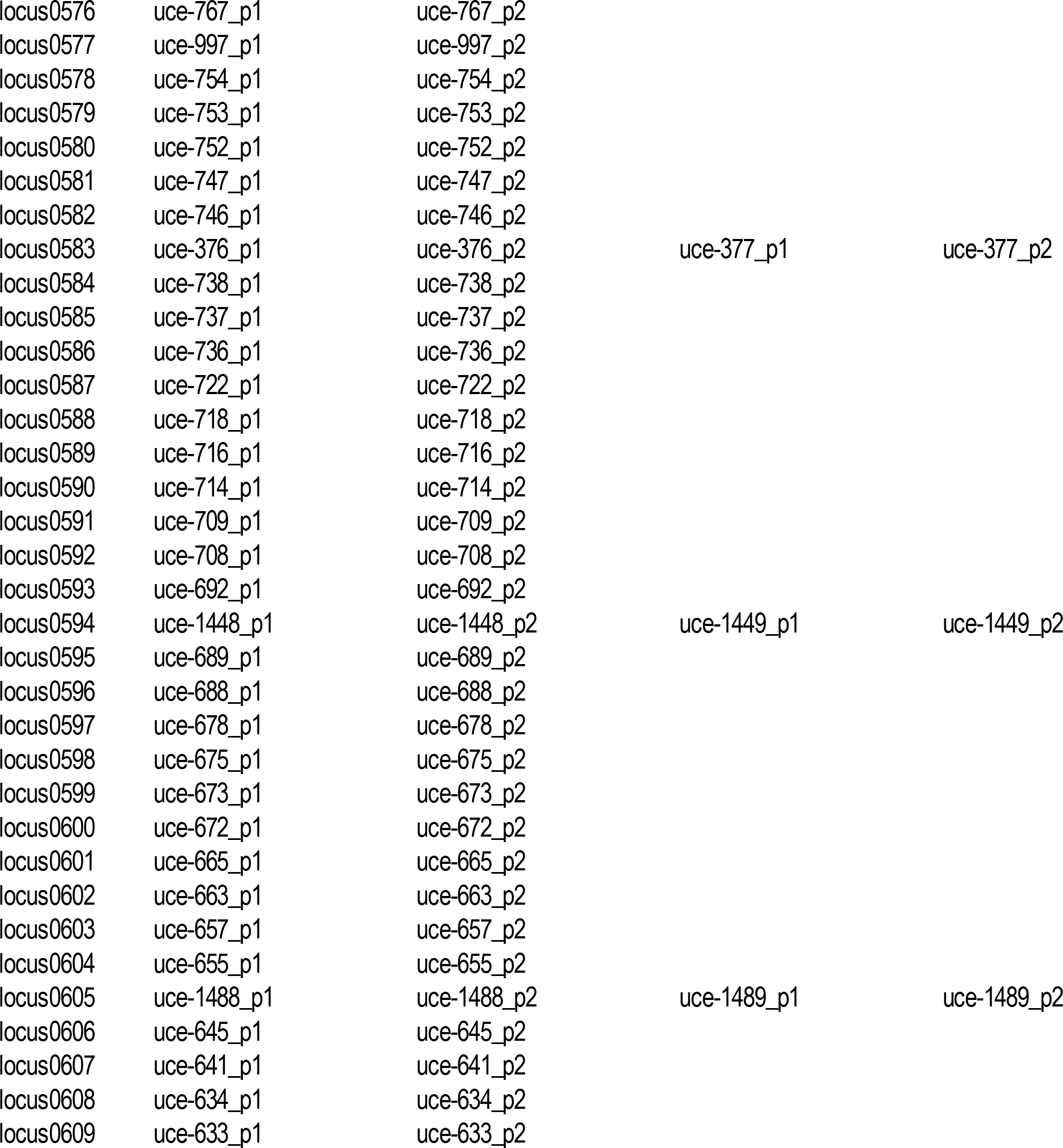

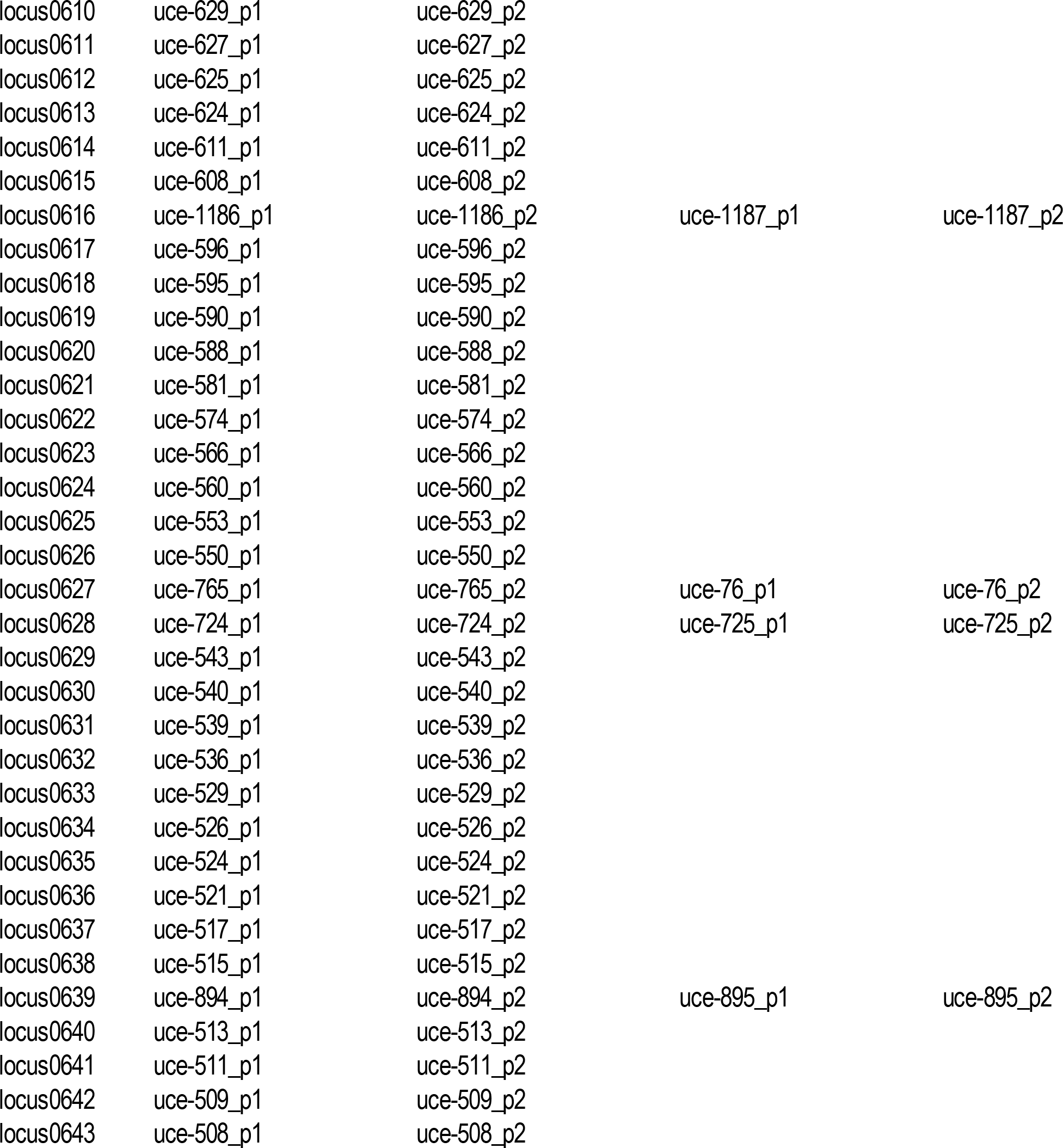

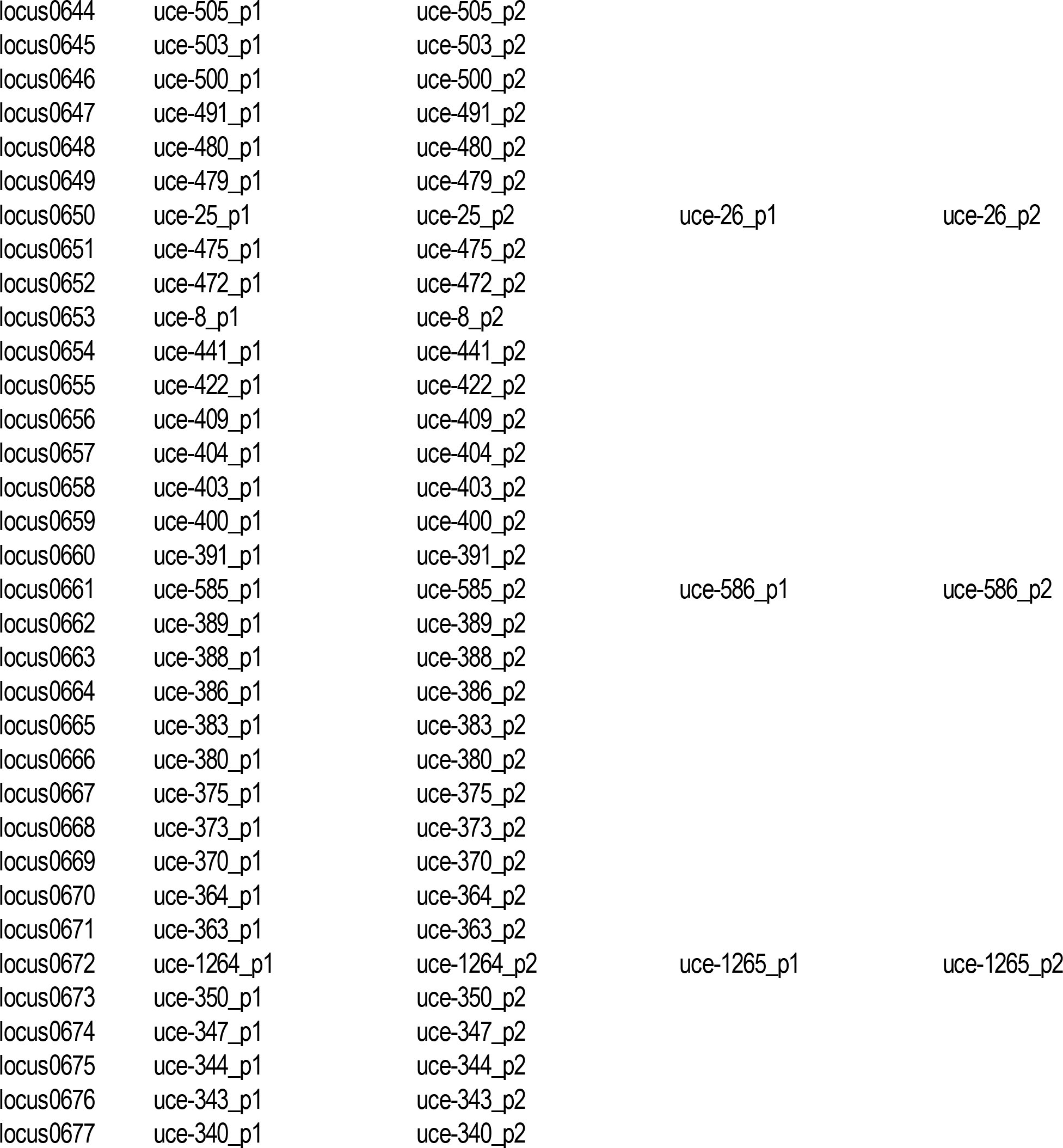

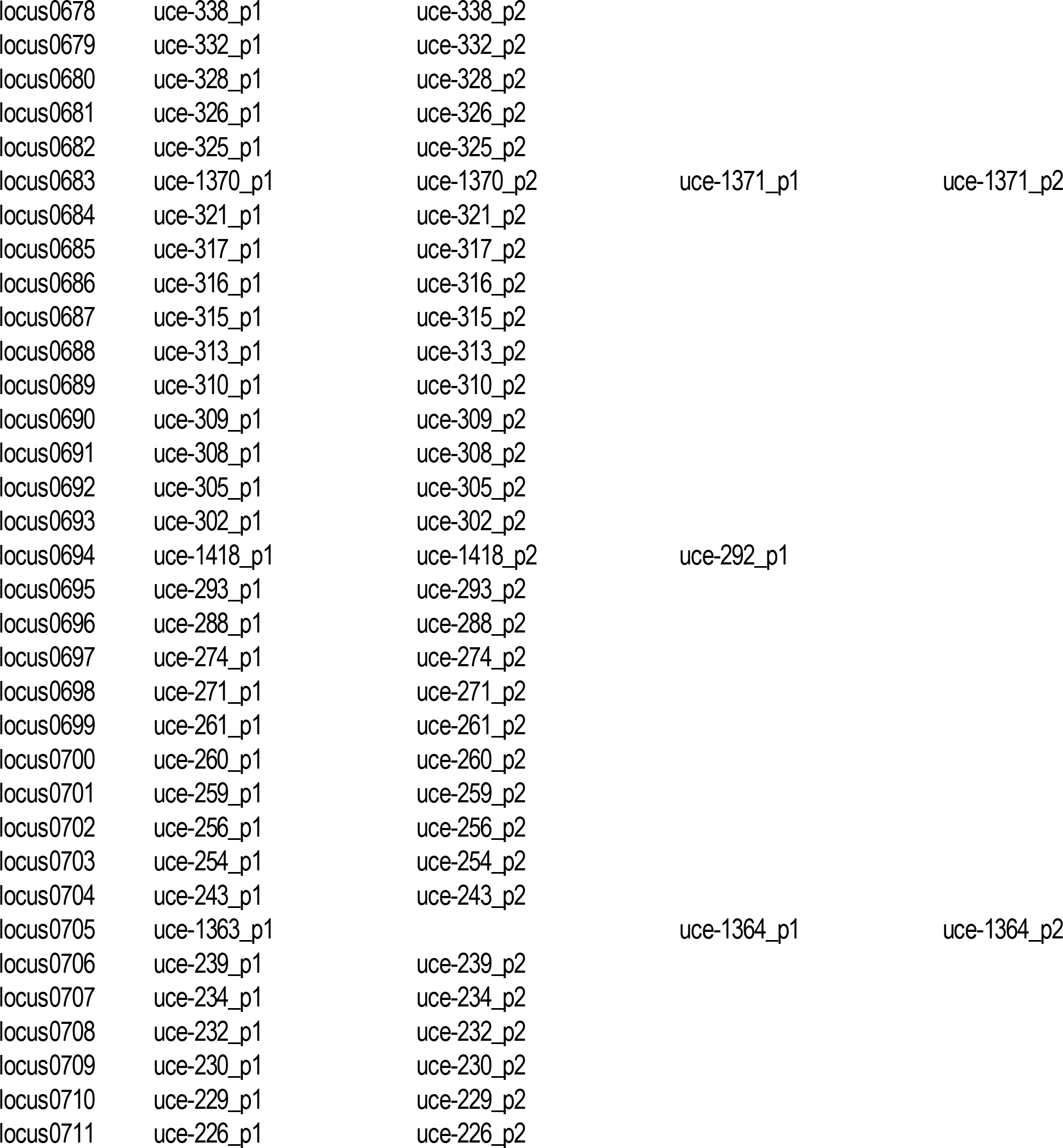

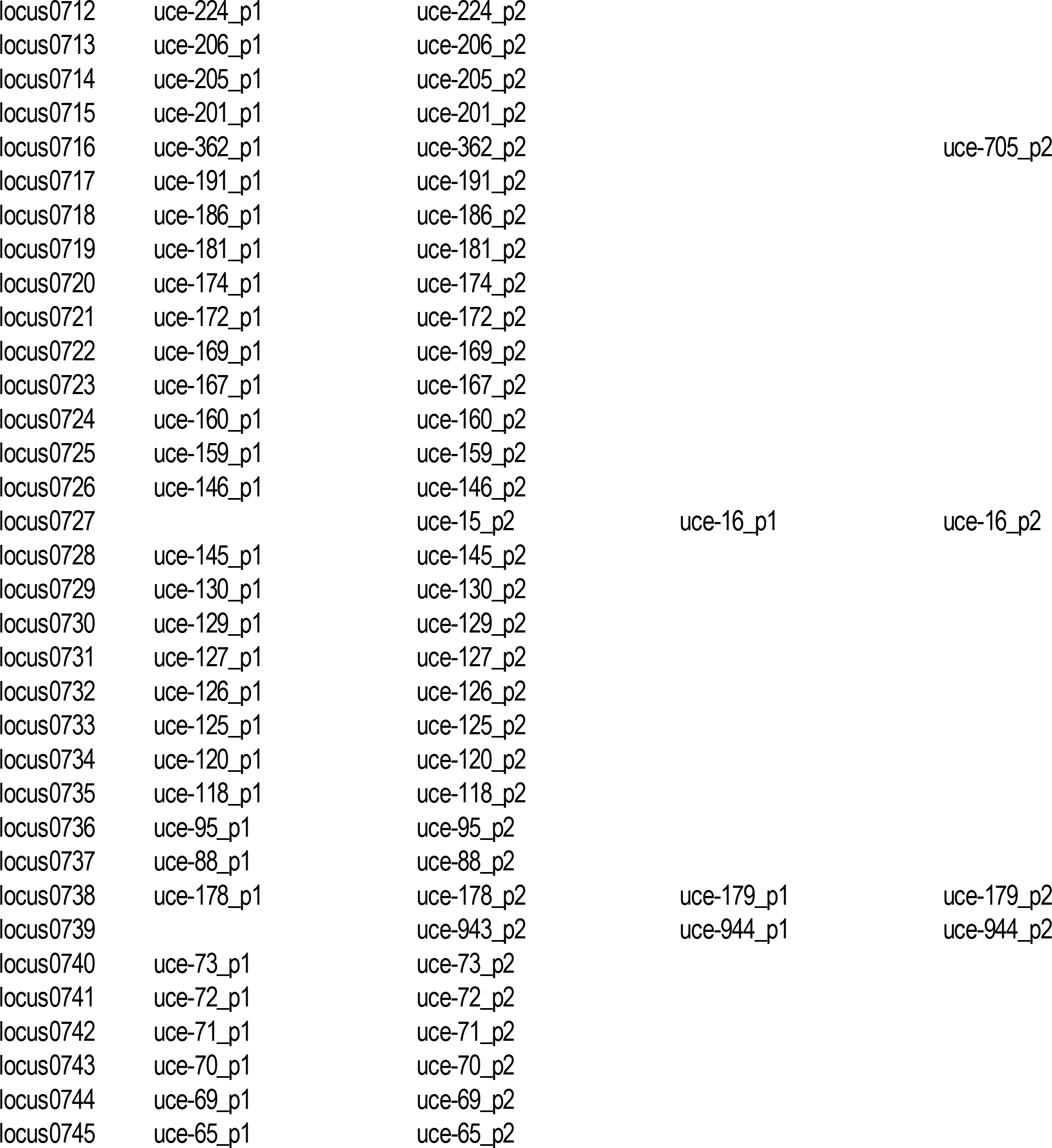

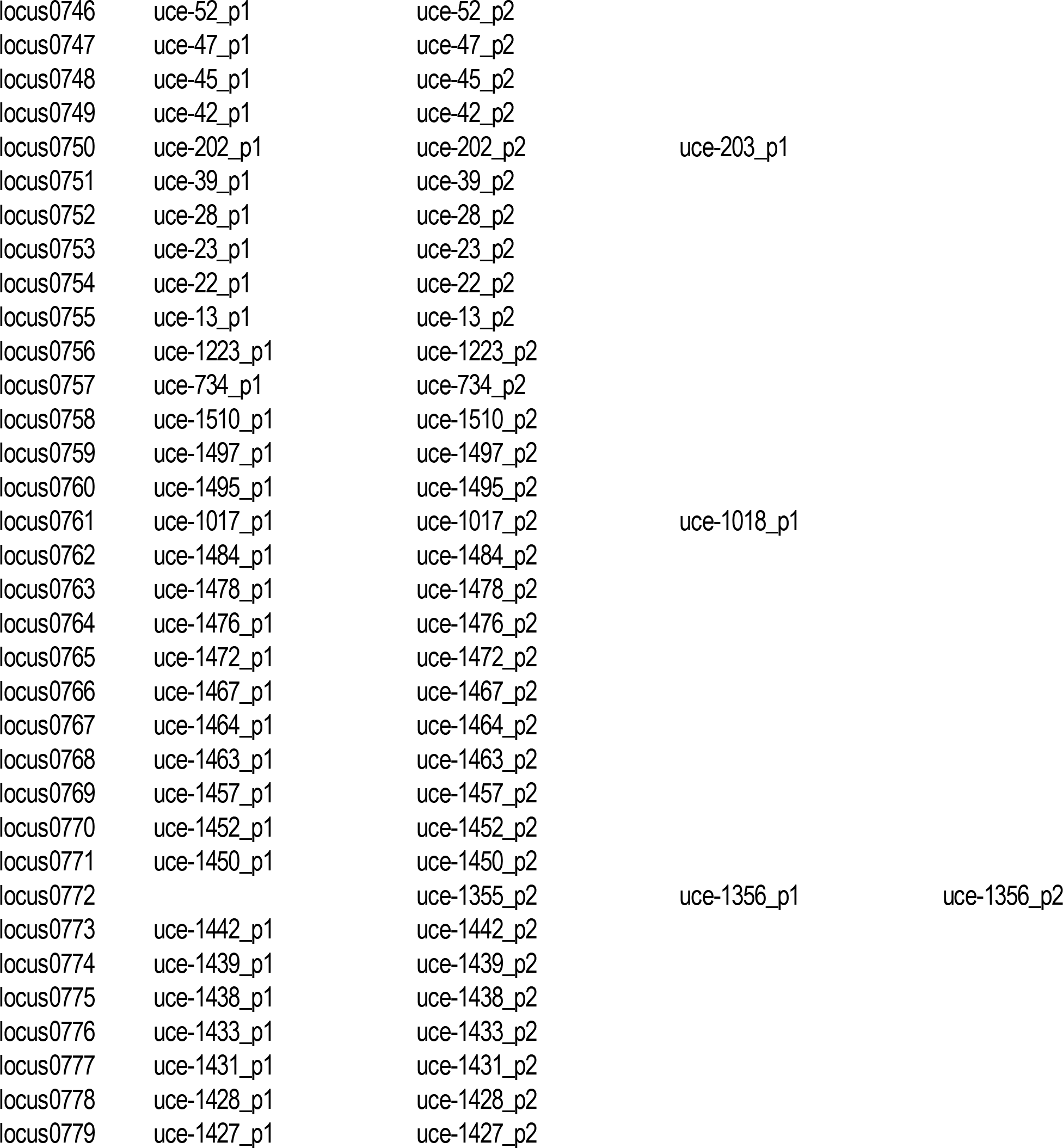

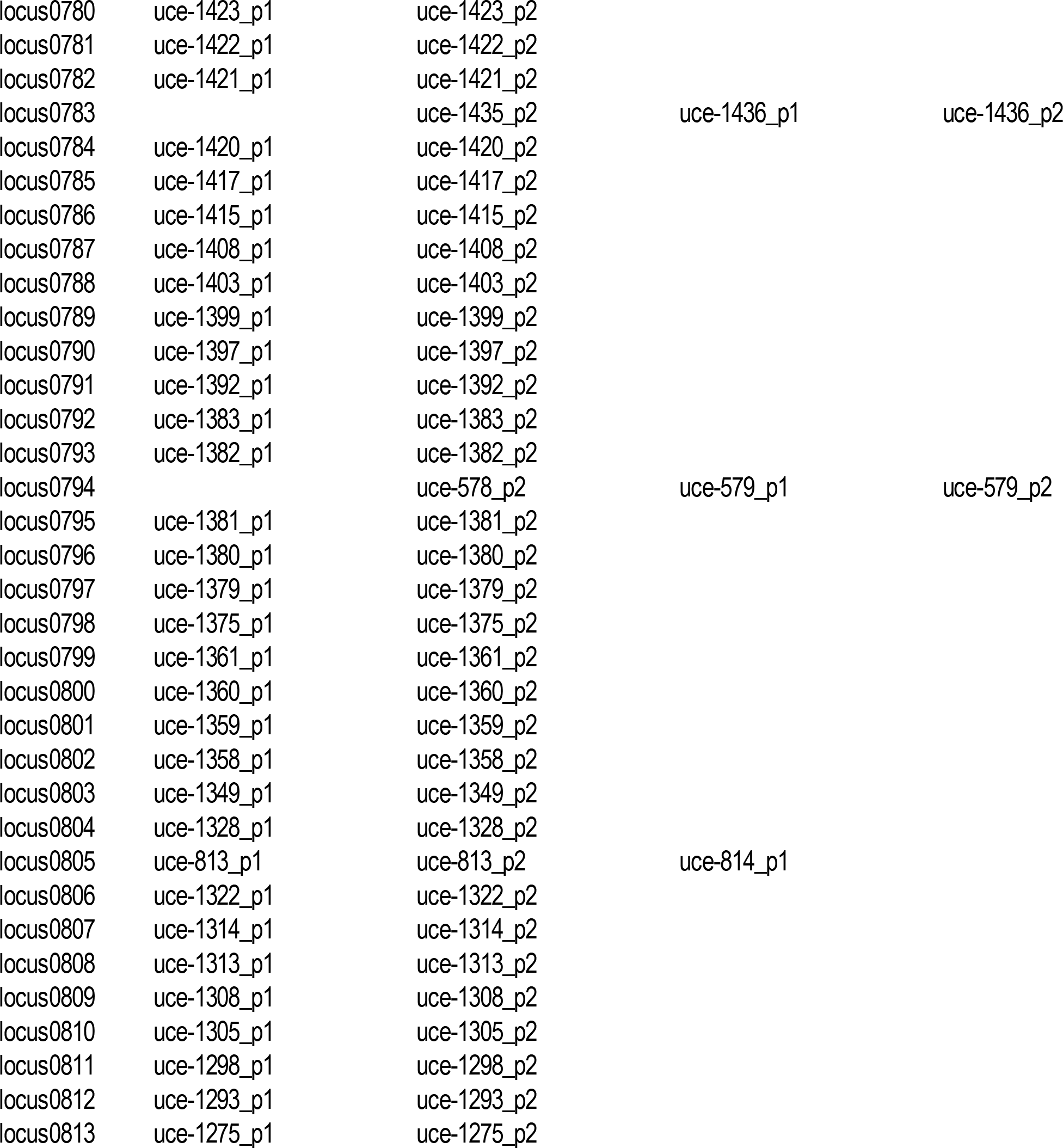

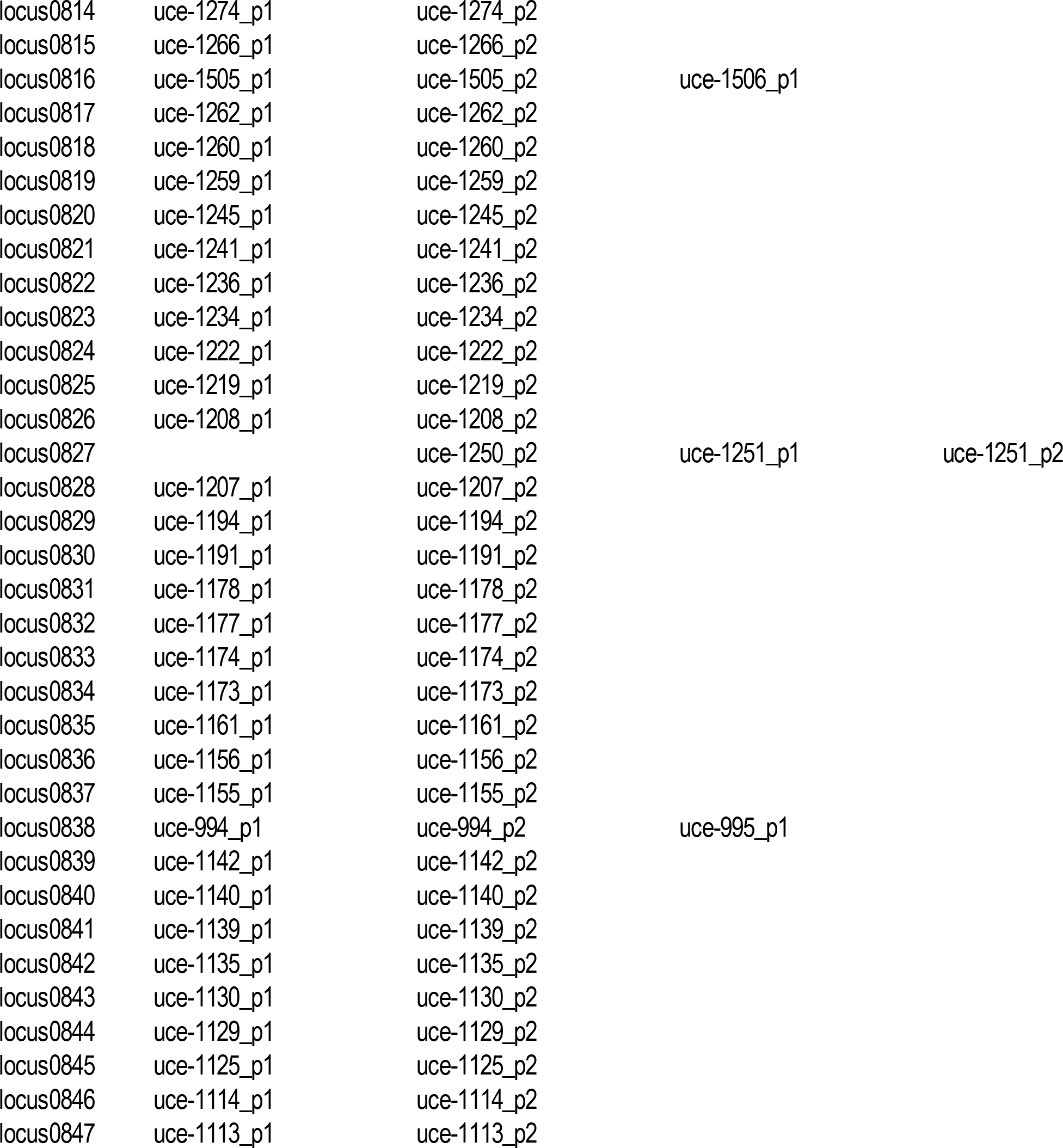

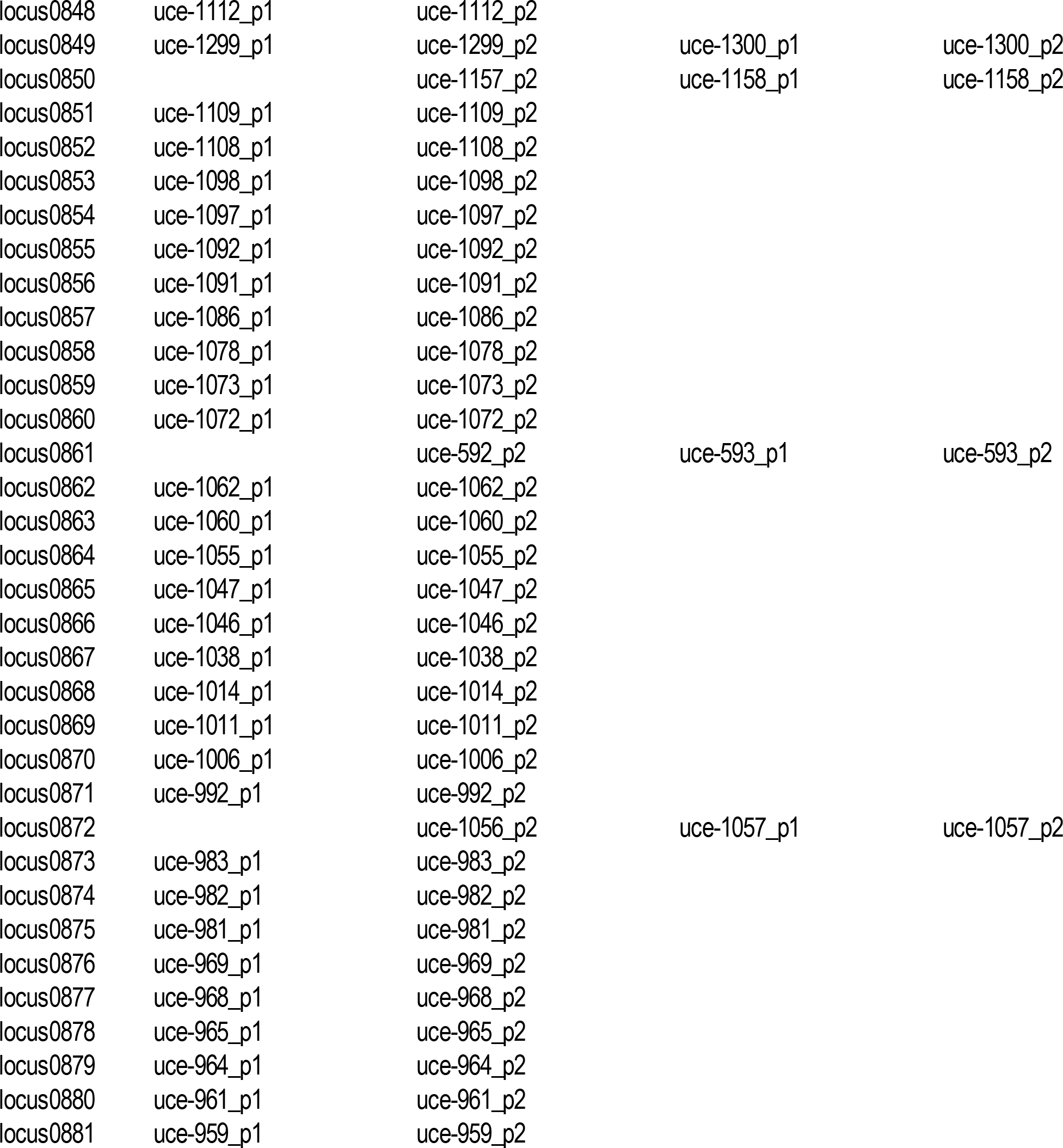

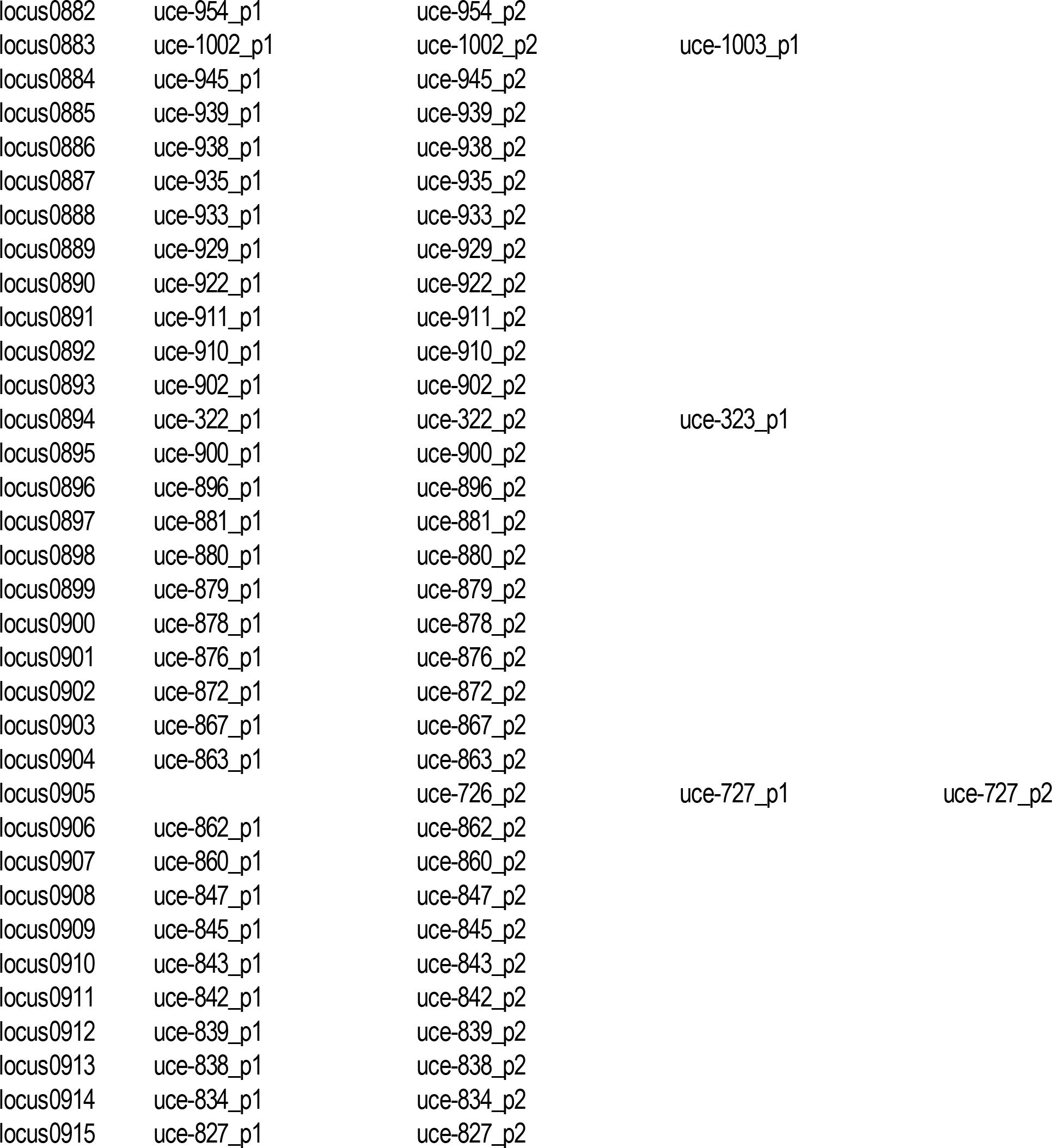

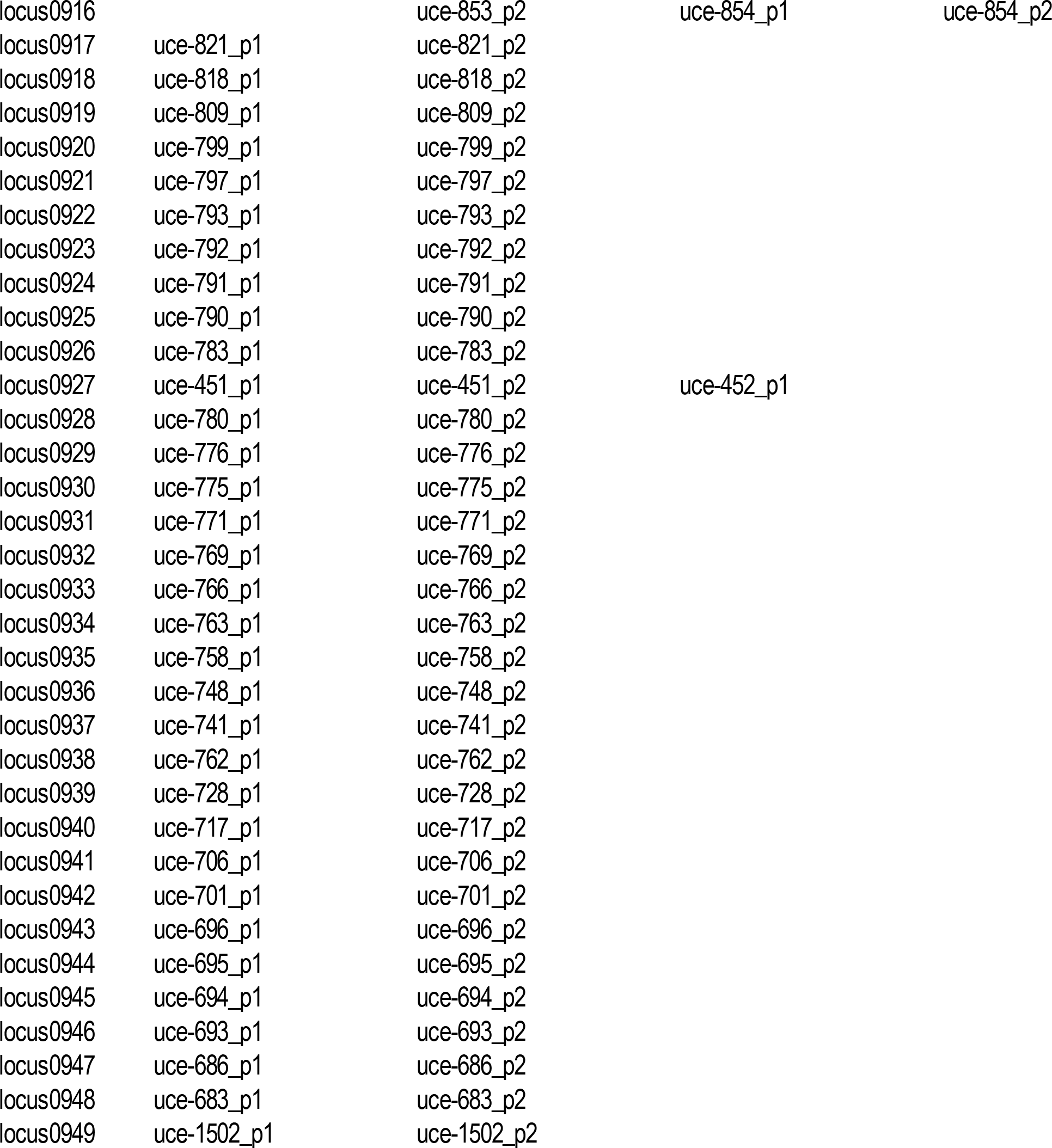

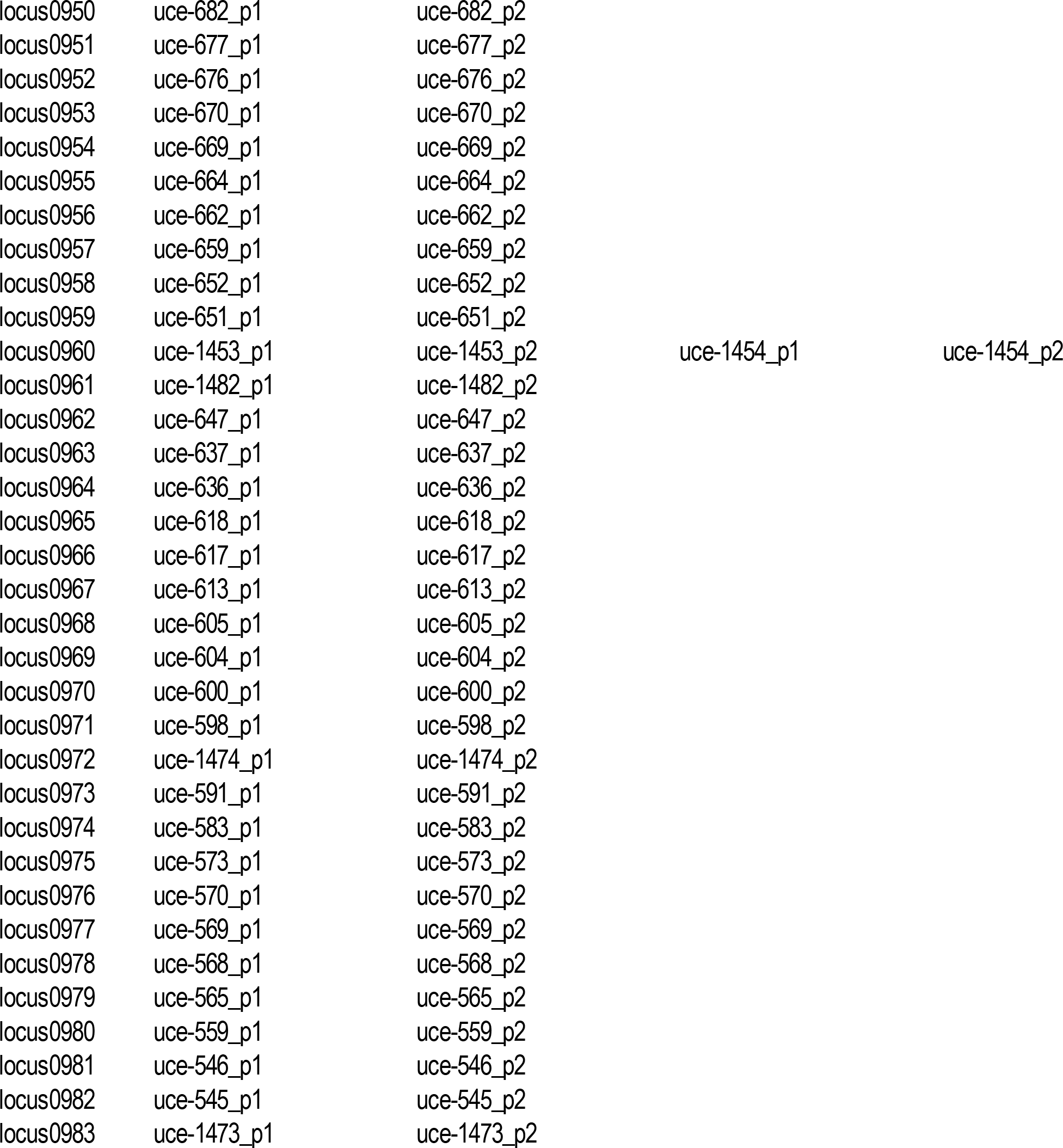

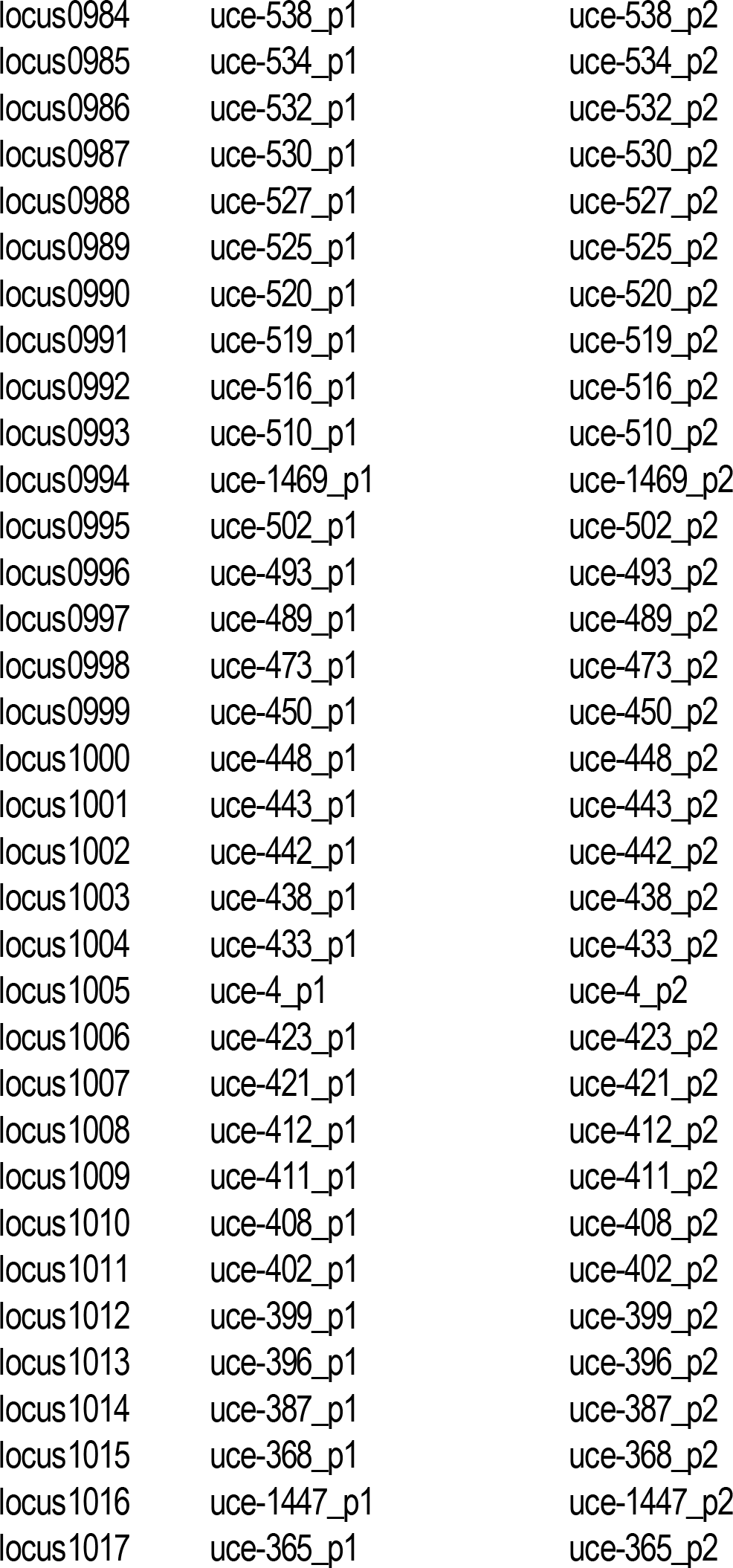

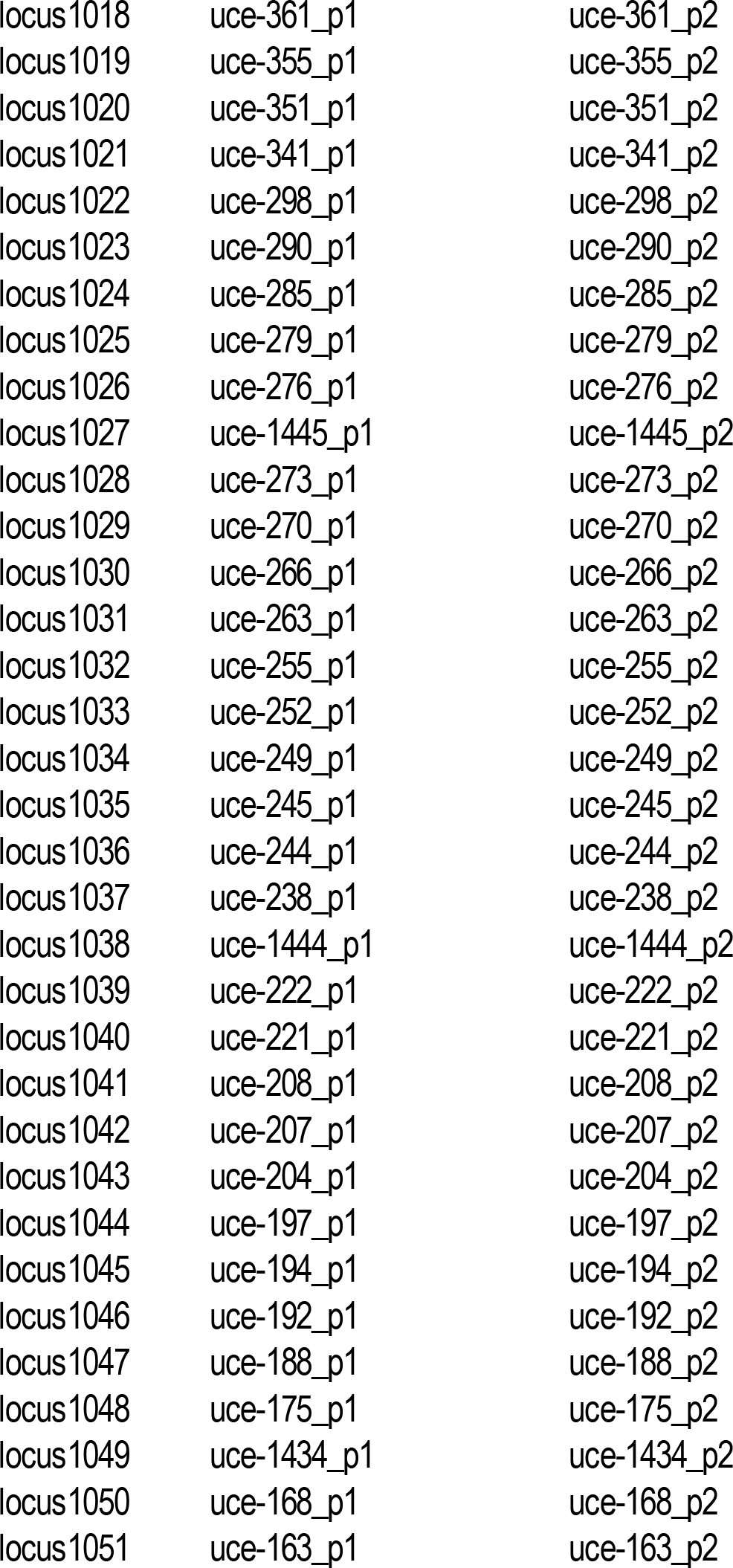

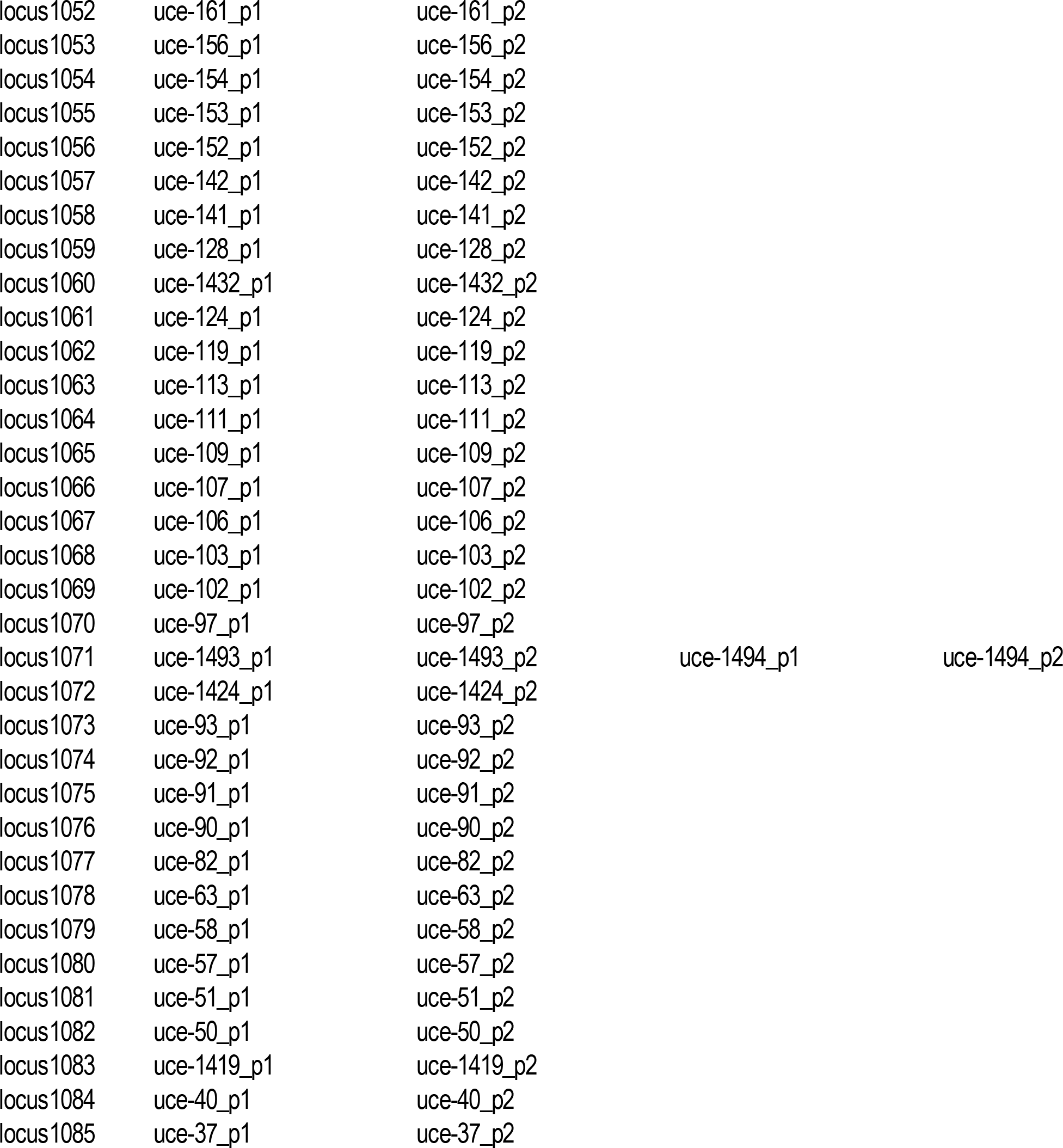

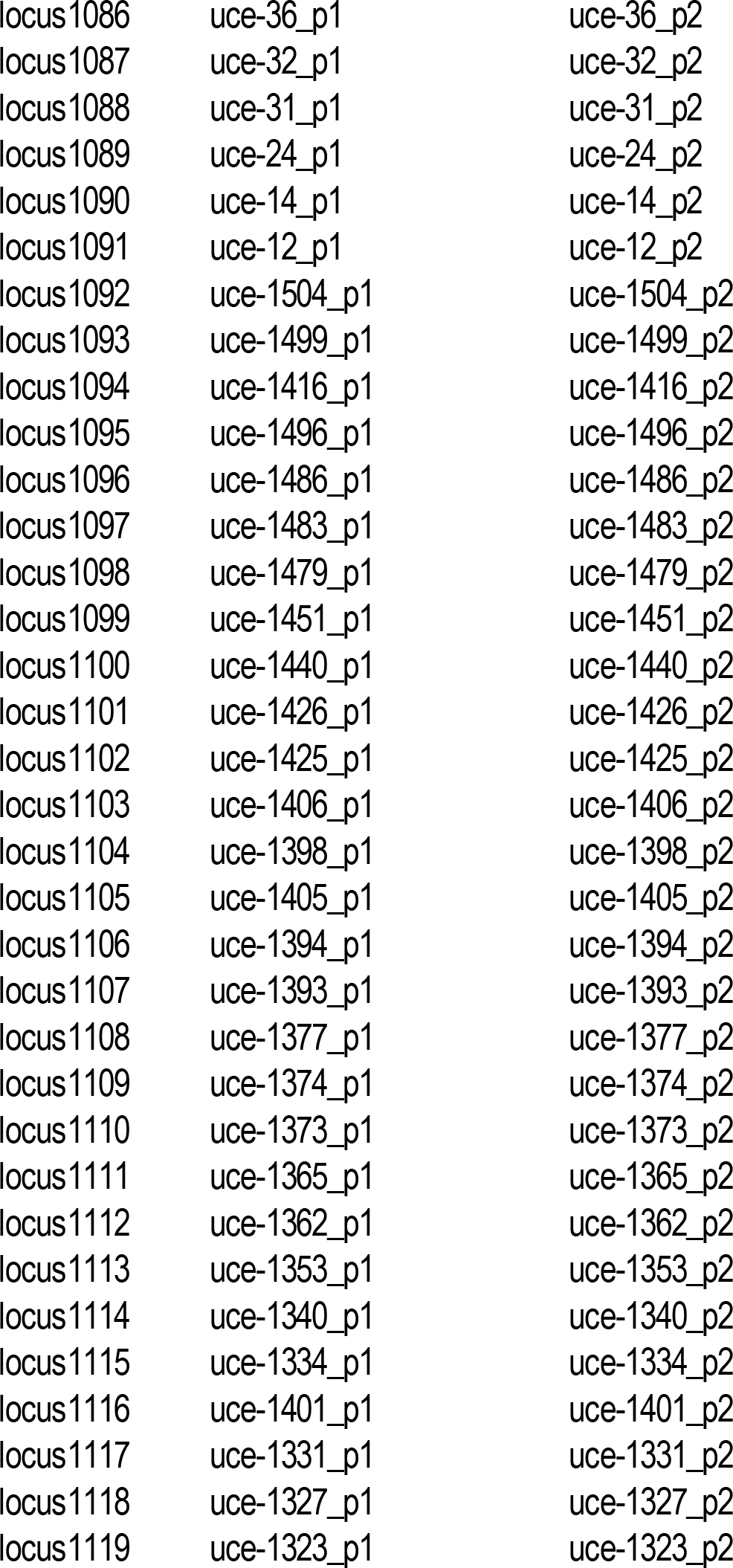

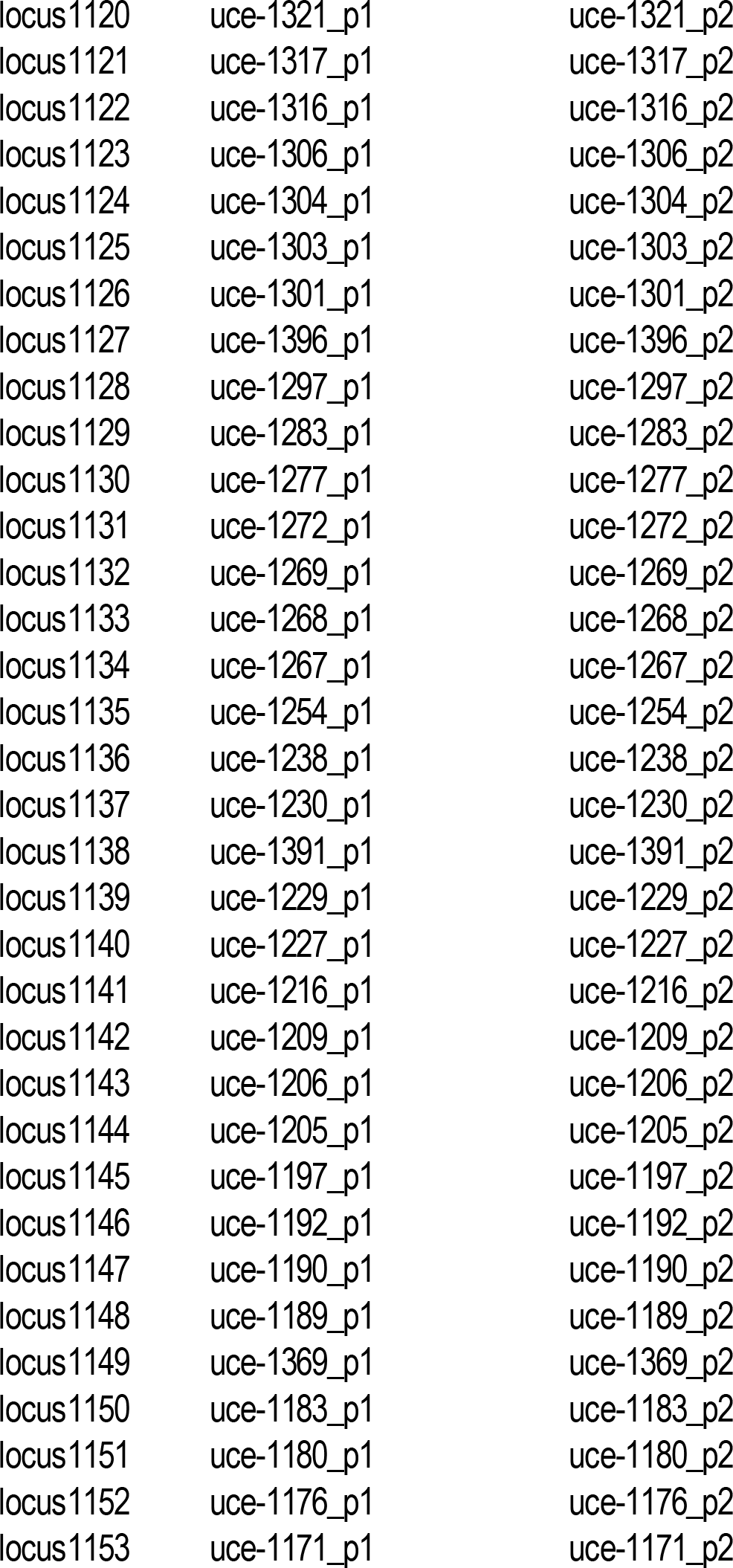

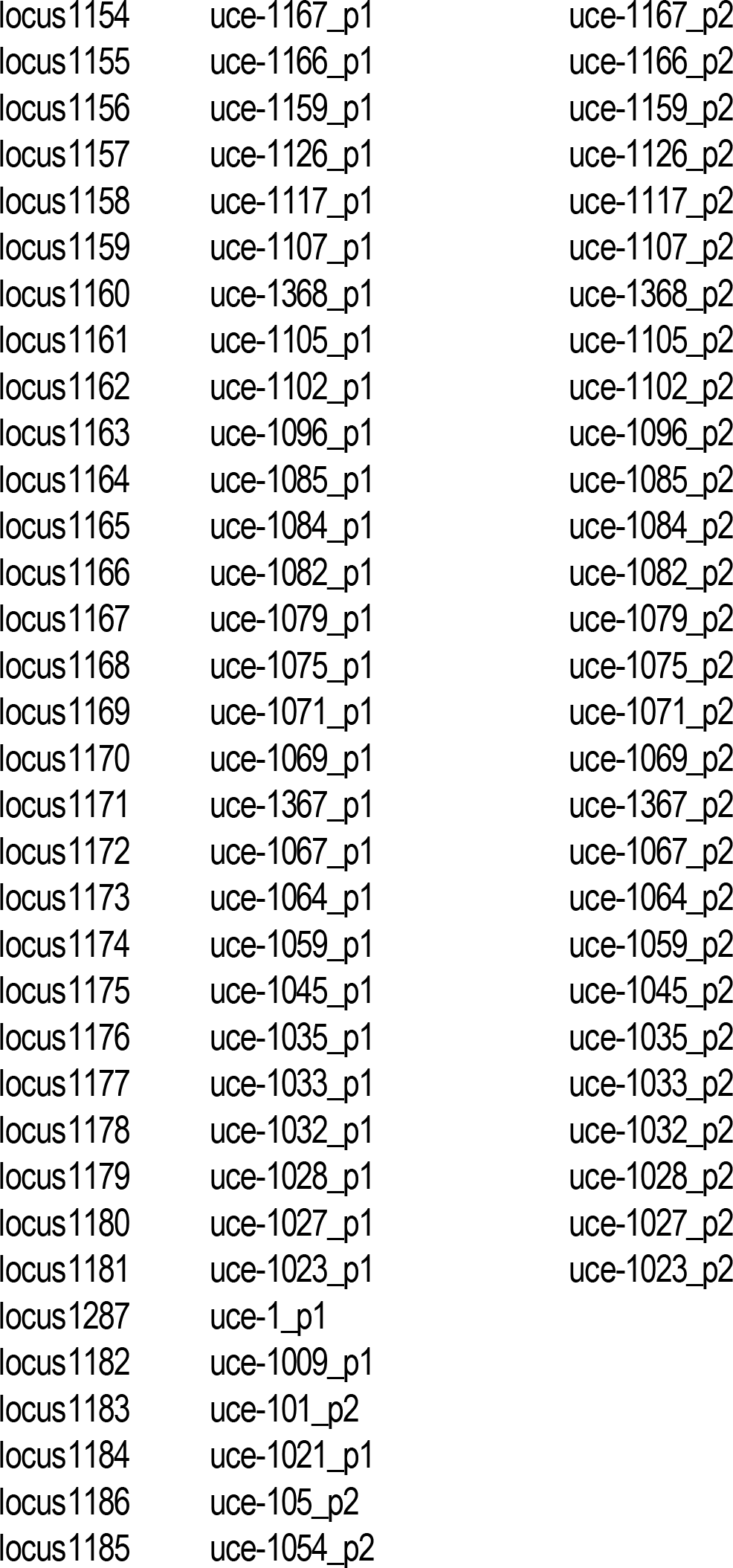

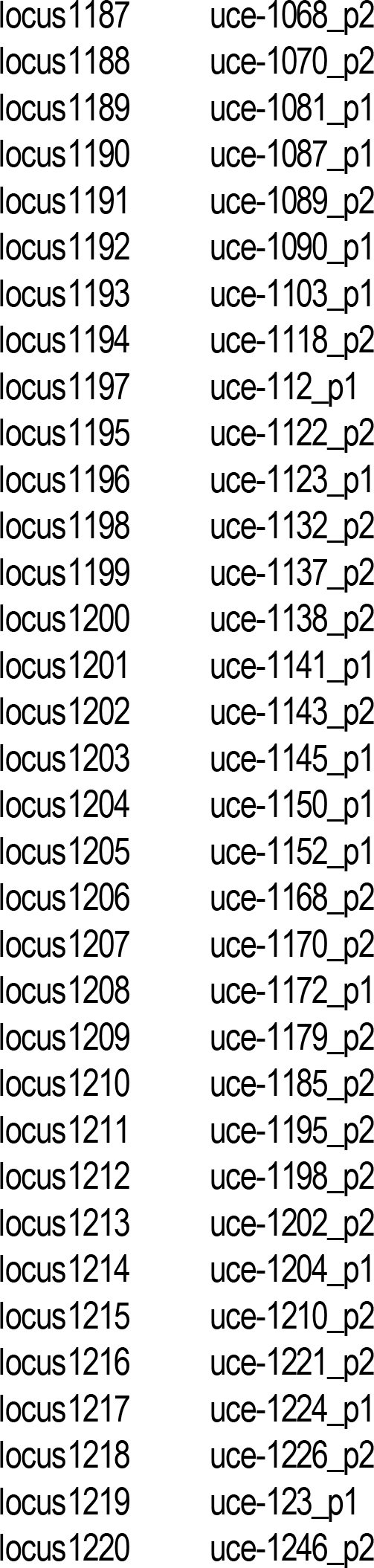

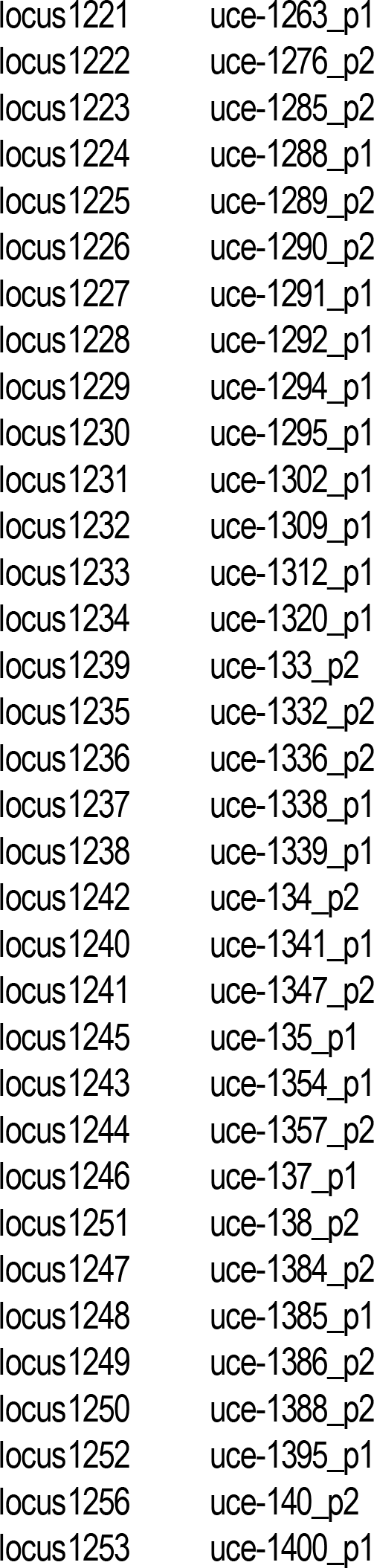

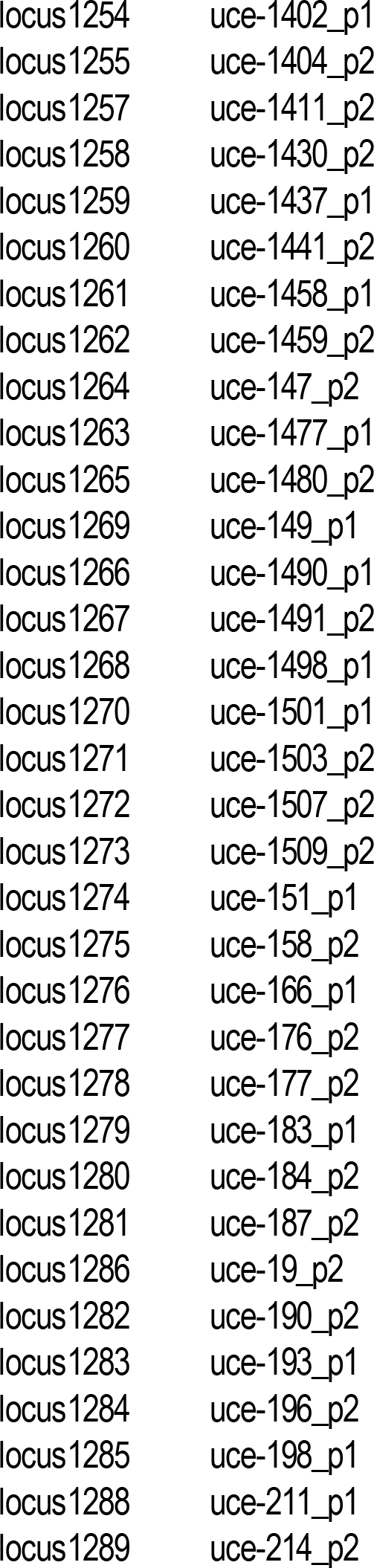

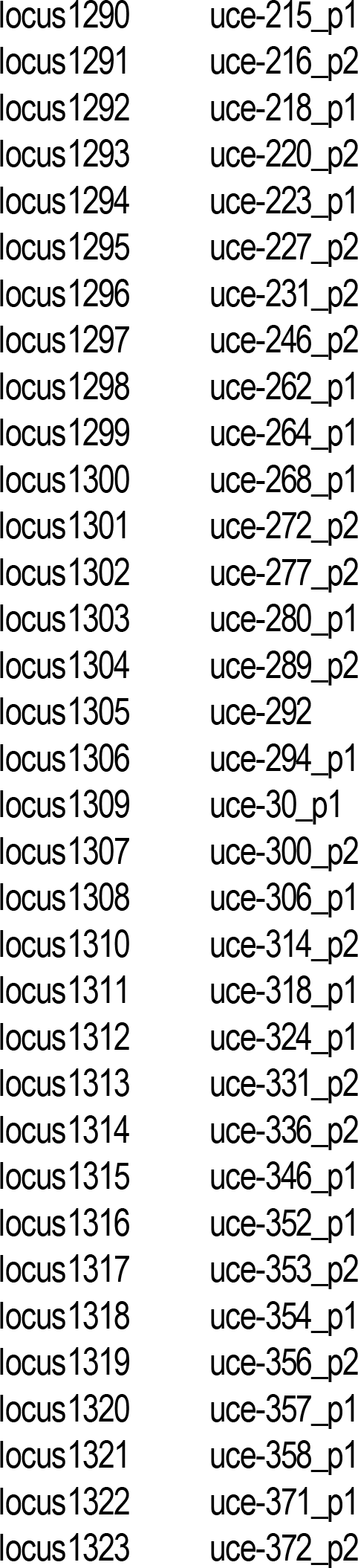

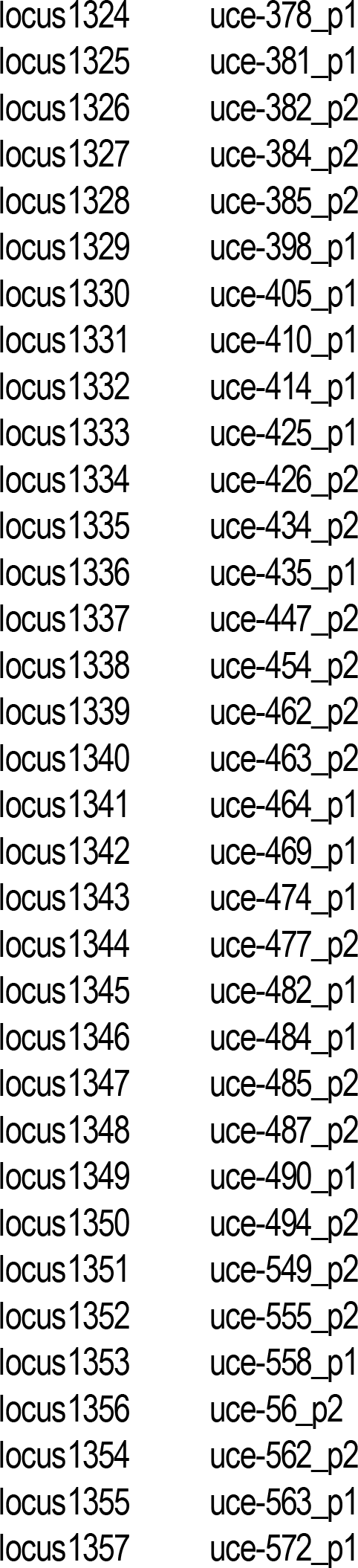

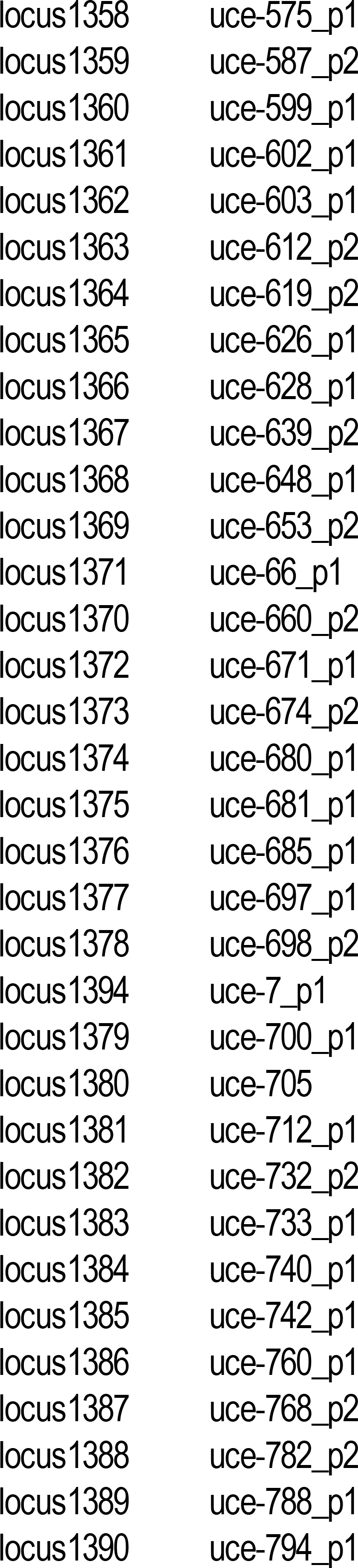

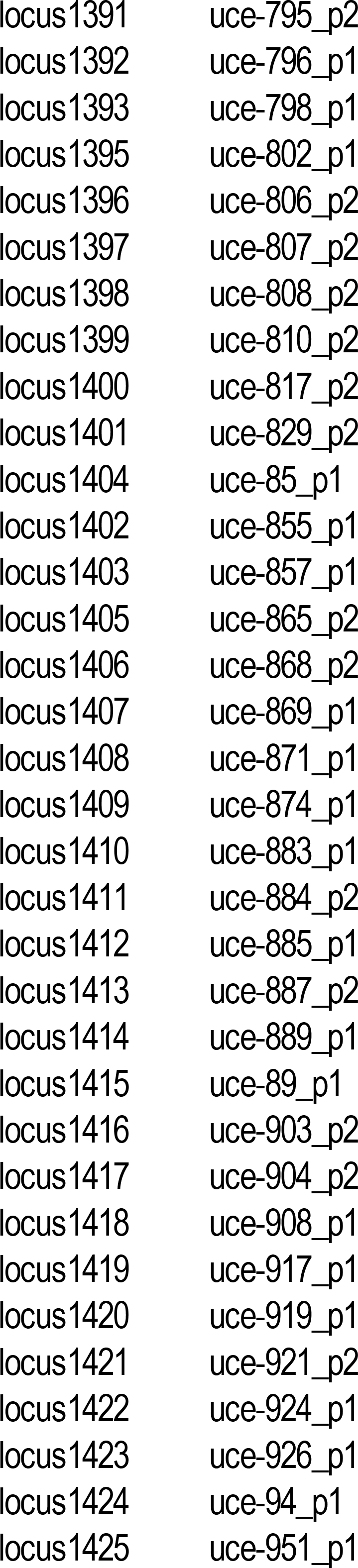

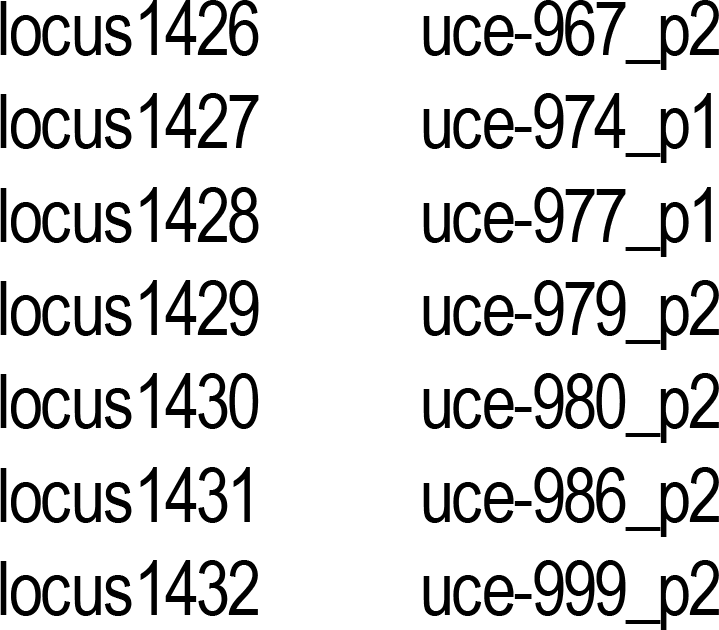

